# Effects of individual base-pairs on *in vivo* target search and destruction kinetics of small RNA

**DOI:** 10.1101/2020.07.24.215095

**Authors:** Anustup Poddar, Muhammad S. Azam, Tunc Kayikcioglu, Maksym Bobrovskyy, Jichuan Zhang, Xiangqian Ma, Piyush Labhsetwar, Jingyi Fei, Digvijay Singh, Zaida Luthey-Schulten, Carin K. Vanderpool, Taekjip Ha

## Abstract

Base-pairing interactions mediate intermolecular target recognition in many biological systems and applications, including DNA repair, CRISPR, microRNA, small RNA (sRNA) and antisense oligo therapies. Even a single base-pair mismatch can cause a substantial difference in biological activity but presently we do not yet know how the target search kinetics *in vivo* are influenced by single nucleotide level changes. Here, we used high-throughput sequencing to identify functionally relevant single point mutants of the bacterial sRNA, SgrS, and quantitative super-resolution microscopy to probe the mutational impact on the regulation of its primary target, *ptsG* mRNA. Our super-resolution imaging and analysis platform allowed us to further dissect mutational effects on SgrS lifetimes, and even subtle changes in the *in vivo* rates of target association, *k*_on_, and dissociation, *k*_off_. Mutations that disrupt binding of a chaperone protein, Hfq, and are distal to the mRNA annealing region still decreased *k*_on_ and increased *k*_off_, providing an *in vivo* demonstration that Hfq directly facilitates sRNA-mRNA annealing. Single base-pair mismatches in the annealing region reduced *k*_on_ by 24-31% and increased *k*_off_ by 14-25%, extending the time it takes to find and destroy the target mRNA by about a third, depending on whether an AU or GC base-pair is disrupted. The effects of disrupting contiguous base-pairing are much more modest than that expected from thermodynamics, suggesting that Hfq also buffers base-pair disruptions.

## Introduction

Myriad biological systems use base-pairing interactions for target recognition where proteins mediate base-pairing interactions between two physically separated strands. Such base-pairing-mediated targeting is found in a wide range of processes including DNA repair^1^, noncoding RNA-based gene regulation^2,3^, bacterial immunity using CRISPR^4^, and therapies using anti-sense oligonucleotides^5^. They all rely on base-pairing interactions above a threshold for specificity. How do they achieve both accuracy and speed to sample through thousands of potential targets and rapidly reject non-targets? Recent advances in single-molecule imaging technologies made it possible to explore the kinetic parameters of target recognition and non-target rejection *in vitro*^6–12^, and in a limited number of cases, inside living cells^13^. However, we do not yet know the impact of single nucleotide changes in *in vivo* target search kinetics, even though such minute changes can have large functional consequences. Our goal here is to quantify the mutational impact on base-pairing-mediated target search kinetics *in vivo*. We used bacterial gene regulation by small RNA (sRNA) as a model system.

Among the many examples of non-coding RNA-based gene regulation are microRNAs and long non-coding RNAs in eukaryotes and sRNAs in bacteria and archaea^14,3,15^. Often, bacterial sRNAs regulate gene expression at a post-transcriptional level during stress, for example, in iron limitation stress^16^, osmotic and acid stress^17^, and oxidative stress^18^. Our work here studied the sRNA SgrS, which is produced in response to glucose-phosphate stress.^19^

A disparity between sugar uptake and its metabolism gives rise to stress; a faster uptake leads to an accumulation of glucose-6-phosphate and activation of SgrR, a transcription factor. This stimulates the *sgrS* gene to transcribe SgrS, which reduces sugar transport, promotes efflux and reroutes cellular metabolism^20–22^. Sugar stress conditions are provoked in most studies by subjecting cells to α-methylglucoside (αMG), a sugar analogue that gets phosphorylated during import to form αMG-6-phosphate, which cannot be further processed metabolically. *E. coli* SgrS, a 227-nt sRNA, binds reversibly and dynamically to its primary target, *ptsG* mRNA^23^, which codes for the EIICB domain of the glucose phosphotransferase system. Binding between the RNAs, aided by a hexameric RNA chaperone protein Hfq, blocks the *ptsG* ribosome binding site, thereby inhibiting translation of new glucose transporters (Fig. 1a). This sRNA-mRNA complex also gets degraded by endoribonuclease RNase E, thus reducing the cellular concentration of *ptsG* mRNA. Hfq is important for the stability of sRNAs in general and *in vitro* studies have shown that Hfq increases the rate of annealing between sRNA and its target mRNA sequences^24–26^. Whether Hfq also directly facilitates annealing between sRNA and mRNA *in vivo* is unknown for any sRNA because it has not been possible to separate the effects of Hfq on sRNA stability and sRNA-mRNA annealing.

**Figure 1.**
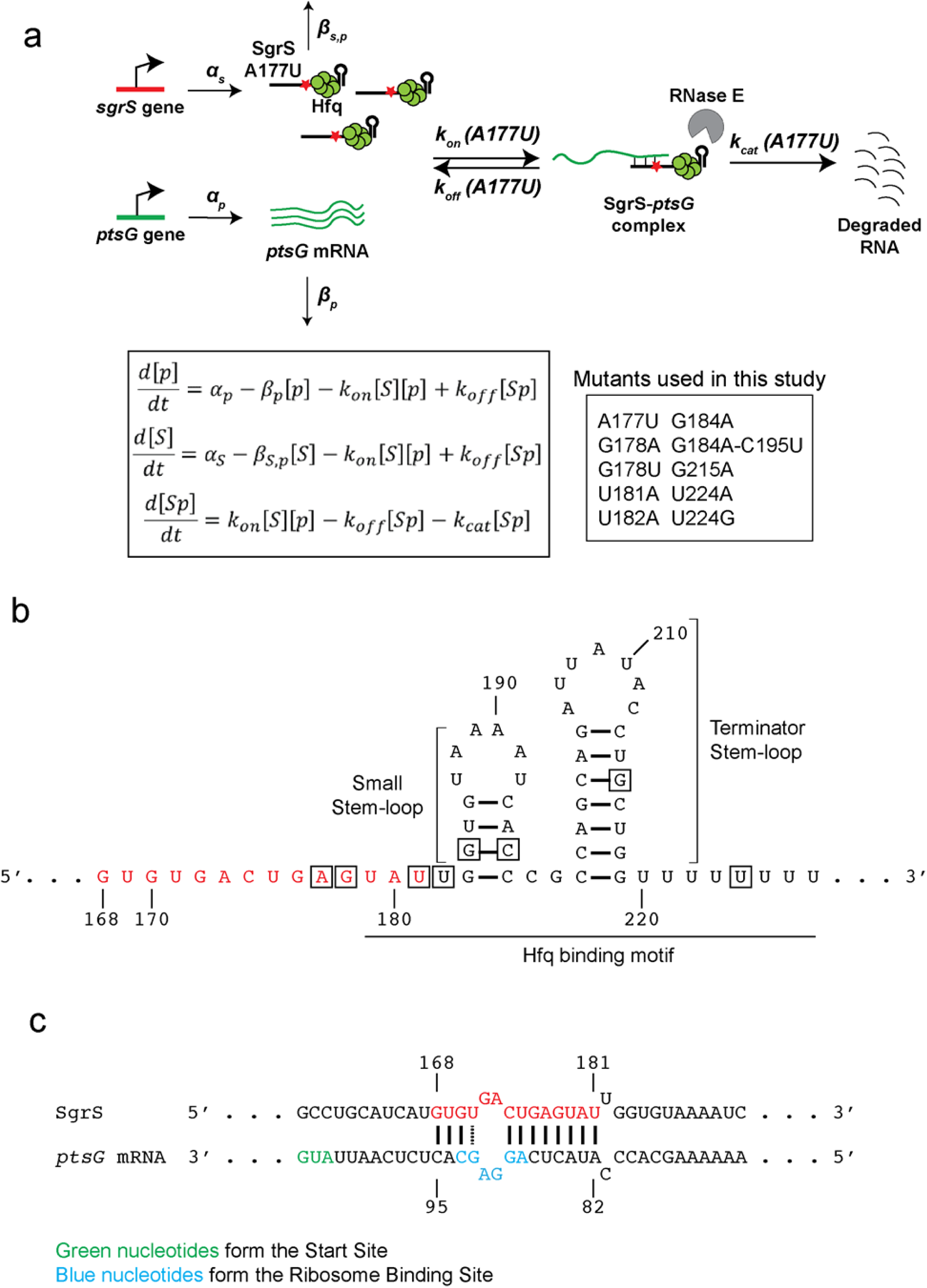
Target search kinetics of SgrS. **(a)** Kinetic scheme of *ptsG* mRNA degradation induced by wild-type SgrS sRNA and the different SgrS point mutants. The figure shows one of the SgrS mutant strains, A177U. The steps are described in detail in the main text and the inset shows the mutants used in this study. [*p*], [*S*] and [*Sp*] are the concentrations of *ptsG* mRNA, SgrS and the SgrS-*ptsG* complex, respectively, in their mass-action equations. **(b)** Secondary structure of SgrS sRNA from nucleotide 168 to the poly-U tail. The nucleotide positions where mutations were made are boxed. The nucleotides involved in base pairing with *ptsG* mRNA are red. **(c)** Base-pairing interaction between SgrS and *ptsG* mRNA showing the complementary region, start site and ribosome binding site.

SgrS contains a 3’ Hfq-binding region predicted to contain two stem-loops, the small stem-loop and the terminator stem-loop that is larger, followed by a U-rich tail (Fig. 1b)^27,28^. An optimal length of U-rich tail, with seven nucleotides or more^27,29^, is required for the formation of functional sRNAs and for efficient Hfq binding, and Hfq binding to the two stem-loops is critical for target regulation^27,28,30–32^.

Nucleotides 168-187 of SgrS are partially complementary to the *ptsG* 5’-UTR (Fig. 1c).^20^ Nucleotides 168-181, if presented as a 14 nt long oligonucleotide alone, are sufficient for full repression of *ptsG* translation *in vitro* and *in vivo*^33^. Among these, G176 and G178 have been shown to be most important for the annealing between SgrS and *ptsG* mRNA^24^.

Previously, we developed a two-color 3D super-resolution imaging and modeling platform to determine *in vivo* target search kinetics for wild-type SgrS regulation of *ptsG*^34^. The bimolecular association rate constant *k*_on_ between the RNAs was 2×10^5^ M^-1^s^-1^, which is within the wide range of reported Hfq-mediated sRNA and target mRNA association rates *in vitro* despite the crowded cellular environment and large excess of non–target RNA molecules. The dissociation rate constant *k*_off_ was 0.2 s^-1^; 10 to 100-fold larger than *in vitro* estimates of other sRNA-mRNA pairs^32,35,36^. Its non-zero value showed that even for the correct target, binding is reversible. The large dissociation constant *K*_D_ (=*k*_off_/k_on_) of ∼1 μM explained why more than a hundred SgrS molecules are needed for rapid *ptsG* mRNA regulation. The rate constant for co-degradation, *k*_cat_, was surprisingly high, 0.4 s^-1^, suggesting that RNA degradation machineries accompany the target search complex formed between SgrS and Hfq so that as soon as RNAs bind each other, RNAs can be degraded without waiting for the arrival of downstream degradation machineries. Here, by expanding the scale of this quantitative imaging-based investigation by an order of magnitude to include 10 SgrS mutants, we aimed to determine how *k*_on_, *k*_off_ and *k*_cat_ are affected by single nucleotide changes.

We formulated a pipeline of experiments to identify and examine the key regions in SgrS responsible for the annealing and regulation of *ptsG*. We used Sort-Seq, a high-throughput method that can estimate the impact of different mutations on the overall activity of the fluorescence reporter system chosen^37–39^. From the Sort-Seq results, we identified the regions in the SgrS sequence important for the overall regulation and chose nine single nucleotide substitution mutants. *E. coli* strains containing these mutations or one double substitution mutation in their endogenous chromosomal copy were constructed and studied using single-molecule fluorescence *in situ* hybridization (smFISH) followed by 2-color 3D super-resolution imaging and modeling to determine *k*_on_, *k*_off_ and *k*_cat_. Our results show that the two stem-loops at the 3’ end of SgrS play important roles in the activity of the sRNA. We also provide *in vivo* evidence that Hfq directly facilitates SgrS-*ptsG* mRNA base-pairing. Importantly, we were able to unambiguously ascribe relative contributions of single base-pairs to sRNA lifetimes and target search kinetics, allowing us to quantify by how much the rates of mRNA binding and rejection are influenced by eliminating a single base-pair between them.

## Results

### Sort-Seq reveals SgrS nucleotides important for target regulation

We employed a high-throughput Sort-Seq approach to identify SgrS regions important for *ptsG* regulation. We created a low copy number reporter plasmid containing a partial *ptsG* sequence (105 nt 5’-UTR along with the first 30 nt coding sequence of *ptsG* mRNA) and superfolder GFP-coding sequence (*ptsG*-*sf*GFP)^40^ (Supplementary Fig. 1) and transformed it into *E. coli* strain MB1 (Δ*ptsG*, Δ*sgrS, lacI*^*q*^, *tetR*). The *sgrS* mutation library was constructed by random mutagenesis PCR of a plasmid^40^ containing the *sgrS* sequence (Supplementary Fig. 1) and was then transformed into the MB1 strain containing the reporter plasmid (Fig. 2a). The expression of *ptsG*-*sf*GFP and *sgrS* were under the control of P_Llac-O1_ and P_Ltet-O1_, respectively, and were induced by Isopropyl β-D-1-thiogalactopyranoside (IPTG) and anhydrotetracycline (aTc). Upon induction by IPTG, cells containing the target reporter (*ptsG*-*sf*GFP) alone showed bright fluorescence, while those co-transformed with the plasmid containing wild-type *sgrS* showed weak GFP fluorescence in the presence of both IPTG and aTc in single-cell imaging (Supplementary Fig. 2) and flow cytometry analysis (Fig. 2b), indicating an effective repression of the reporter. Cells co-transformed with the *sgrS* mutant library showed a broad distribution of GFP fluorescence indicating highly variable levels of regulation by mutants (Fig. 2b). Based on the flow cytometry results, the cells were collected in five intensity bins, and the plasmids were extracted from the cells. For each bin, the mutated *sgrS* sequence from position 149 to 227 was amplified by PCR and sequenced. Sequencing was limited to this region because the 5’ region, up to nucleotide 153 and coding for the 43 amino acid peptide SgrT, is not involved in base-pairing-dependent mRNA regulation^41,42^.

**Figure 2.**
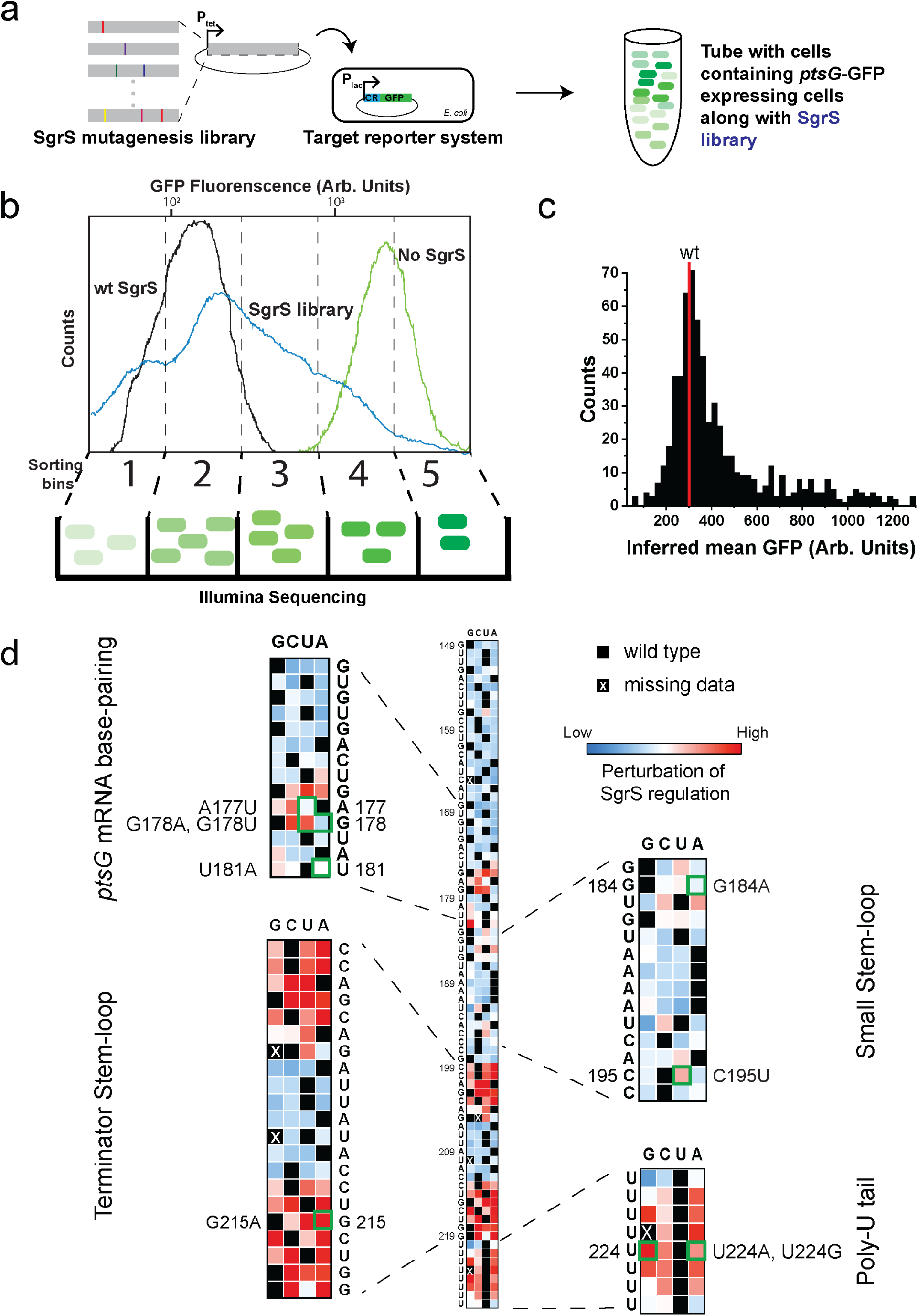
Mapping efficacy of SgrS regulation of ptsG mRNA with respect to its sequence. **(a)** Preparation of SgrS mutation library. Mutations were introduced into the SgrS plasmid using mutagenesis PCR. This library was then transformed into an *E. coli* strain already transformed with the *ptsG* mRNA plasmid fused with a GFP reporter. **(b)** Sorting of the cells and sequencing. The cells with two-plasmid co-expression were sorted using flow cytometry. The SgrS library (blue) shows GFP fluorescence that spans the region from the wild-type SgrS (black) to target (*ptsG*)-only (green). Cells were sorted into 5 evenly spaced (log scale) fluorescence bins and the occupancy percentages were 18.74%, 33.76%, 30.91%, 13.83% and 2.76% respectively. The cells from each bin were grown, DNA was purified, barcoded and sequenced using Illumina sequencing platform. **(c)** Histogram of the Sort-Seq measurements from the SgrS library from two replicates combined. The mean fluorescence for the wild-type SgrS is shown in red. **(d)** Heat map showing the effect of mutations on SgrS regulation of *ptsG* mRNA starting from nucleotide 149 to 227. The colors in the boxes are scaled from blue (low) to red (high), according to the level of perturbation of SgrS regulation. Black squares represent the wild-type base at each position and the black boxes with white crosses show the position of the mutants missing in the experiment. Text shows the wild type sequence of SgrS. Insets show the four regions of SgrS, viz. base-pairing region, small stem-loop, terminator stem-loop and the poly-U tail.

Using the relative abundance of sequences in each bin and the GFP fluorescence levels from the flow cytometry analysis, we calculated, for each single point mutation, the average fluorescence intensity of cells sharing the same mutation as a measure of the regulation defect (Fig. 2d).^39^ High average single cell fluorescence would correspond to SgrS mutants that are highly defective in regulation of *ptsG* reporter expression and vice versa. The degree of perturbation to the regulatory capacity is color-coded in the heatmap grid, ranging from the least (blue) through intermediate (white) to the most (red). Nucleotides 149 to 174 showed little to no perturbation of SgrS regulation as shown by the blue squares (Fig. 2d). In contrast, the region where SgrS can base-pair with *ptsG* mRNA (U175 to G186) displayed perturbations across a wide range as shown by the white and red squares in the grid. Specifically, previous studies showed that G176C or G178C eliminates the SgrS’s ability to downregulate *ptsG* while C174G and G170C only weakly perturbs SgrS function *in vivo* and *in vitro*^24,31^. The corresponding squares in our heatmap grid (Fig. 2d) show red or dark red for G176C and G178C, and white or light blue for C174G and G170C, validating our Sort-Seq results. We also see that SgrS regulation is hampered if there are mutations in the small stem-loop region (nts 183-196 (Fig 1b)), the terminator stem-loop region (nts 199-219) and the poly-U tail (nts 220-227). The largest effect is seen in the stem region of the terminator stem-loop, C199 to G205 and C213 to G219, where we see the darkest red, highlighting the importance of this stem-loop. These stem-loop regions and the poly-U tail play a role in Hfq binding^27,28^, and our Sort-Seq analysis therefore confirms that Hfq interaction is important for SgrS function in the cell.

Based on Sort-Seq results, we picked nine single point mutations for further investigation. These include mutations in the target annealing region (A177U, G178A, G178U, U181A), U-rich region upstream of the small stem-loop (U182A), the small stem-loop (G184A), the terminator stem-loop (G215A) and the poly-U tail (U224G, U224A).

### SgrS mutation effects on regulation of *ptsG* reporter

To examine the effect of the selected SgrS mutations on *ptsG* regulation, we monitored the effect of wild-type and seven of the SgrS mutants (plasmid-encoded and expressed from an inducible promoter) on the activity of a chromosomal *ptsG’-’lacZ* translational fusion (Fig. 3a). The wild-type SgrS almost completely eliminated β-galactosidase activity whereas the mutants showed regulation defects of various degrees consistent with the Sort-Seq data. SgrS G215A, which disrupts the terminator stem-loop structure, showed the largest defect. To test if the regulatory defects can be explained by a reduction of SgrS levels, for example, due to shorter cellular lifetimes associated with impaired Hfq binding, we performed Northern blot analysis. We found that SgrS abundance is not affected for four of the mutants (A177U, G178U, G178A and G184A) and is reduced by 40-50% for mutations in the terminator stem-loop or poly-U tail (G215A, U224G and U224A) (Fig. 3c). Interestingly, the latter three mutants showed large increases in readthrough transcription, suggesting that transcription termination is defective (Fig. 3d). These observations are consistent with a previous study which showed that SgrS molecules with an extended 3*’* region do not interact with Hfq^43^. Overall, single point mutations outside the large terminator stem-loop and poly-U tail have minimal impact on SgrS abundance and their regulatory defects cannot be explained by SgrS abundance changes.

**Figure 3.**
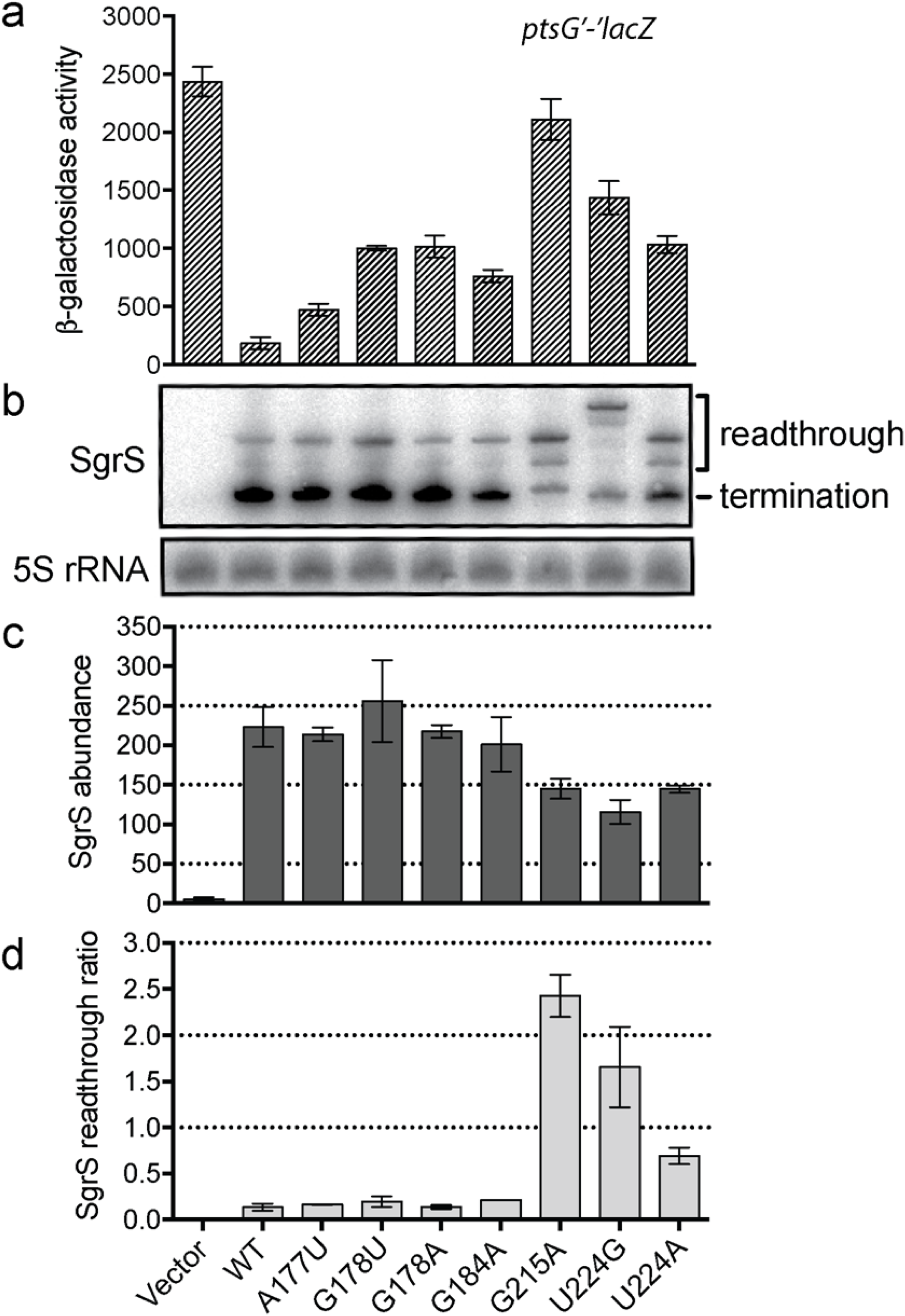
Regulation of *ptsG’-’lacZ* translational fusion by SgrS point-mutants. **(a)** Regulation of chromosomal *ptsG’-’lacZ* translational fusion by wild-type SgrS and A177U, G178U, G178A, G184A, G215A, U224A, U224G mutant variants (plasmid-encoded) was assessed using β-galactosidase activity assay. Standard error was calculated based on data from four biological replicates. **(b)** RNA was extracted simultaneously with β-galactosidase activity assay and Northern blot was performed using probes specific for SgrS sRNA and 5S rRNA (control). Full-length (227 nt), properly terminated SgrS transcripts are labeled as “termination” products, and longer transcripts that arose due to transcriptional readthrough are labeled as “readthrough” products. **(c)** Band intensities of total SgrS transcripts (termination+readthrough) were measured, and 5S-normalized values were plotted as “SgrS abundance” (steady state transcript abundance of SgrS mutants). **(d)** Band intensities of SgrS termination and readthrough products were measured and 5S-normalized ratios (readthrough/termination) were calculated and plotted for each SgrS mutant as “SgrS readthrough ratio”.

### Super-resolution imaging of specific chromosomal SgrS mutants

A set of 9 single point mutants of SgrS were chosen for further analysis using quantitative imaging (A177U, G178A, G178U, G184A, U181A, U182A, G215A, U224A, U224G). To avoid potential complications arising from SgrS overexpression, we created these mutations in the endogenous chromosomal copy of SgrS. A177, G178 and U181 are in the seed (target base-pairing) region, G184 is in the small stem-loop region. U181 and U182 are in the U-rich region upstream of the small stem-loop, previously shown to bind Hfq^27^. We also constructed a double-mutant G184A-C195U which restores the small stem-loop structure. G215 is in the terminator stem-loop region and U224 is in the poly-U tail, both of which provide major binding sites for Hfq. These mutant alleles in the background of strains with wild-type RNase E or a C-terminally truncated RNase E were grown, and glucose-phosphate stress was induced using αMG for a varied amount of time before cell fixation and permeabilization. We performed two-color 3D super-resolution imaging of the SgrS sRNAs labeled with up to 9 FISH probes conjugated to Alexa Fluor 647 and the *ptsG* mRNAs labeled with up to 28 FISH probes conjugated to CF568. Δ*sgrS* and Δ*ptsG* strains were also examined to correct for the background arising from nonspecific binding of FISH probes. The wild-type strain showed an increase in SgrS copy number over time after sugar stress induction (Fig. 4, Supplementary Fig. 3). At the same time, the copy number of *ptsG* mRNA showed a decrease (Fig. 4, Supplementary Fig. 3). We used a density-based clustering algorithm^34^ to determine the copy numbers of RNAs along with the copy number of SgrS-*ptsG* mRNA complexes. Super-resolution imaging was especially important for quantifying sRNA-mRNA complexes because at conventional microscopy resolution there was too much false co-localization between sRNA and mRNA.

**Figure 4.**
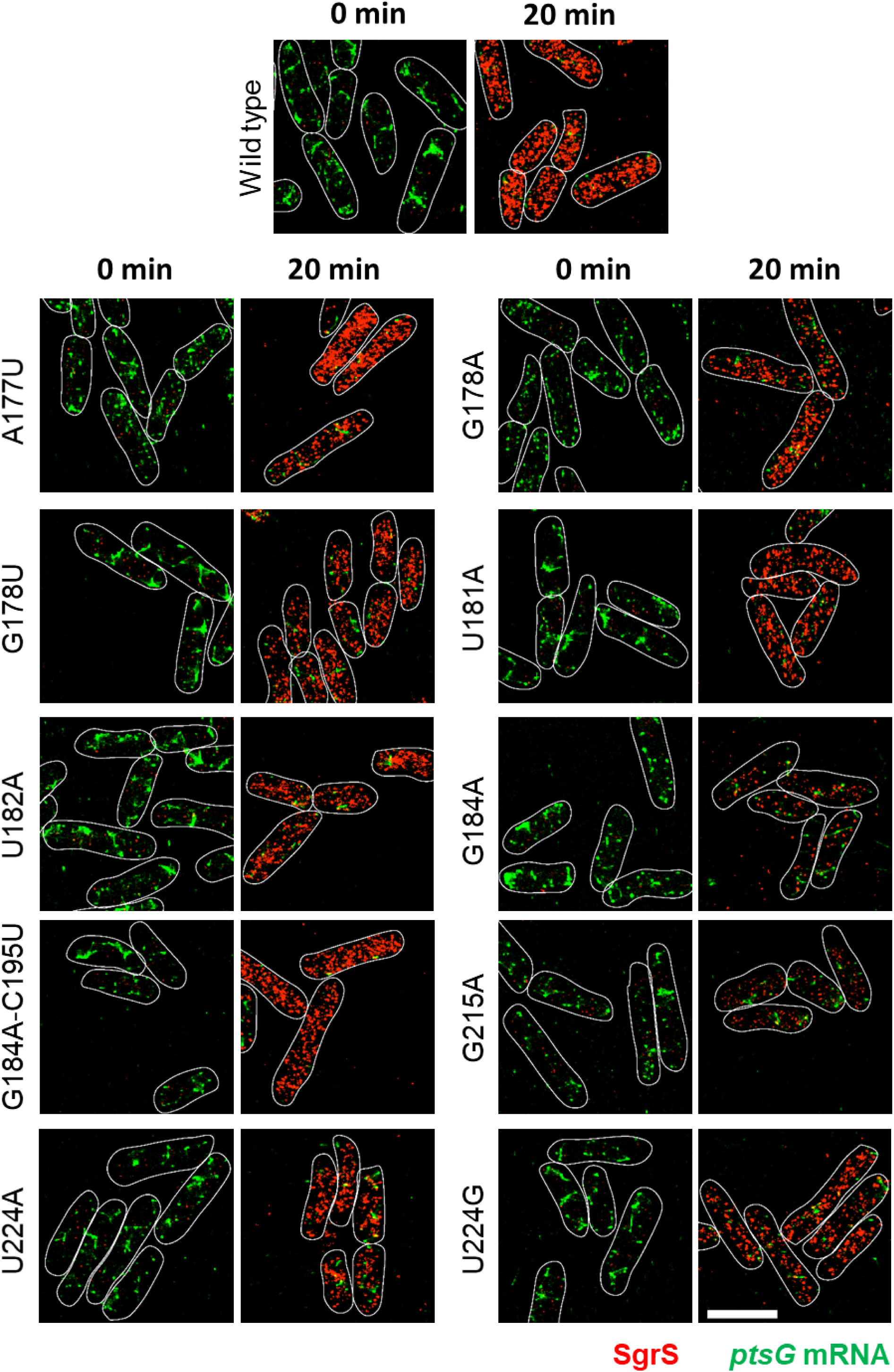
3D super-resolution images projected in 2D planes. The panels show SgrS (red) and *ptsG* mRNA (green) labeled by smFISH for the wild-type and the mutant strains, A177U, G178A, G178U, U181A, U182A, G184A, G184A-C195U, G215A, U224A, U224G before and after 20 minutes of αMG (non-metabolizable sugar analogue) induction. Cell boundaries are denoted by white solid lines. Scale bar is 2 μm.

The accumulation of the mutant SgrS sRNAs was lower than for wild-type SgrS with an accompanying impairment in *ptsG* mRNA degradation for all single point mutants examined, showing that their regulatory functions are perturbed (Fig. 4, Supplementary Fig. 4-9, 11-13). The single cell distribution of RNA copy numbers also showed a decreased accumulation of SgrS, with the histogram peaking at lower copy numbers 20 min after αMG induction, and the histograms for *ptsG* mRNA peaked at higher copy numbers per cell compared to the wild-type (Fig. 5b, d, h, Supplementary Fig. 52). The lowest accumulation of SgrS was seen for G184A and G215A, and they also showed the most impaired mRNA degradation (Fig. 4, 5a, c, g, Supplementary Fig. 9, 11). These two mutations occur in two separate stem-loop regions, both of which participate in Hfq binding.^27,28^ The double mutant, G184A-C195U, which restores base-pairing in the small stem-loop via a compensatory mutation, eliminated the negative impact of G184A as seen by recovery of SgrS accumulation and regulation of *ptsG* mRNA (Fig. 4, 5e, f, Supplementary Fig. 10). This suggests that the disruption of the stem-loop structure, not of G184 basepairing with *ptsG* mRNA, is primarily responsible for regulatory defects of G184A.

**Figure 5.**
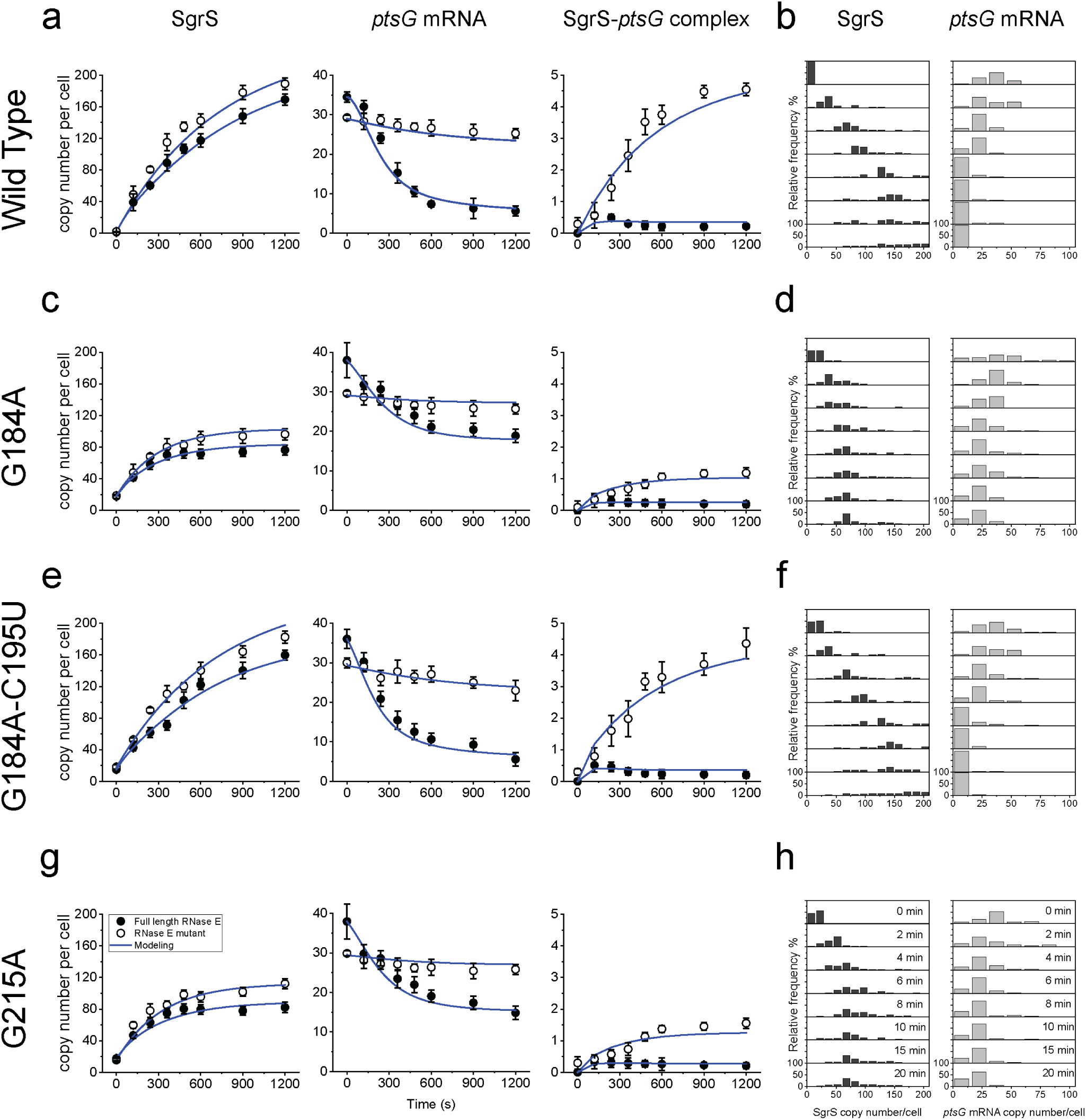
Time dependent changes in the copy numbers of SgrS and *ptsG* mRNA and estimation of kinetic parameters. Time course changes and corresponding modeling curves for the SgrS, *ptsG* mRNA and SgrS-*ptsG* complex in **(a)** wild-type, **(c)** G184A mutant strain, **(e)** G184A-C195U mutant strain, **(g)** G215A mutant strain. Average copy numbers per cell are plotted against time. Rate constants obtained for these mutants are shown in Figure 6 and in Supplementary Table 3. Weighted *R*^*2*^’s for modeling are also reported in Supplementary Table 4. Error bars in **(a), (c), (e), (g)** are standard errors from 80 to 150 cells in each case. Histograms showing the change in distribution of SgrS and *ptsG* mRNA copy numbers for **(b)** wild-type, **(d)** G184A mutant strain, **(f)** G184A-C195U mutant strain, **(h)** G215A mutant strain for 80-150 cells in each case.

These imaging data by themselves cannot tell us whether regulatory defects are due to changes in target binding kinetics or due to changes in the SgrS stability. Therefore, we next determined the lifetimes of wild-type and mutant SgrS molecules.

### Intrinsic lifetimes of SgrS mutants

In order to calculate the target-independent lifetime of SgrS, we induced SgrS expression using αMG and then added rifampicin to stop transcription globally. RT-qPCR was performed vs time after rifampicin treatment to quantify the SgrS level. The wild-type SgrS showed minimal intrinsic degradation over a period of 2 hours after the addition of rifampicin but it showed rapid degradation in the presence of ongoing transcription (10.4 ± 0.7 min), suggesting that SgrS degradation is normally dominated by co-degradation with its various target mRNAs (Fig. Supplementary Fig. 25). The intrinsic degradation was also minimal for SgrS A177U mutant (Supplementary Fig. 25), suggesting that in the absence of co-degradation, a mutation in the target-annealing region does not destabilize SgrS. In contrast, intrinsic degradation of G184A was rapid (lifetime of 6.3 min) and so was the intrinsic degradation of wild-type SgrS in Δ*hfq* strain (lifetime of 5.1 min) (Supplementary Fig. 25), indicating that Hfq is required for the target-independent stability of SgrS and the small stem-loop is important for Hfq binding.

### Lifetime of SgrS mutants

In order to determine the effective lifetime of SgrS mutants, which includes the contributions from intrinsic degradation and co-degradation with target mRNA, the strains carrying chromosomal mutations were treated with αMG for 10 minutes before rinsing it away. SgrS decay over time was then monitored through imaging of fixed cells. The wild-type SgrS showed a degradation rate of 0.0016 ± 0.0001 s^-1^ (lifetime of 10.4 ± 0.7 min) and all of the mutants showed higher rates, the highest being for G184A with 0.0046 ± 0.0003 s^-1^ (lifetime of 3.6 ± 0.2 min), followed by G215A with 0.00345 ± 0.0003 s^-1^ (lifetime of 4.8 ± 0.4 min) (Fig. 6a, Supplementary Fig. 26-32, 34-36). Because G184A and G215A disrupt the small and terminator stem-loop regions, respectively, our data suggest that both stem-loop regions are important for SgrS stability *in vivo*. G184A-C195U recovered the stability of SgrS to the wild-type level with an identical degradation rate within error (Fig. 6a, Supplementary Fig. 33). Rifampicin-chase experiments did not show any difference in the lifetime of *ptsG* mRNA between all mutant strains (Supplementary Fig. 26-37), showing that the mutations in SgrS have no effect on *ptsG* mRNA stability when SgrS is not induced.

**Figure 6.**
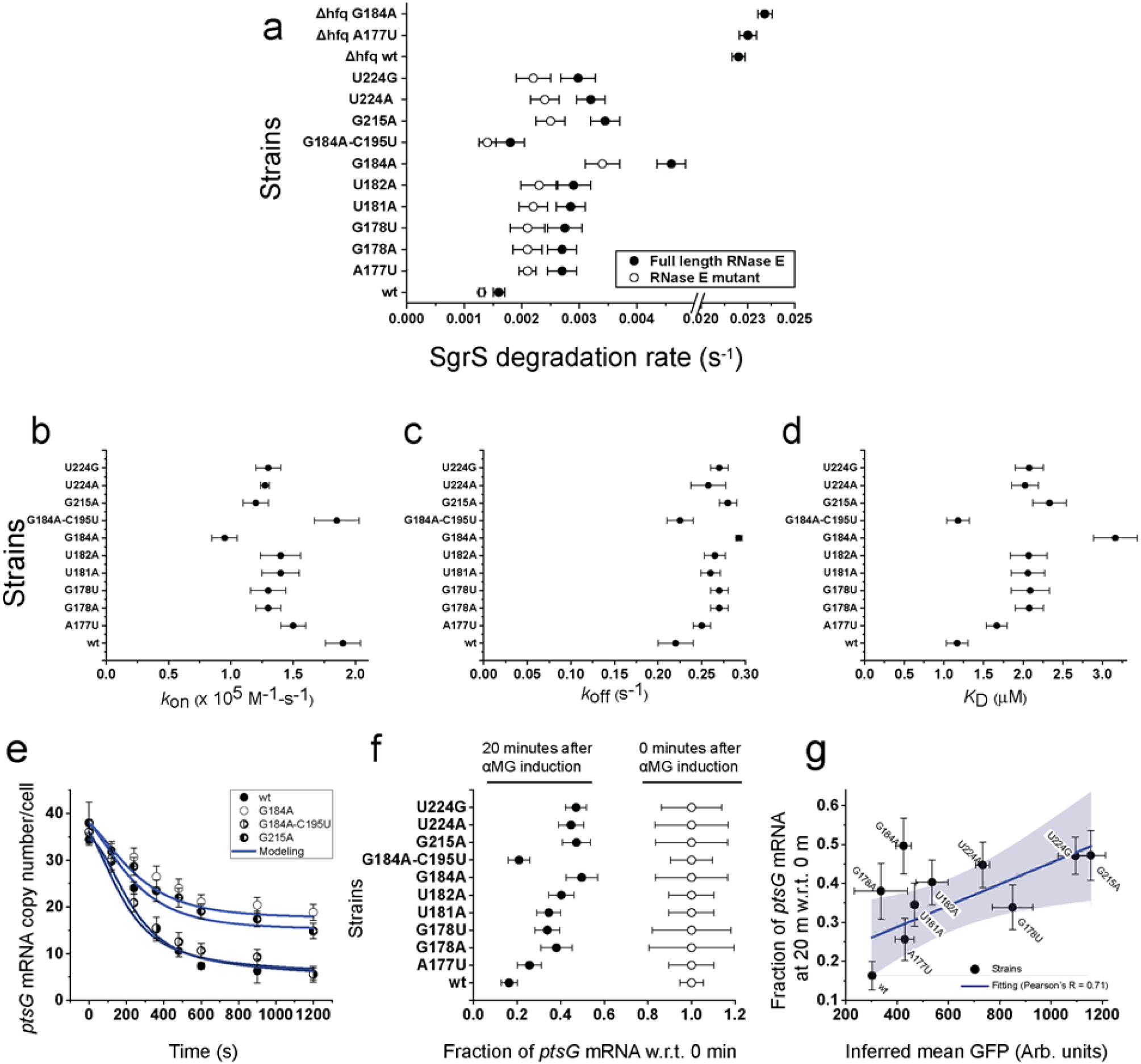
Calculation of various parameters and correlation with Sort-Seq. **(a)** Degradation rates of SgrS for the wild-type and the strains A177U, G178A, G178U, U181A, U182A, G184A, G184A-C195U, G215A, U224A, U224G, Δ*hfq* wild type, Δ*hfq* A177U, Δ*hfq* G184A for full length RNase E and RNase E mutants. Error bars represent standard deviation from two experimental replicates. **(b-d)** *k*_on_, *k*_off_, *K*_D_ measured from the time dependent modeling curves of the SgrS, *ptsG* mRNA and SgrS-*ptsG* mRNA complexes for the wild-type and strains A177U, G178A, G178U, U181A, U182A, G184A, G184A-C195U, G215A, U224A, U224G. These were determined simultaneously in the wild-type and RNase E mutants. Error bars report standard deviation from the independent fitting on two replicates. **(e)** Time course changes in *ptsG* mRNA for the wild-type, G184A, G184A-C195U, G215A mutant strains. Error bars represent standard errors from 80-150 cells in each case. **(f)** Fractional change in *ptsG* mRNA copy numbers for the wild-type and the mutants A177U, G178A, G178U, U181A, U182A, G184A, G184A-C195U, G215A, U224A, U224G before and after 20 minutes αMG induction. Error bars represent standard errors from 80-150 cells in each case. **(g)** Comparison of the SgrS regulation efficacy calculated from Sort-Seq assay and the imaging-based analysis. Error bars in the x-axis are standard deviations calculated from two experimental replicates and those in the y-axis are as described in **(f)**. The fitting is shown in blue and the grey region shows the 95% confidence region.

The degradation rate of SgrS in Δ*hfq* strains was much higher, about 0.022 ± 0.004 s^-1^ (lifetime of 0.76 ± 0.14 min), for wild-type and all SgrS mutants (Fig. 6a, Supplementary Fig. 38-40) ^23^. This 14-fold increase in degradation rate for sRNAs in Δ*hfq* strains confirms that Hfq is indispensable for the stability of SgrS^27,44^. Because none of the SgrS mutants in the *hfq*^*+*^ cells showed degradation rates as high as in Δ*hfq* strains, these SgrS mutations are only partially deleterious to the interactions with Hfq.

Mutations in the base-pairing regions (A177U, G178U and G178A) reduced the lifetime of SgrS in the imaging-based experiment even though they are not expected to alter Hfq binding. Because our Northern blot analysis of overexpressed SgrS showed that these mutations do not change SgrS abundance, the intrinsic degradation is unlikely to be affected by the mutations. Instead, we attribute the discrepancy to mutation-induced alterations in co-degradation of SgrS with other SgrS target mRNAs.

### Target search and destruction kinetics of SgrS mutants

Once we obtained the average copy numbers of SgrS, *ptsG* mRNA and the SgrS-*ptsG* complex per cell as a function of time after SgrS induction for the full length RNase E and RNase E mutant cases (Supplementary Fig. 14-24), we used a previously developed deterministic kinetic model to describe the SgrS-*ptsG* regulation kinetics (Fig. 1a)^34^. We used the experimentally-determined degradation rate for *ptsG* mRNA, *β*_*p*_, to calculate the *ptsG* transcription rate *α*_p_ using *α*_*p*_ = *β*_*p*_ × [*p*]_0_, where [*p*]_0_ is the steady state copy number of *ptsG* mRNA at t=0. By globally fitting the six time courses of the three RNA species with or without RNase E mutation that inhibits co-degradation, we obtained *k*_on_, *k*_off_ and *k*_cat_ for the wild-type and mutant SgrS. *k*_*on*_ for the wild-type strain was (1.9 ± 0.2) x 10^5^ M^-1^-s^-1^ and *k*_*off*_ was 0.22 ± 0.02 s^-1^ giving a *K*_*D*_ of 1.16 ± 0.14 μM, comparable to the previously published results (Fig. 6b-d)^34^. *k*_*on*_ was lower for all single point mutants compared to the wild-type and the reduction ranged from 24% for A177U to 53% for G184A. *k*_*off*_ was higher for all the mutants and the increase ranged from 14% for A177U to 33% for G184A giving a dissociation constant, *K*_*D*_ of 1.67 ± 0.13 μM and 3.08 ± 0.33 μM, respectively (Fig. 6b-d). Rate of transcription of SgrS and *k*_cat_ were not affected by the mutations within error (Supplementary Fig. 42-43). To test the possibility that the apparent changes in *k*_on_ and *k*_off_ are due to fitting errors and that the regulatory deficiencies can be explained solely by reduction in SgrS lifetimes, we repeated the global fitting procedure while keeping the *k*_on_ and *k*_off_ values fixed at the wild-type values. The fits were considerably worse, and were especially poor for copy number curves of *ptsG* mRNA and SgrS/mRNA complex (Supplementary Fig. 44-51). Therefore, our procedure of obtaining the mutation effects on *k*_on_ and *k*_off_ is robust.

When we restored the base-pairing in the small stem-loop by adding a compensatory mutation to G184A, the mutant that showed the largest changes to *k*_on_ and *k*_off_ (G184A-C195U), *k*_on_ and *k*_off_ returned to the wild-type values within error (Fig. 6c-d). Nucleotides 168-187 in SgrS were originally proposed to participate in base-pairing with the *ptsG* mRNA^20^ but a subsequent study showed that only nucleotides 168-181 are required for basepairing^33^. Here we found that binding kinetics is similar between the wild-type and G184C-C195U, strongly suggesting that a mutation at G184 primarily acts through disruption of the small stem-loop structure, thereby affecting Hfq binding, instead of through direct disruption of base-pairing of G184 with the target strand. Even though our data suggest that G184 is not involved with SgrS-*ptsG* mRNA base-pairing, its mutation negatively affected annealing kinetics, decreasing *k*_on_ and increasing *k*_off_. Therefore, our results support the dual roles of Hfq: first to increase sRNA stability (Fig. 6a) and second to directly facilitate SgrS-*ptsG* binding.

The A to U mutation at position 177 removes an AU base-pair, breaking 8 base-pairs, the longest stretch of contiguous base-pairing between SgrS and *ptsG* mRNA into segments of 4 and 3 base-pairs. This disruption gives a reduction in association rate of 24% and an increase in dissociation rate by 14%. The two mutations at position 178 eliminate a GC base-pair and breaking the same 8 base-pairs into segments of 3 and 4 base-pairs. G178A and G178U mutants gave a reduction of association rate by 31-32% and an increase of dissociation rate by 23-25%. The larger effects of G178 mutations compared to A177U are likely due to the loss of GC over AU base-pair. Consistent with this suggestion, a mutation at U181, losing an AU base-pair, decreased the association rate by 26% and increased the dissociation rate by 18%, very similar to A177U values.

The G215A mutation in the terminator stem-loop and the mutations U224A and U224G in the poly-U tail showed *k*_on_ decreases and *k*_off_ increases even though they should not change complementarity between SgrS and *ptsG* mRNA. The substantial effects on binding kinetics must therefore be due to defects in Hfq’s ability to facilitate the annealing reaction, further providing *in vivo* evidence of direct facilitation of base-pairing between sRNA and mRNA by Hfq.

### Difference in regulation outcome between imaging and Sort-Seq experiments

To examine if the regulation outcomes for SgrS mutants is consistent between our quantitative imaging experiments and Sort-Seq analysis, we used the fractional decrease of *ptsG* mRNA over the first 20 min after sugar stress induction as a measure of the SgrS regulation of *ptsG* mRNA target in imaging-based analysis. (Fig. 6e-g). Plotting these values vs the inferred GFP signals obtained from the Sort-Seq experiments, we observed a relatively weak correlation (Pearson’s R = 0.71), suggesting that the translation reporter-based Sort-Seq method is not able to fully capture the regulation defects of SgrS mutations. For example, G178A which had a large deficiency in regulation in imaging experiments showed almost the wild-type level regulation in Sort-Seq. A large defect in regulation was shown in previous studies where nucleotide 178 was mutated and our imaging-based approach is in accordance to this finding^24^. SgrS overexpression in Sort-Seq may have overcome the negative effect of mutations through mass action when the defect is primarily in binding kinetics. For the G215A, U224A and U224G mutant strains, however, Sort-Seq showed large regulatory deficiencies, suggesting that their defects cannot be overcome by overexpression. There are several possible explanations. First, because Sort-Seq relies on the translational output, mutations that disrupt translational inhibition but not RNA co-degradation may not be scored well in imaging-based experiments. However, *in vitro* translation experiments showed that mutation of G178 to C eliminates translation inhibition by SgrS^33^, making it unlikely that defects in RNA-RNA annealing do not affect translational repression. Second, even when these mutants can bind Hfq, the complex may be defective in mediating RNA annealing. Third, these mutations also interfere with proper termination as shown by readthrough transcripts (Fig. 3). It was shown previously that the readthrough products of SgrS transcription do not bind Hfq *in vivo* and *in vitro*^43^. Weakening of the terminator stem-loop or reduction of the slippery Us must be causing transcription readthroughs that produce regulation-defective products, and much of the effect of G215A, U224A or U224G may be due to improper termination. It has also been shown that readthrough transcription of SgrS is suppressed under stress conditions, providing an additional layer of regulation^43^. Finally, the incorporation of the GFP in the reporter system may have affected the stability of the mRNA.

## Discussion

Previous studies have measured the effect of mutations in the regulation of mRNA targets of SgrS^45–47^ and have shown the importance of Hfq, sequence complementarity between SgrS and its targets, and RNA secondary structures^27,33,48,49^. Hfq has also been shown to promote structural changes to the RNAs, which in turn helps in the annealing and, consequently, regulation^50–52^. Our study provides a quantitative description of the process of target search and off-target rejection by determining the kinetic parameters as a function of single nucleotide changes in functionally important regions. The *k*_on_ and *k*_off_ values determined in this study depict the apparent rate constants because we did not explicitly include Hfq binding in our model. *k*_on_ in particular should have contributions from Hfq binding to SgrS, target search by SgrS/Hfq and subsequent annealing.

We used IntaRNA^53–56^ to predict the energy of interaction between SgrS and *ptsG* mRNA and found that it changes by ∼6.4 kcal/mol for the G178 point mutations whereas the change is only around ∼4.3 kcal/mol for A177U. Our study agreed with the ranking because we saw lower rates of association and higher rates of dissociation for G178 than A177U. However, the magnitude of the effect is much more modest compared to a simple prediction based on the energetic penalty. Both mutations introduce a mismatch within eight contiguous base-pairs, incurring large energetic penalties. For example, 6.4 kcal/mol would correspond to a change in the equilibrium binding constant by a factor ∼60,000 instead of ∼2 we observed. Therefore, Hfq must be buffering the effect of breaking internal base-pairs in short helices. How this is achieved is presently unknown.

We found that the rate of co-degradation remains high, ∼0.3 s^-1^, even with SgrS mutations. The co-degradation of the SgrS-*ptsG* complex is brought about by the degradosome, in which RNase E is a key component. Hfq copurifies with RNase E and SgrS^57^, and at least one sRNA (MicC) has been shown to mediate the interaction between Hfq and the C-terminal part of RNase E *in vitro*^58^ and *in vivo*^59^. It has been hypothesized that the sRNA-Hfq-RNase E complex forms first and subsequently the complementary mRNA binds to this complex, aided by Hfq, followed by a coupled or sequential degradation of the RNA pair^58^. The changes in *k*_on_ and *k*_off_ in our study account for the disruption of the SgrS-*ptsG* mRNA annealing, but once a stable complex with all four components forms, the co-degradation occurs at the same rate irrespective of the SgrS mutations.

Because *k*_cat_ did not change with SgrS mutations, the probability that a single binding event will cause co-degradation of sRNA and mRNA decreases with a mutation-induced increase in *k*_off_. On average, the wild-type SgrS would take 1.73 (= (*k*_off_+*k*_cat_)/*k*_cat_) binding events before co-degradation. This number increases to 1.81 when AU basepairing is disrupted by a mutation and increases further to 1.9 when GC basepairing is disrupted. In addition, the binding rate at the full SgrS accumulation condition, 100-200 copies per cell, would correspond to 0.48 μM (assuming 0.7 μm^3^ per cell), and the wild-type SgrS would take about 11 s to bind *ptsG* mRNA. If a mutant SgrS were present at the same concentration, it would take ∼13.6 s and ∼14.4 s to bind for disrupting a single AU and GC base-pair, respectively. The overall time it takes to degrade the target would increase from 19 s to 24.6 s and 27.4 s for disrupting an AU and GC base-pair, respectively. Although we examined only the effect of SgrS mutation in this study, if we assume that a mutation in the target mRNA breaking a single base-pair has a similar impact, we can conclude that target search and destruction will take 29-44 % longer for nearly cognate off-target RNA containing a single base-pair mismatch.

The poly-U sequence at the 3’ end of sRNAs is an important Hfq binding module^29,43^ and binds to the proximal face of the ring-shaped Hfq hexamer^29,50^. Because Hfq forms a stable 1:1 complex with SgrS^27^, a single Hfq hexamer must bind the poly-U tail, both of the stem-loops, and the UA-rich region upstream of the small stem-loop simultaneously. How this is achieved requires further structural studies, but the UA-rich region has been found in other sRNAs and has been proposed to bind the rim of the Hfq hexamer^27,51,60–62^. The binding of Hfq with target mRNAs, however, has been studied extensively and it is known that Hfq brings about a distortion in the mRNA structure, promoting the base-pairing between the RNAs^63,64^. In this study we showed that the U224 mutations in the poly-U tail caused the rate of association of the RNAs to decrease and the rate of dissociation to increase. Because U224 is distant from the mRNA annealing region of SgrS, our data showed that Hfq directly facilitates RNA-RNA annealing *in vivo*. The same effect was observed from other mutants that disrupt Hfq binding without changing the mRNA annealing region of SgrS, and collectively our work presents the first *in vivo* evidence that Hfq directly facilitates target binding. It should be emphasized that a careful accounting of SgrS mutations’ effects on SgrS lifetimes was necessary to reach this conclusion. Microscopic mechanisms for Hfq’s role in sRNA-mRNA annealing are still a subject of active research^60,65,66^, and may be investigated in the future using our analysis platform.

## Materials and Methods

### Construction of plasmids for Sort-Seq studies

A *ptsG-sf*GFP reporter system was constructed, containing 105 nt 5’ UTR and 30 nt coding sequence of *ptsG* mRNA, which coded for the first 10 amino acids of PtsG protein, and this was fused by a 42 nt linker sequence and the superfolder GFP coding sequence. The reporter system was subcloned from the pZEMB8 plasmid. A plasmid, pAS06 was constructed by inserting this reporter sequence into the low copy plasmid pAS05 between the XhoI and XbaI restriction sites and the expression of the reporter system was under the control of P_Llac-O1_.

The SgrS sRNA sequence was inserted in between the NdeI and BamHI restriction sites of the medium copy plasmid pZAMB1 and its expression was under the control of P_Ltet-O1_. The sgrS mutation library was prepared by using the plasmid pZAMB1 as a template for mutagenesis PCR and also as a vector to insert the *sgrS* mutation sequence.

### Cell culture and induction for Sort-Seq studies

The *E. coli* MB1 strain (Δ*ptsG*, Δ*sgrS, lacI*^*q*^, *tetR*) was transformed with plasmids (pAS06 for *ptsG*-*sf*GFP and pZAMB1 for *sgrS* or the SgrS mutation library) and grown at 37 °C in LB Broth Miller (EMD) overnight with the respective antibiotics (100 μg/ml ampicillin (Gold Biotechnology, Inc.)) for pAS06 plasmid and 30 μg/ml chloramphenicol (Sigma-Aldrich) for pZAMB1 plasmid and the *sgrS* mutation library). The following day, the cell culture was diluted 200-fold into fresh LB Broth with respective antibiotics and were grown until OD_600_ reached 0.1-0.2 as measured using an Educational Spectrophotometer (Fisher Scientific Education). The culture was diluted again to an OD_600_ of 0.001 and supplemented with 1 mM IPTG (Sigma-Aldrich) to induce the expression of PtsG-*sf*GFP and 50 ng/ml aTc to induce the expression of SgrS or the SgrS mutation library. The *E. coli* cells were collected and treated further for the next set of experiments.

### SgrS sRNA mutagenesis experiment

Agilent Genemorph II Random Mutagenesis Kit (Agilent Technologies) was used to perform mutagenesis PCR on SgrS using the protocol adapted from previously published work from Levine’s lab^39^. 1 ng of pZAMB1 plasmid, with the *sgrS* sequence, was used to conduct mutagenesis PCR for 15 cycles. The yields of the individual mutants were increased by amplifying the product using Phusion® High-Fidelity DNA Polymerase (New England Biolabs) and purified using QIAquick Spin Columns (Qiagen). The PCR products were then digested with NdeI (New England Biolabs) and BamHI (New England Biolabs) and purified by QIAquick Spin Columns (Qiagen). The pZAMB1 vector was also prepared by digestion with NdeI and BamHI followed by purification using QIAquick Spin Columns (Qiagen). The vector and the PCR insert were used to prepare 4 ligation reactions by mixing with T4 Ligase (New England Biolabs).

The products from all the reactions were combined and purified using QIAquick Spin Columns (Qiagen) into water. 5 μl of the purified ligation product was then transformed into MB1 strain, which was pre-transformed with pAS06 plasmid expressing *ptsG*-*sf*GFP. These transformed cells were then recovered and diluted into LB Broth supplemented with 100 μg/ml ampicillin and 30 μg/ml chloramphenicol and grown overnight at 37 °C. The following day, the culture was centrifuged, aliquoted as frozen stocks and used for imaging and flow cytometry experiments.

### Epifluorescence Imaging

1 ml culture of the *E. coli* strain to be imaged was grown from an overnight culture till OD_600_ = 0.1-0.2. It was then chilled on ice followed by centrifugation at 6000 g, 4 °C for 1 minute to form a cell pellet. Then they were washed with ice-cold 1X PBS twice and resuspended in 100 μl 1X PBS.

1.5% (w/v) agarose gel was prepared by dissolving agarose in 1X PBS. A few μl cell suspension was sandwiched between a No. 1.5 glass coverslip (VWR) and a thin slab of the agarose gel. The sample was then imaged.

The epifluorescence images were acquired by a Nikon Ti Eclipse microscope (Nikon Instruments, Inc.) using an oil immersion objective (1.46 NA, 100X) which spans an area of around 133×133 μm^2^ for DIC (no filter, autofluorescence) and fluorescence imaging (Ex 480-500 nm, Em 509-547 nm, exposure time 200 ms). The images were acquired using an EMCCD camera (Andor). They were processed using the NIS-Element AR software (Nikon Instruments, Inc.).

### Fluorescence-Activated Cell Sorting

The *E. coli* strain to be sorted was cultured overnight in LB Broth with appropriate antibiotics. The following day, the liquid culture was diluted 200-fold and cultured with antibiotics until OD_600_ = 0.1-0.2. The cells were then diluted to OD_600_ = 0.001 in LB Broth with antibiotics and 1 mM IPTG and/or 50 ng/ml aTc were added corresponding to the strain of *E. coli* and the plasmids it is carrying. They were grown till OD_600_ = 0.1-0.2, washed with ice-cold 1X PBS twice and kept on ice before flow cytometry analysis or fluorescence-activated cell sorting. The sorting and analysis were done in a MoFlo XDP Cell Sorter (Beckman Coulter) using a 488nm 200mW laser.

### Preparation of the sample for sequencing

The cells sorted into the batches were grown in LB Broth supplemented with 30 μg/ml chloramphenicol to saturation. We extracted the plasmids with E.Z.N.A. Plasmid Mini kit (Omega, D6942-02). To generate sequencing amplicons, we followed Illumina 16S sequencing protocol. We used 5 ng of each plasmid elute as PCR template and amplified out the portion of interest using 0.5 μM of primers annealing to the region of the *sgrS* sequence under consideration with Phusion 2X Mastermix (NEB, M0531L). We employed 20 cycles of 10 s at 98 °C denaturation, 20 s at 63 °C primer annealing, 10 s at 72 °C elongation phases preceded by additional initial denaturation at 98 °C for 30 s and followed by 72 °C final extension for 2 min. To clean up the product, we incubated the PCR product with 20 μl Ampure XP beads (Beckman Coulter, A63880) for 5 min. We retained the bead-bound material after keeping for 2 minutes on a magnetic rack (GE, 1201Q46). We washed the beads twice with 80% ethanol, air-dried for 10 min and eluted the material in 53 μl 10 mM Tris pH 8.5 by incubation for 2 min. We collected 45-50 μl bead-free liquid 2 min after placing the material on a magnetic rack.

### Illumina Next-Gen Sequencing

We performed 8 additional cycles of PCR with Nextera 24-Index kit for indexing before sample pooling (Illumina, FC-121-1011), for which we used 7.5 μl of the above elute as template, 7.5 μl each of the suitable i5 and i7 primers with 38 μl Phusion 2X Mastermix. We followed manufacturer’s recommended thermal cycling protocol (95 °C 3min, 98 °C 30s, 55 °C 30s, 72 °C 30s, 72 °C 5min). We also bead-purified 55 μl of this final product with 56 μl Ampure-XP beads and eluted with 28 μl 10 mM Tris pH 8.5 buffer. We pooled the final products based on their Nanodrop reading at equal molar stoichiometry and diluted the sample down to 4 nM in 10 mM Tris buffer. We alkaline-denatured by mixing 5 μl DNA sample with 0.2 M NaOH and incubating for 5 min at room temperature. We diluted this product down to 20 pM in Hbf buffer. We loaded a final mixture of 465 μl Hbf buffer, 120 μl pooled 20 pM library and 15 μl denatured 20 pM PhiX control library (Illumina, FC-110-3001) after a 2 minute heat treatment at 96 °C followed by a 5 min incubation on ice. We used a 150 cycles MiSeq v3 reagent kit (Illumina, MS-102-3001) to perform a single-end sequencing for 150 cycles. We used the manufacturer’s default algorithm for base calling and de-multiplexing of the constituent samples.

### Intensity Moment Calculation

We parsed the raw .fastq output files via a simple home-made C++ script compiled with GCC v. 7.5 and plotted with GNU Octave v. 4.2. The analysis scripts can be accessed via the Gitlab page https://gitlab.com/tuncK/sortseq/-/tree/master and the raw data can be obtained from TH upon request.

We only imported the base calls of each read, thus including all sequences regardless of their quality factors. We directly extracted from each read the subsequence excluding the PCR adaptors, i.e. bases 23 to 128. Out of this list of subsequences, we detected ones that are exact duplicates of each other by building a red-black binary search tree. Among all such groups, we only considered SgrS sequence variants that are represented by at least 10 distinct reads in the data. We compared the observed sequence of each group with the wild-type SgrS sequence i.e. that of the plasmid used as error-prone PCR template.

We normalized the raw number of reads of each group by both the total number of reads and the fraction of cells falling under each gate. As such, we defined a weighted average intensity to each individual mutant along this 106 base long SgrS segment that we probed. Referred to as the “intensity moment” from now on, we calculated the following quantity:

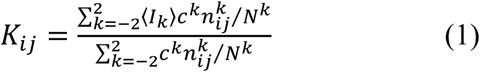

where, K_ij_ is the intensity moment of the mutant carrying a single substitution mutation at the i’th base position to nucleotide type j rather than the wt base. c^k^ is the overall fraction of cells that are sorted into the k’th bin based on the GFP intensity histogram that FACS acquisition software reports. n^k^ is the number of reads carrying a single substitution mutation at base position i to base type j and detected in the k’th FACS bin. N^k^ is the total number of acceptable reads in the dataset. 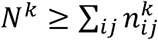 due to experimental errors as well as reads carrying multiple substitutions due to the stochastic nature of error-prone PCR. ⟨*I*_*k*_⟩ is the median intensity of the cells falling into the k’th bin as reported by FACS. For the representative intensity of each bin, we used the median intensity reported by the FACS device.

In the figures, we reported the standard score of each entry given by

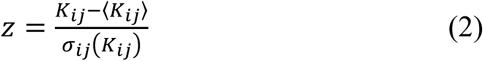

### Construction of bacterial strains

The oligonucleotides, plasmids, and strains used in this study are listed in Supplementary Tables 1 and 2. *E. coli* K12 MG1655 derivatives were used for all experiments. P1 transduction^67^ or λ-red recombination^68^ were used to move alleles between strains. DNA fragments were PCR amplified using Q5® Hot Start High-Fidelity 2X Master Mix (NEB) and oligonucleotides described in Supplementary Table 2. A set of plasmids (Supplementary Table 1) were used as templates to PCR amplify the wild-type and *sgrS* mutants A177T, G178T, G178A, and G184A using single-stranded oligos (Supplementary Table 2) containing 5’ and 3’ homology to the flanking regions of *cat-sacB* cassette (MB205). DNA fragments containing the *sgrS* mutants G215A, T224G, T224A, T181A, and T182A were PCR amplified from MG1655 genomic DNA using oligonucleotides listed in Supplementary Table 2.

Translational *ptsG’-’lacZ* reporter fusion under the control of the P_BAD_ promoter (strain MB130) was constructed by PCR amplifying fragment of interest using primer pair MBP201F/MBP201R containing 5’ homologies to P_BAD_ and *lacZ*. PCR product was recombined into *E. coli* PM1205 using λ Red-mediated homologous recombination and counter-selection against *sacB* as described previously^69^. Marked *λattB::lacI*^*q*^*-PN25tetR-spec*^*R*^ was introduced into MB130 strain by P1 *vir* transduction^70^ to produce MB168 strain (Supplementary Table 1). Plasmid pZAMB1 harboring *sgrS* under the control of the P_LtetO-1_ promoter was constructed by PCR amplifying *sgrS* from *E. coli* MG1655 chromosomal DNA using oligonucleotides containing NdeI and BamHI restriction sites. PCR products and vector pZA31R^71^ were digested with NdeI and BamHI (New England Biolabs) restriction endonucleases. Digestion products were ligated using DNA Ligase (New England Biolabs) to produce plasmid containing P_LtetO-1_-*sgrS* allele^40^. Single nucleotide mutations in SgrS were introduced by QuikChange mutagenesis procedure using oligonucleotides with mismatched bases at desired locations as following: A177T (A177T-F/ A177T-R), G178T (G178T-F/G178T-R), G178A (G178A-F/G178A-R), G184A (G184A-F/ G184A-R), G215A (G215A-F/ G215A-R), T224A (U224A-F/U224A-R), T224G (T224G-F/T224G-R) (Supplementary Table 2).

### β-galactosidase assay

Bacterial strains were cultured overnight in MOPS Rich medium with 25 μg ml^-1^ chloramphenicol (Cm) and subcultured 1:100 to fresh MOPS Rich medium containing Cm and 0.0005% L-arabinose. Cells were grown at 37°C with shaking to OD_600_∼0.15 and 30 ng ml^-1^ anhydrotetracycline (aTc) was added to induce expression of SgrS from the plasmid and cells grown for another hour to OD_600_∼0.5. β-galactosidase assay was then performed according to previously described protocol^70^.

### Northern blot analysis

Bacterial strains were cultured and β-galactosidase assay performed as described above. Simultaneously, aliquots of the same culture were taken and total RNA was extracted as described previously^72^. RNA concentrations were measured spectrophotometrically and 15 μg of RNA were resolved on 6% polyacrylamide gel electrophoresis. RNA was transferred to 0.2 μm pore-size Nytran N (Whatman) membrane as described previously^73^. Membrane was prehybridized for 45 min in ULTRAhyb (Ambion) solution at 42°C. Blots were probed overnight with 5’-biotinylated SgrS-bio or ssrA-bio probes specific for SgrS sRNA and 5S rRNA respectively (Supplementary Table 2). BrightStar BioDetect kit (Ambion) was used for detection. ImageJ (National Institutes of Health; ^74^) was used to measure band densities from four independent experiments.

### Measurement of intrinsic degradation rates of SgrS

Strain DB166, MB206, MB209 and XM199 were cultured overnight at 37 °C and diluted 1:100 to a fresh LB medium and the cultures were grown at 37 °C to OD_600_ ∼ 0.3. To induce SgrS expression, αMG was added to final concentration of 0.5% and the cells were grown for additional 30 minutes. Rifampicin was added to final concentration of 250 μg/ml and the cells were grown for another 5 minutes. At this point, cells were harvested (t=0 timepoint) for RNA extraction. Three biological replicates were harvested for each time-point.

Cells from 1.0 ml of culture were mixed with 2 ml of RNA protect reagent (Qiagen). The mixture was pelleted at 4000 rpm for 10 minutes and then discard the supernatant. Total RNA was isolated using Direct-Zol RNA miniPrep (Zymo) kit following the manufacturer’s instruction. Genomic DNA was removed by DNaseI provided by the Kit. Finally total RNA was eluted in 40 μl of nuclease free water. First-strand cDNA was synthesized from 1 μg of total RNA using Superscript™ IV First-Strand cDNA Synthesis SuperMix kit according to the manufacturer’s protocol (Invitrogen, USA).

The primers used to amplify SgrS are: OSA499 (GATGAAGCAAGGGGGTGCCC) and OSA500 (CAATACTCAGTCACACATGATGCAGGC)

The primers used to amplify housekeeping gene *rrsA* are: OXM187 (ATTCCGATTAACGCTTGCAC) and OXM188 (AGGCCTTCGGGTTGTAAAGT)

Real-time PCR was performed using SYBR Green master mix (Fisher) and Eppendorf Realplex in a 96-well plate. Each reaction is comprised of 1x SYBR Green master mix, 100 nM of each primer, 2 μl of 1:50 diluted cDNA in a total of 10 μl reaction volumes. Each plate contains “no template” controls for individual transcripts as well as housekeeping transcripts such as *rrsA* for every sample as an internal control.

Delta delta Ct method was used to analyze the qPCR data. The transcripts turnover rates were calculated based on the non-linear fit with one phase exponential decay curves using GraphPad software.

### Cell culture, fixation and permeabilization for smFISH and super-resolution imaging

The wild-type *E. coli* strain (DJ480) was grown overnight at 37 °C, 250 rpm in LB Broth Miller (EMD), the RNase E mutant was grown in 25 µg/ml kanamycin (Kan) (Fisher Scientific), the SgrS A177U, G178U, G178A, U181A, U182A, G184A, G184A-C195U, G215A, U224A, U224G mutants were grown in LB Broth with 50 μg/ml spectinomycin (Spec) (Sigma-Aldrich) and the RNase E mutants of the respective SgrS mutations were grown in LB Broth with 25 μg/ml kanamycin and 50 μg/ml spectinomycin. The following day, the overnight cultures were diluted 100-fold into MOPS EZ rich defined medium (Teknova) with 0.2% glucose and the respective antibiotics, and allowed to grow at 37 °C and 250 rpm until the OD_600_ reached 0.15-0.25. α-methyl D-glucopyranoside (αMG) (Sigma-Aldrich) was used to introduce sugar-phosphate stress and subsequently induce SgrS sRNA expression. A specific volume of liquid was taken out of the culture after 0, 2, 4, 6, 8, 10, 15, 20 minutes of incubation and mixed with formaldehyde (Fisher Scientific) to a final concentration of 4% for the fixation of the cells.

Δ*sgrS* and Δ*ptsG* strains were grown overnight in LB Broth Miller (EMD) at 37 °C and 250 rpm using 25 μg/ml kanamycin and 10 μg/ml tetracycline (Tet) (Sigma-Aldrich) respectively. The next day the cultures were diluted 100-fold into MOPS EZ rich defined medium (Teknova) with 0.2% glucose (Sigma-Aldrich) and the respective antibiotics and left to grow at 37 °C and 250 rpm again till the OD_600_ reached 0.2. The cells were then mixed with formaldehyde (Fisher Scientific) to a final concentration of 4% to fix the cells.

Following the formaldehyde fixation, the cells were incubated at room temperature for 30 minutes and subsequently centrifuged at 3214 x g for 10 minutes at room temperature. The pellets were resuspended in 200 μl 1X PBS and then washed 3 times, each time performing centrifugation at 600 x g for 4 minutes and resuspending in 200 μl 1X PBS. The cells were then permeabilized with 70% ethanol, shaken at room temperature for 1 hour and stored at 4 °C prior to fluorescence *in situ* hybridization.

### Single-molecule fluorescence *in situ* hybridization (smFISH)

Stellaris Probe Designer was used to design the smFISH probes and they were ordered from Biosearch Technologies (https://www.biosearchtech.com/). The probe labeling was performed by using equal volumes of each probe. The final volume of sodium bicarbonate was adjusted to 0.1 M by adding 1/9 reaction volume of 1 M sodium bicarbonate (pH = 8.5). 0.05-0.25 mg of Alexa Fluor 647 succinimidyl ester (Life Technologies) or CF 568 succinimidyl ester (Biotium) dissolved in 5 μl DMSO was mixed with the probe solution. The dyes were kept at a molar excess of 20-25 fold relative to the probes. The reaction mixture was incubated in the dark at 37 °C with gentle vortexing overnight. The following day the reaction was quenched by using 1/9 reaction volume of 3 M sodium acetate (pH = 5). Ethanol precipitation followed by P-6 Micro Bio-Spin Columns (Bio-Rad) were employed to remove unconjugated dyes.

60 μl of permeabilized cells were centrifuged at 600 x g for 4 minutes and the pellets were washed with FISH wash solution (10% formamide in 2X Saline Sodium Citrate (SSC) buffer). They were then resuspended along with the probes in 15 μl of FISH hybridization buffer (10% dextran sulfate (Sigma-Aldrich), 1 mg/ml *E. coli* tRNA (Sigma-Aldrich), 0.2 mg/ml Bovine Serum Albumin (BSA) (NEB), 2 mM vanadyl ribonucleoside complexes (Sigma-Aldrich), 10% formamide (Fisher Scientific) in 2X SSC). The number of probes used for sRNA SgrS was 9, they were labeled with Alexa Fluor 647 and the concentration of the labeled probes was 50 nM. The number of probes used for *ptsG* mRNA was 28, they were labeled with CF 568 and the labeled probe concentration was 15 nM. The reaction mixtures were incubated in the dark at 30 °C overnight. The following day, the cells were suspended in 20X volume FISH wash solution and centrifuged. They were resuspended in FISH wash solution, incubated at 30 °C for 30 minutes and centrifuged, and this was repeated 3 times. After the final washing step, the cells were pelleted and resuspended in 20 µl 4X SSC and stored at 4 °C prior to imaging.

### Single-molecule localization-based super-resolution imaging

The labeled cells were immobilized on 1.0 borosilicate chambered coverglass (Thermo Scientific Nunc Lab-Tek) treated with poly-L-lysine (Sigma-Aldrich) and imaged with imaging buffer (50 mM Tris-HCl (pH = 8.0), 10% glucose (Sigma-Aldrich), 1% β-mercaptoethanol (Sigma-Aldrich), 0.5 mg/ml glucose oxidase (Sigma-Aldrich) and 0.2% catalase (Sigma-Aldrich) in 2X SSC).

3D super-resolution imaging was performed using an Olympus IX-71 inverted microscope with a 100X NA 1.4 SaPo oil immersion objective. Sapphire 568-100 CW CDRH (568 nm) (Coherent) and DL-640-100-AL-O (647 nm) (Crystalaser) were used for two-color imaging and DL405-025 (405 nm) (Crystalaser) was used for the reactivation of the dyes. The laser excitation was controlled by mechanical shutters (LS6T2, Uniblitz). The laser lines were reflected to the objective using a dichroic mirror (Di01-R405/488/561/635, Semrock) The emission signal was collected by the objective and then they passed through an emission filter (FF01-594/730-25, Semrock for Alexa Fluor 647 or HQ585/70M 63061, Chroma for CF 568) and the excitation laser was cleaned using notch filters (ZET647NF, Chroma; NF01-568/647-25×5.0, Semrock and NF01-568U-25, Semrock). The images were captured on a 512×512 Andor EMCCD camera (DV887ECS-BV, Andor Tech). 3D imaging was achieved by introducing astigmatism using a cylindrical lens with focal length 2 m (SCX-50.8-1000.0-UV-SLMF-520-820, CVI Melles Griot) in the emission path between two relay lenses of focal lengths 100 mm and 150 mm. Each pixel corresponded to 100 nm in this setup. The z-drift of the setup was controlled by the CRISP (Continuous Reflective Interface Sample Placement) system (ASI) and the region of interest for imaging was selected using an xy-sample stage (BioPrecision2, Ludl Electronic Products). The storm-control software written in Python by Zhuang’s group and available at GitHub (https://github.com/ZhuangLab/storm-control) was used for image acquisition.

After acquiring a DIC image of the sample area, two-color super-resolution imaging was performed. 568 nm laser excitation was used for CF 568 after completing the image acquisition for Alexa Fluor 647 using 647 nm laser excitation. Fluorophore bleaching was compensated and moderate signal density was maintained by increasing the 405 nm laser power slowly. Imaging was completed when most of the fluorophores had photobleached and the highest reactivation laser power was reached.

Fluorescent nanodiamonds (140 nm diameter, Sigma Aldrich) were utilized for mapping of the two channels. These nanodiamonds nonspecifically attached to the surface of the imaging chambers and were excited by both 647 nm and 568 nm lasers. They generated localization spots in the final reconstructed images that was used for mapping.

### Image Analysis

The raw data was acquired using the Python-based acquisition software and it was analyzed using a data analysis algorithm which was based on work published previously by Zhuang’s group.^75,76^ The peak identification and fitting were performed using the method described before.^34^ The z-stabilization was done by the CRISP system and the horizontal drift was calculated using Fast Fourier Transformation (FFT) on the reconstructed images of subsets of the super-resolution image, comparing the center of the transformed images and corrected using linear interpolation.

### Clustering Analysis and copy number calculation

A density-based clustering analysis algorithm (DBSCAN) was employed to calculate the RNA copy numbers. The algorithm used was the same as previously published,^34^ but the Nps and Eps values were updated for the SgrS and *ptsG* images, since, we used CF 568 instead of Alexa Fluor 568 and we also used a different 405 nm laser to reactivate the dyes. The SgrS (9 probes labeled with Alexa Fluor 647) images were clustered using Nps = 3 and Eps = 15 and the *ptsG* (28 probes labeled with CF 568) images were clustered using Nps = 10 and Eps = 25 and these numbers were empirically chosen. A MATLAB code was used as before for the cluster analysis. Δ*sgrS* and Δ*ptsG* strains were grown, prepared, imaged and analyzed in the same manner as before and they were used for the measurement of the background signal due to the non-specific binding of Alexa Fluor 647 and CF 568.

The SgrS image with no αMG induction for the wild-type *E. coli* cells (DJ480) was considered to be the low SgrS copy number sample, where it was assumed that one cluster was equivalent to one RNA and the *ptsG* image with 20 minute αMG induction for the wild-type *E. coli* cells was considered to be the low *ptsG* copy number sample. The copy numbers of the RNAs were calculated in the same manner using MATLAB codes as described previously.^34^

### Colocalization analysis

To calculate the copy number of SgrS-*ptsG* complexes, colocalization analysis was performed in order to calculate the percentage of *ptsG* colocalized with SgrS. The average radius of a *ptsG* mRNA cluster was calculated to be around 40 nm. That value was used as the radius to consider a 3D spherical volume from the center of the *ptsG* cluster. The SgrS spots corresponding to clusters found in this volume were taken to be colocalized with the *ptsG* cluster. The base-pairing mutant strain was considered a negative control (Supplementary Fig. 41a) and percentage of colocalization was plotted against SgrS copy number and fit with a line (y = a*x) to act as a calibration for colocalization by chance (Supplementary Fig. 41b). The coefficient, a, was used a correction factor for colocalization calculation as, final colocalization = calculated colocalization – a*SgrS copy number.

### SgrS and *ptsG* mRNA half-life measurements

The *ptsG* mRNA degradation rates were calculated using a rifampicin-chase experiment. The wild-type (DJ480) *E. coli* cells, the SgrS A177U, G178A, G178U, U181A, U182A, G184A, G184A-C195U, G215A, U224A, and U224G were grown in LB Broth with the respective antibiotics at 37 °C, 250 rpm overnight. The following day, the overnight cultures were diluted 100-fold in MOPS EZ rich defined medium supplemented with 0.2% glucose and they were grown at 37 °C, 250 rpm. When the OD_600_ reached 0.15-0.25 rifampicin (Sigma-Aldrich) was added to a final concentration of 500 µg/ml. This was taken as the 0-minute time point for the experiment and aliquots were taken at 2, 4, 6, 8, 10, 15, 20 minutes after the addition of rifampicin and fixed in the same manner described before. The cells were labeled by FISH probes, imaged and analyzed by the same process mentioned. The natural logs of the copy numbers were plotted against time and the slope of the linear fitting was used to calculate the lifetime of the RNA. The reciprocal of the lifetimes gave the degradation rates.

The SgrS degradation rates were calculated for the above strains and the wild-type Δ*hfq*, A177U Δ*hfq*, G184A Δ*hfq* mutants by stopping the transcription of SgrS by removing αMG from the media. The wild-type *E. coli* cells, the mutants and the RNase E mutants were grown overnight as described before in LB Broth with the respective antibiotics. The cells were diluted the following day and grown in MOPS EZ rich defined medium with the respective antibiotics till OD_600_ 0.15-0.25. SgrS transcription was induced in the cells using αMG and growing them for 10 minutes. The cells were then washed twice with centrifugation and resuspension with cold, fresh media devoid of αMG and finally resuspended in pre-warmed media at 37 °C. Aliquots were taken at 0, 2, 4, 6, 8, 10, 15, 20 minutes (0, 2, 4, 6, 8 minutes for the Δ*hfq* strains) and fixed as described before. The cells were then treated, imaged and analyzed to calculate the degradation rates as mentioned before.

### Modeling of SgrS-induced *ptsG* mRNA degradation

#### Kinetic model and experimental measurements of the parameters

The mass-action equations used for the wild-type *E. coli* cells and the chromosomal mutations are shown below:

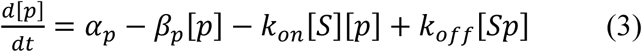

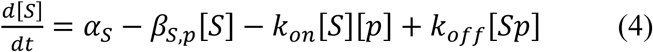

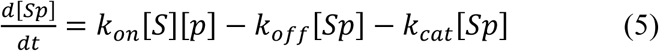

In the above equations, the changes in the concentration of *ptsG*, SgrS and the SgrS-*ptsG* complex over time are shown. *α*_*p*_, *α*_*S*_ are the transcription rates of the *ptsG* mRNA and SgrS respectively; *β*_*p*_, *β*_*S,p*_ are respectively the endogenous degradation rate of ptsG mRNA and the degradation rate of SgrS excluding the co-degradation with *ptsG* mRNA; *k*_*on*_, *k*_*off*_ are the rates of association and dissociation of SgrS and *ptsG* mRNA and *k*_*cat*_ is the RNase E-mediated co-degradation of SgrS-*ptsG* complex.

We calculated the endogenous degradation rate of *ptsG* mRNA (*β*_*p*_) of the wild-type *E. coli*, chromosomal mutations and the RNase E mutants from the super-resolution imaging and analysis. The degradation rate of SgrS for the cells were calculated by stopping the transcription of SgrS, but this method takes into account target-dependent and target-independent degradation (*β*_*S,total*_). We also calculated the degradation rate for the respective RNase E mutant strains and this measurement gave us target-independent degradation and other RNase E–independent degradation (*β*_*S*0_). These two values provided a higher and lower bound for the endogenous degradation rate of SgrS (*β*_*S,p*_).

The transcription rate of *ptsG* mRNA was calculated using *α*_*p*_ = *β*_*p*_ × [*p*]_0_ and in this equation [*p*]_0_ is the concentration of *ptsG* mRNA before the induction of sugar stress in all of the cases. This was done because it was observed previously^34^ that the *ptsG* mRNA reached an equilibrium in the cells without SgrS-induced degradation. We calculated this for all the cases, viz., wild-type *E. coli*, SgrS mutants and the RNase E mutants and the transcription rate of *ptsG* mRNA did not show any significant change.

RNase E mutant cells are not able to degrade SgrS-*ptsG* complex efficiently, but it is a possibility that the complex can degrade endogenously or via other minor degradation pathways. We kept *k*_*cat*_ as a fitting parameter and used the measured parameters, *α*_*p*_, *β*_*p*_, *β*_*S,total*_and *β*_*S*0_ and the above equations to fit the time courses for all the strains to estimate the 5 parameters; *α*_*S*_, *β*_*S,p*_, *k*_*on*_, *k*_*off*_ and *k*_*cat*_.

#### Parameter search

Poisson weighting (total sum of the squares, 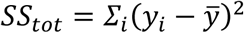 and residual sum of the squares, *SS*_*res*_ = *Σ*_*i*_ (*y*_*i*_ − *f*_*i*_)^2^, where *y*_*i*_ is the experimental data and *f*_*i*_ is the fitted data) was used in the fitting of global *R*^2^ according to the equation:

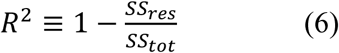

so that no bias was introduced for a particular species. The parameters were selected to maximize the global *R*^2^ for the time course curves of each of the species. The concentrations of the SgrS-*ptsG* complex in all the strains were very close to the background and as a result the total variance became small. *R*^2^ was not helpful to estimate the quality of the fit in these cases. Instead, *χ*^2^’s were calculated as

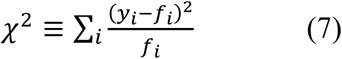

for all those cases and the significance levels (*α*) were reported.

## Acknowledgements

We would like to thank Erel Levine, Divya Balasubramanian, and D. Jin for plasmids and strains. We thank Hao Zhang at the Cell Sorting Core Facility (Bloomberg School of Public Health) for helping us with the flow cytometry and sorting experiments. We appreciate and thank Prof. Sarah Woodson for going through the manuscript and providing insightful suggestions. This work was supported by grants from National Institutes of Health R01 GM112659 (M.B., M.S.A., T.H., J.Z., and A.P.), R35 GM122569 (T.H., J.Z., and A.P.), National Science Foundation PHY 1430124 (T.H., J.Z., and A.P). T.H. is an investigator with the Howard Hughes Medical Institute.

## Author Contributions

A.P., C.K.V., T.H. designed the experiments, with help from M.B. and J.Z. A.P., T.K., and J.Z. performed the Sort-Seq experiments and the Sort-Seq data was analyzed by A.P., T.K., and P.L. M.B. performed the β-galactosidase and Northern blot experiments. M.S.A. and X.M. made the strains that were used in this work. M.S.A. and X.M. performed the qPCR experiments to calculate RNA lifetimes. A.P. performed all super-resolution imaging experiments, with some help from J.Z. A.P. performed the analysis for the imaging experiments with the MATLAB package written by D.S. and J.F. A.P., Z.L.S, C.K.V., and T.H. discussed the data. A.P., T.K., M.S.A., X.M., J.F., C.K.V., and T.H. wrote the manuscript.

## Competing Interests

The authors declare no competing interests.

## Notes

### Competing Interest Statement

The authors have declared no competing interest.

