## Supplementary Information for "Effects of individual base-pairs on *in vivo* target search and destruction kinetics of small RNA"

This document includes:

**Supplementary Figures 1-53**

**Supplementary Tables 1-4**

**Supplementary Text**

**References**

### Supplementary Figures

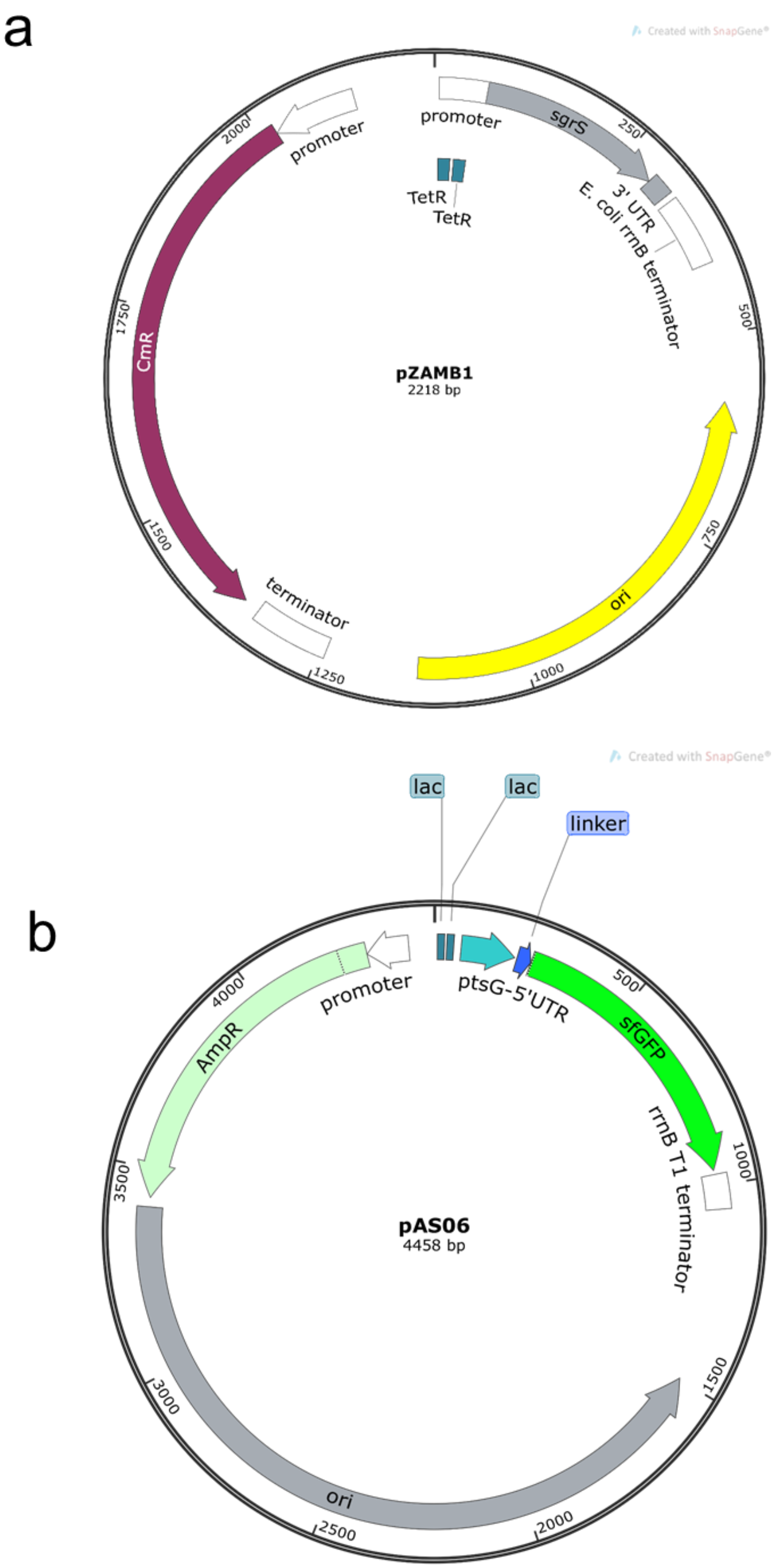

**Supplementary Figure 1. Plasmids used for Sort-Seq. (a)** pZAMB1 plasmid containing wild-type *sgrS* sequence, **(b)** pAS06 plasmid containing *ptsG* 5’ UTR fused to superfolder GFP.

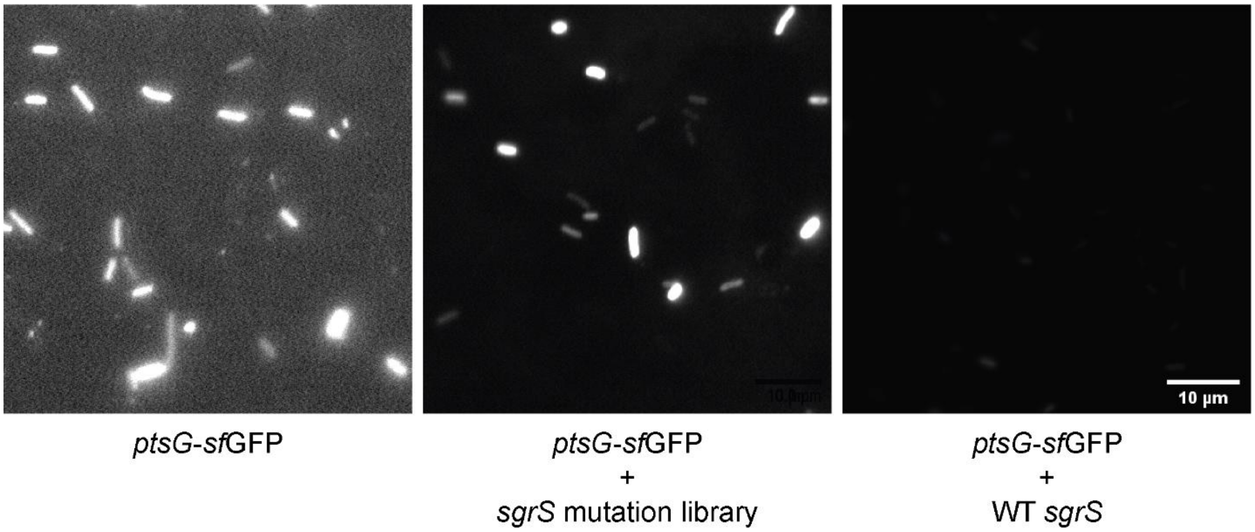


**Supplementary Figure 2. Verification of target-reporter system.** Epifluorescence imaging of cells expressing only ptsG-*sf*GFP, ptsG-*sf*GFP with WT *sgrS* and ptsG-*sf*GFP with *sgrS* mutation library. Scale bar is 10 µm.

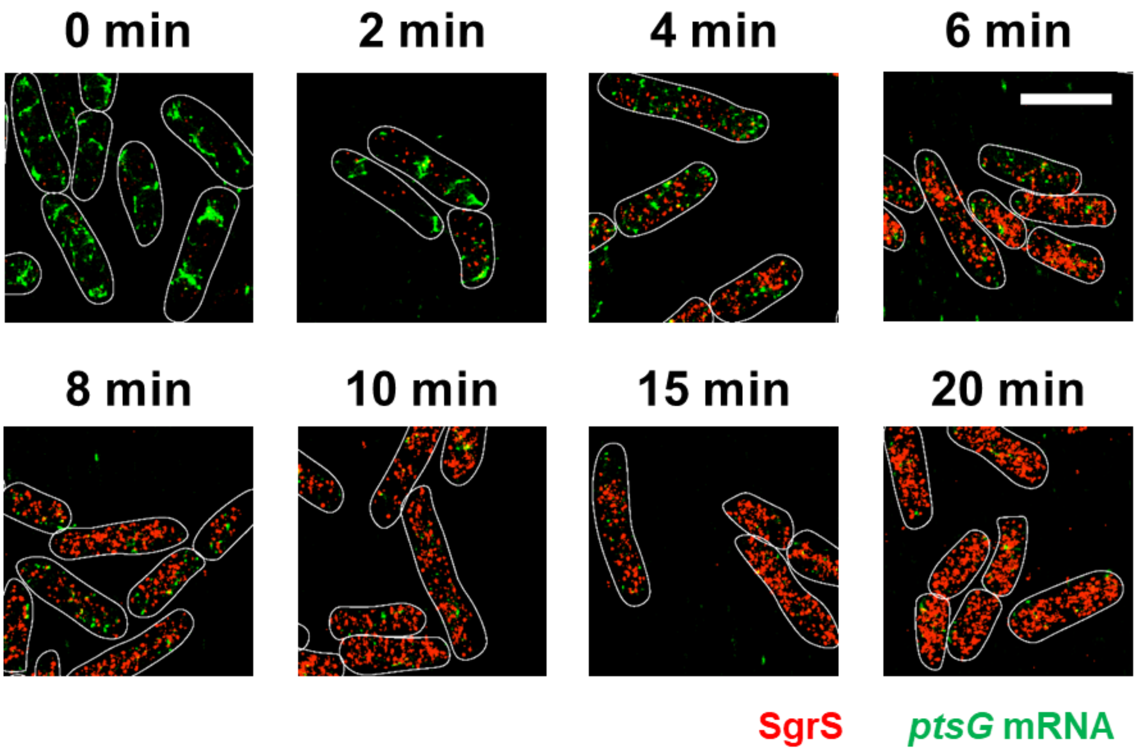


**Supplementary Figure 3. 3D super-resolution images of SgrS (red) and *ptsG* mRNA (green) in the wild-type SgrS strain projected on 2D planes.** The panels show the multi-color images of WT SgrS cells before (0 min) and 2, 4, 6, 8, 10, 15, 20 min after αMG (non-metabolizable sugar analog) induction. White lines denote cell boundaries. Scale bar is 2 μm.

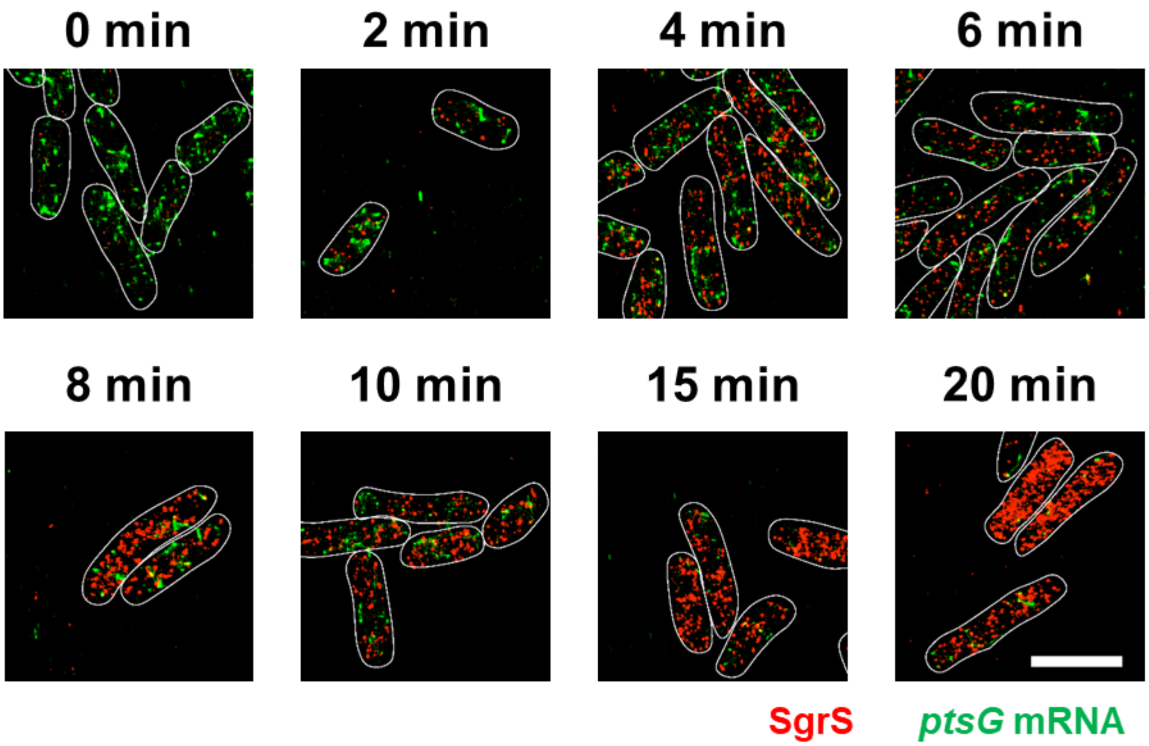


**Supplementary Figure 4. 3D super-resolution images of SgrS (red) and *ptsG* mRNA (green) in the SgrS A177U mutant strain projected on 2D planes.** The panels show the multi-color images of SgrS A177U cells before (0 min) and 2, 4, 6, 8, 10, 15, 20 min after αMG (non-metabolizable sugar analog) induction. White lines denote cell boundaries. Scale bar is 2 μm.

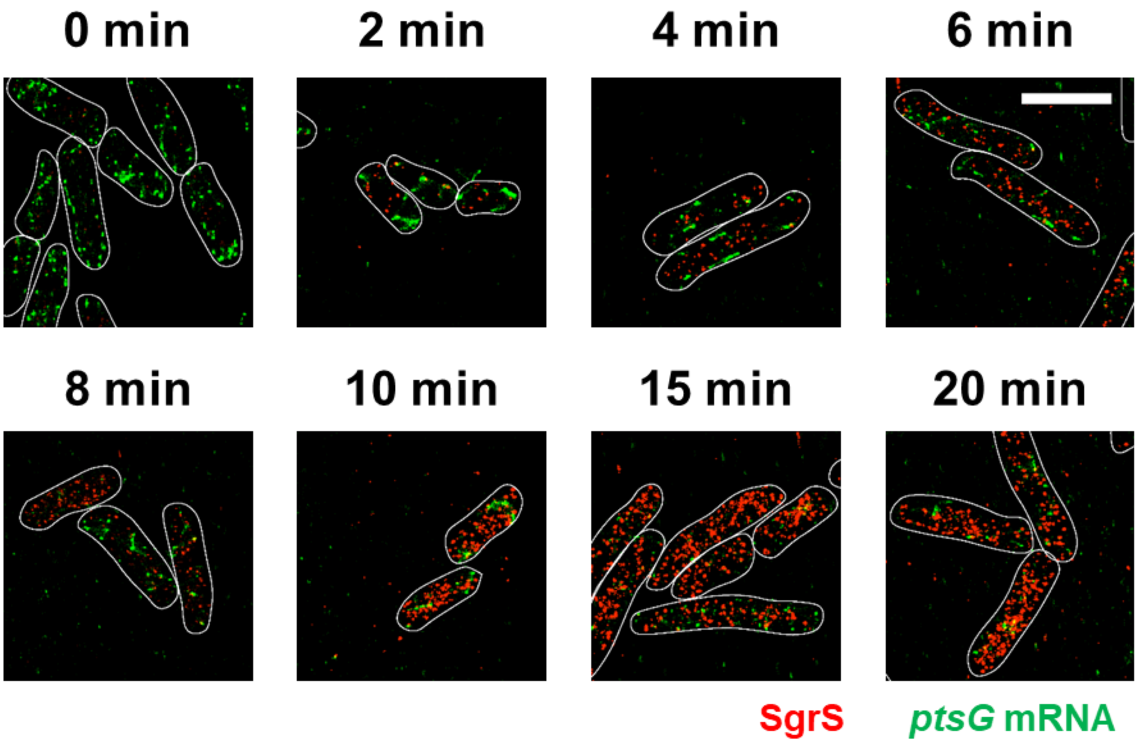


**Supplementary Figure 5. 3D super-resolution images of SgrS (red) and *ptsG* mRNA (green) in the SgrS G178A mutant strain projected on 2D planes.** The panels show the multi-color images of SgrS G178A cells before (0 min) and 2, 4, 6, 8, 10, 15, 20 min after αMG (non-metabolizable sugar analog) induction. White lines denote cell boundaries. Scale bar is 2 μm.

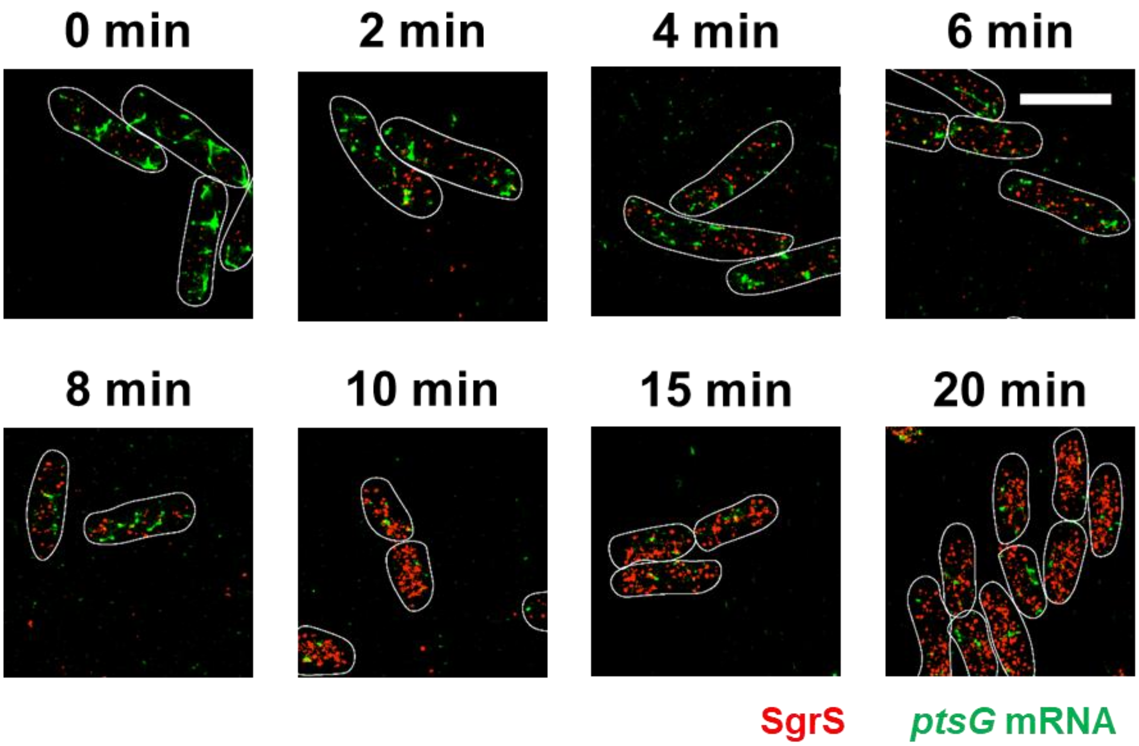


**Supplementary Figure 6. 3D super-resolution images of SgrS (red) and *ptsG* mRNA (green) in the SgrS G178U mutant strain projected on 2D planes.** The panels show the multi-color images of SgrS G178U cells before (0 min) and 2, 4, 6, 8, 10, 15, 20 min after αMG (non-metabolizable sugar analog) induction. White lines denote cell boundaries. Scale bar is 2 μm.

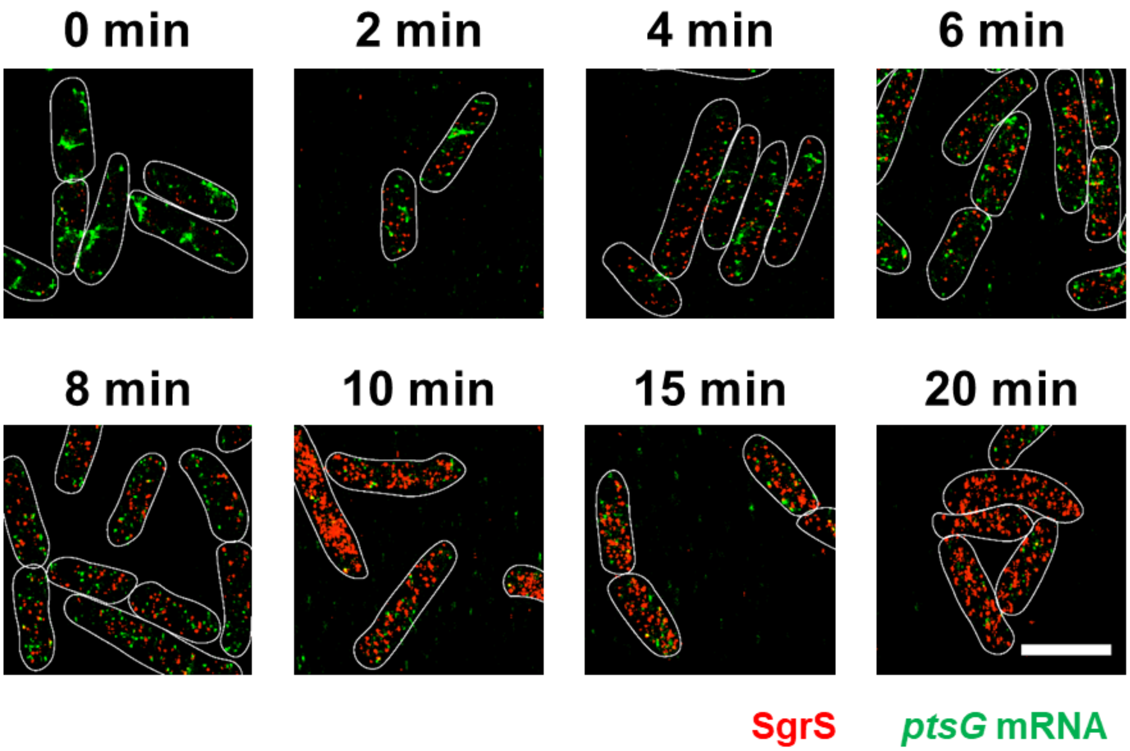


**Supplementary Figure 7. 3D super-resolution images of SgrS (red) and *ptsG* mRNA (green) in the SgrS U181A mutant strain projected on 2D planes.** The panels show the multi-color images of SgrS U181A cells before (0 min) and 2, 4, 6, 8, 10, 15, 20 min after αMG (non-metabolizable sugar analog) induction. White lines denote cell boundaries. Scale bar is 2 μm.

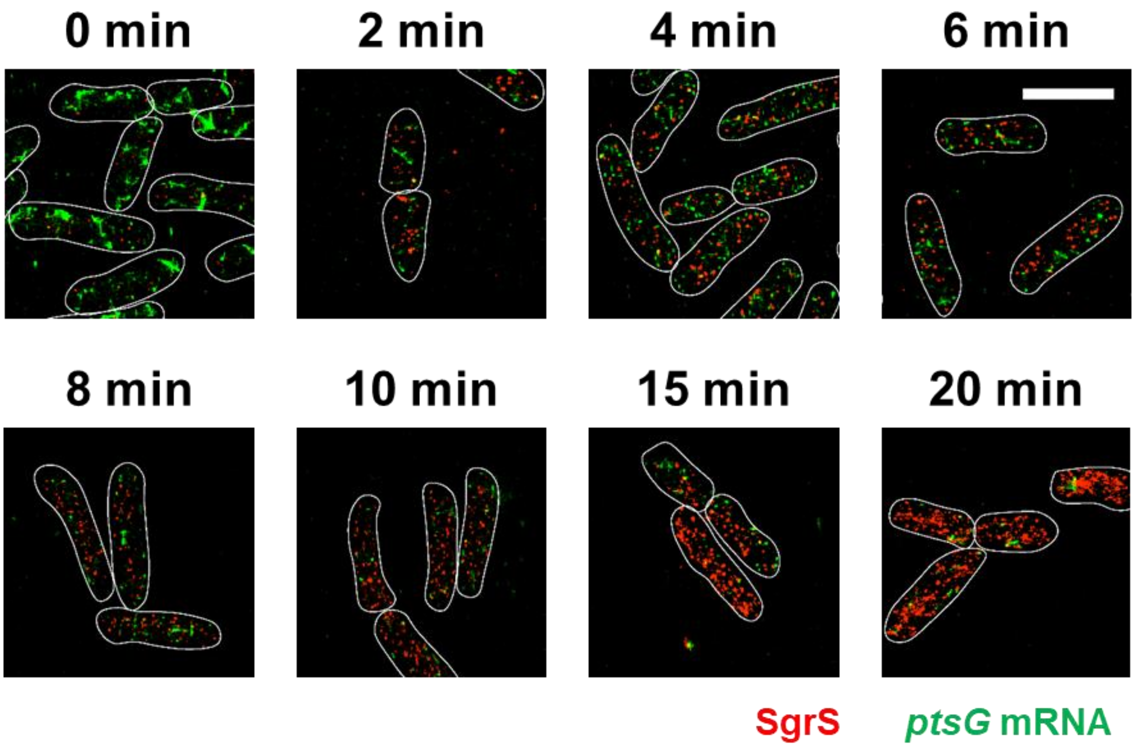


**Supplementary Figure 8. 3D super-resolution images of SgrS (red) and *ptsG* mRNA (green) in the SgrS U182A mutant strain projected on 2D planes.** The panels show the multi-color images of SgrS U182A cells before (0 min) and 2, 4, 6, 8, 10, 15, 20 min after αMG (non-metabolizable sugar analog) induction. White lines denote cell boundaries. Scale bar is 2 μm.

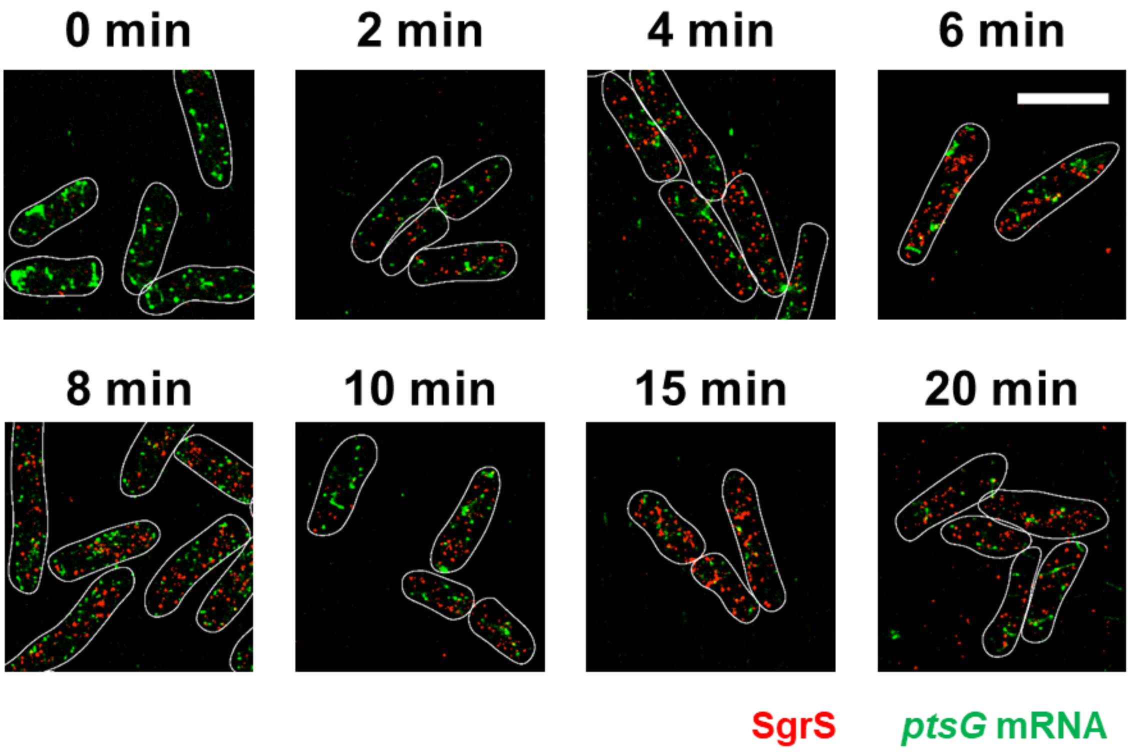


**Supplementary Figure 9. 3D super-resolution images of SgrS (red) and *ptsG* mRNA (green) in the SgrS G184A mutant strain projected on 2D planes.** The panels show the multi-color images of SgrS G184A cells before (0 min) and 2, 4, 6, 8, 10, 15, 20 min after αMG (non-metabolizable sugar analog) induction. White lines denote cell boundaries. Scale bar is 2 μm.

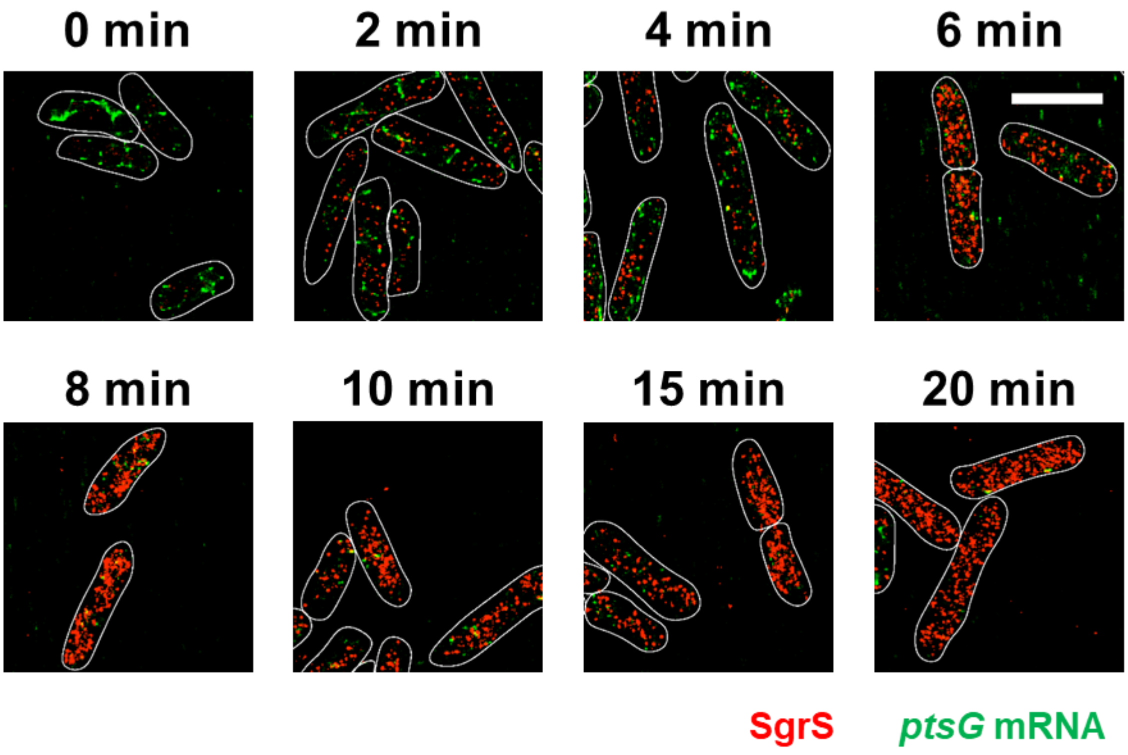


**Supplementary Figure 10. 3D super-resolution images of SgrS (red) and *ptsG* mRNA (green) in the SgrS G184A-C195U mutant strain projected on 2D planes.** The panels show the multi-color images of SgrS G184A-C195U cells before (0 min) and 2, 4, 6, 8, 10, 15, 20 min after αMG (non-metabolizable sugar analog) induction. White lines denote cell boundaries. Scale bar is 2 μm.

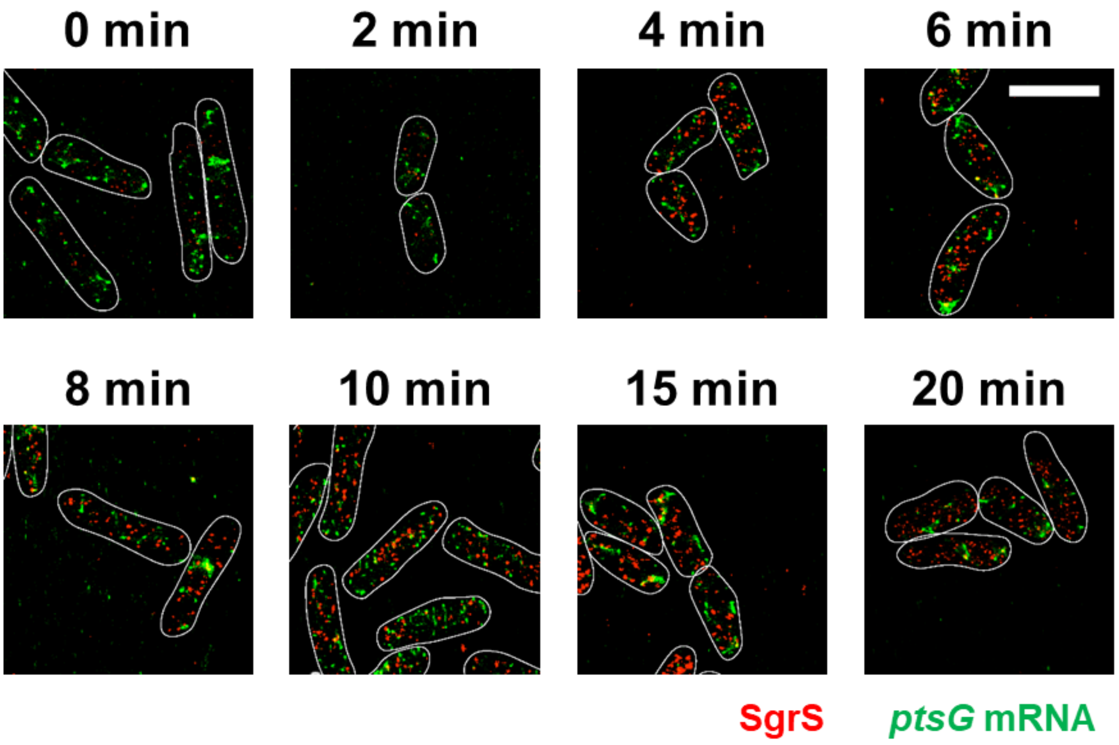


**Supplementary Figure 11. 3D super-resolution images of SgrS (red) and *ptsG* mRNA (green) in the SgrS G215A mutant strain projected on 2D planes.** The panels show the multi-color images of SgrS G215A cells before (0 min) and 2, 4, 6, 8, 10, 15, 20 min after αMG (non-metabolizable sugar analog) induction. White lines denote cell boundaries. Scale bar is 2 μm.

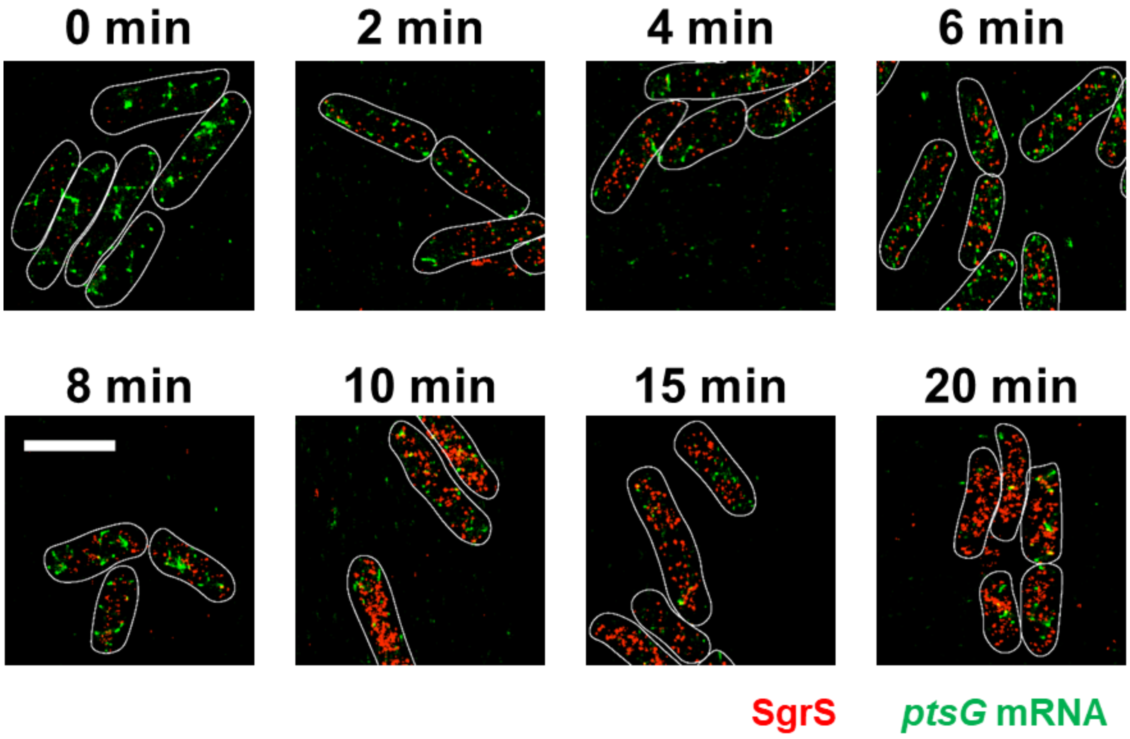


**Supplementary Figure 12. 3D super-resolution images of SgrS (red) and *ptsG* mRNA (green) in the SgrS U224A mutant strain projected on 2D planes.** The panels show the multi-color images of SgrS U224A cells before (0 min) and 2, 4, 6, 8, 10, 15, 20 min after αMG (non-metabolizable sugar analog) induction. White lines denote cell boundaries. Scale bar is 2 μm.

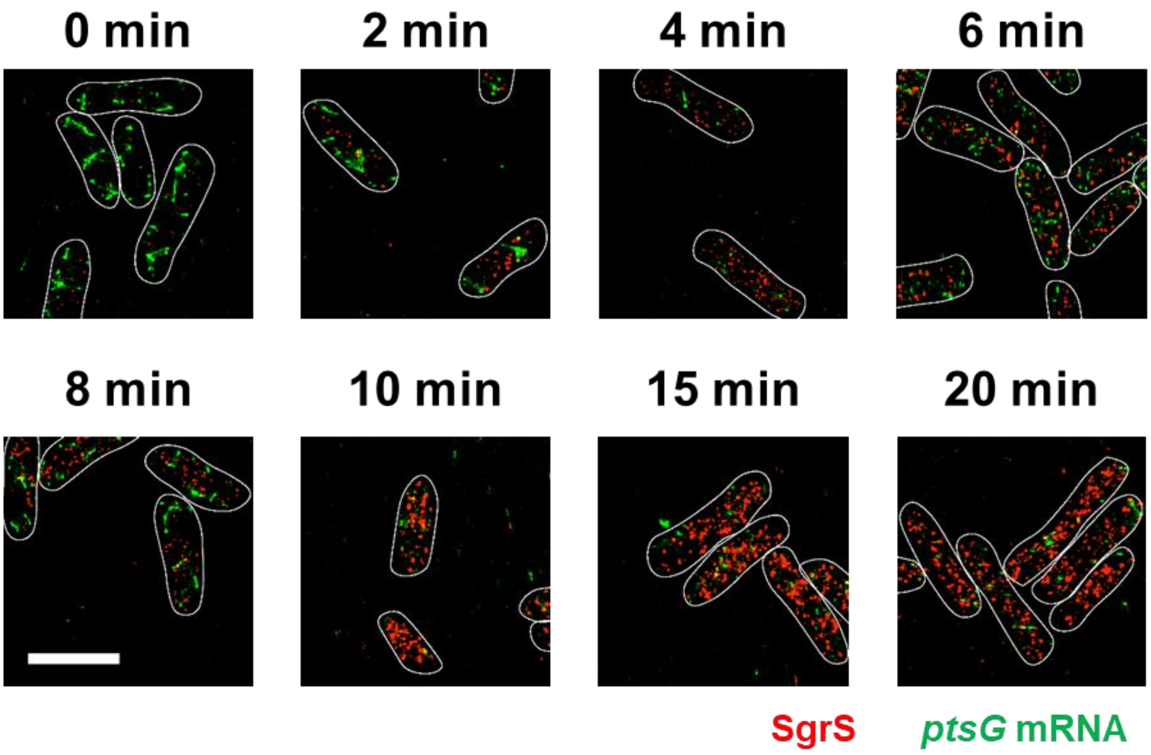


**Supplementary Figure 13. 3D super-resolution images of SgrS (red) and *ptsG* mRNA (green) in the SgrS U224G mutant strain projected on 2D planes.** The panels show the multi-color images of SgrS U224G cells before (0 min) and 2, 4, 6, 8, 10, 15, 20 min after αMG (non-metabolizable sugar analog) induction. White lines denote cell boundaries. Scale bar is 2 μm.

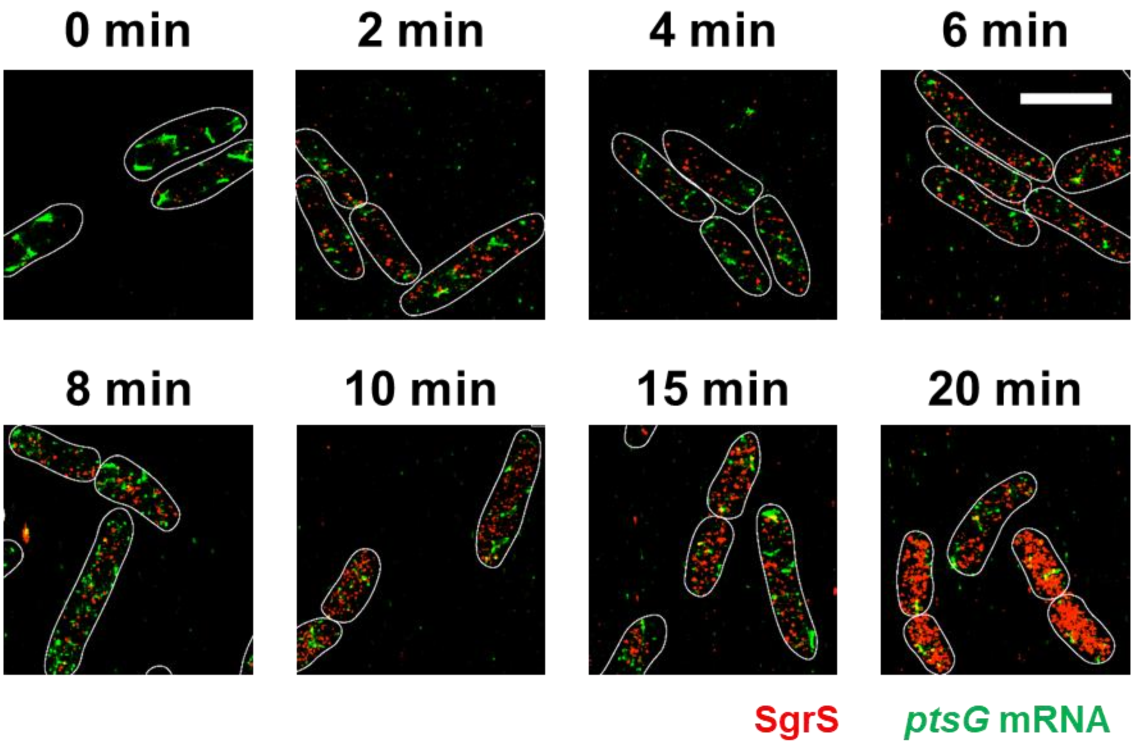


**Supplementary Figure 14. 3D super-resolution images of SgrS (red) and *ptsG* mRNA (green) in the wild-type SgrS RNase E mutant strain projected on 2D planes.** The panels show the multi-color images of WT SgrS RNase E mutant cells before (0 min) and 2, 4, 6, 8, 10, 15, 20 min after αMG (non-metabolizable sugar analog) induction. White lines denote cell boundaries. Scale bar is 2 μm.

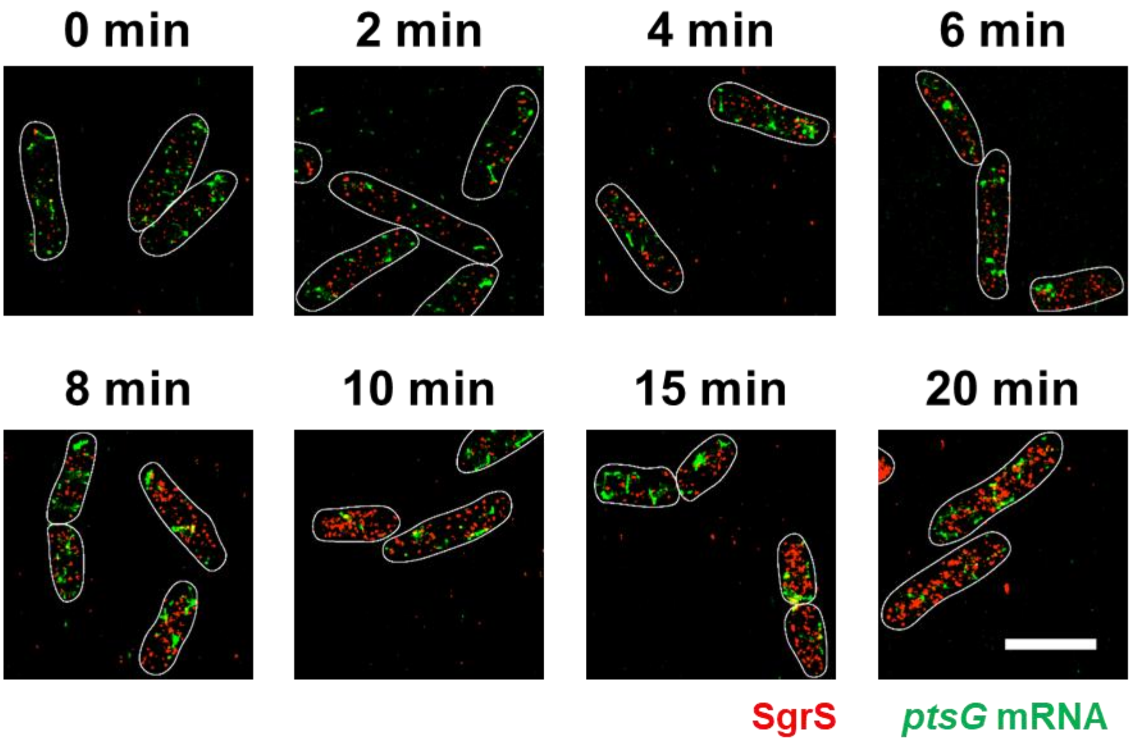


**Supplementary Figure 15. 3D super-resolution images of SgrS (red) and *ptsG* mRNA (green) in the SgrS A177U RNase E mutant strain projected on 2D planes.** The panels show the multi-color images of SgrS A177U RNase E mutant cells before (0 min) and 2, 4, 6, 8, 10, 15, 20 min after αMG (non-metabolizable sugar analog) induction. White lines denote cell boundaries. Scale bar is 2 μm.

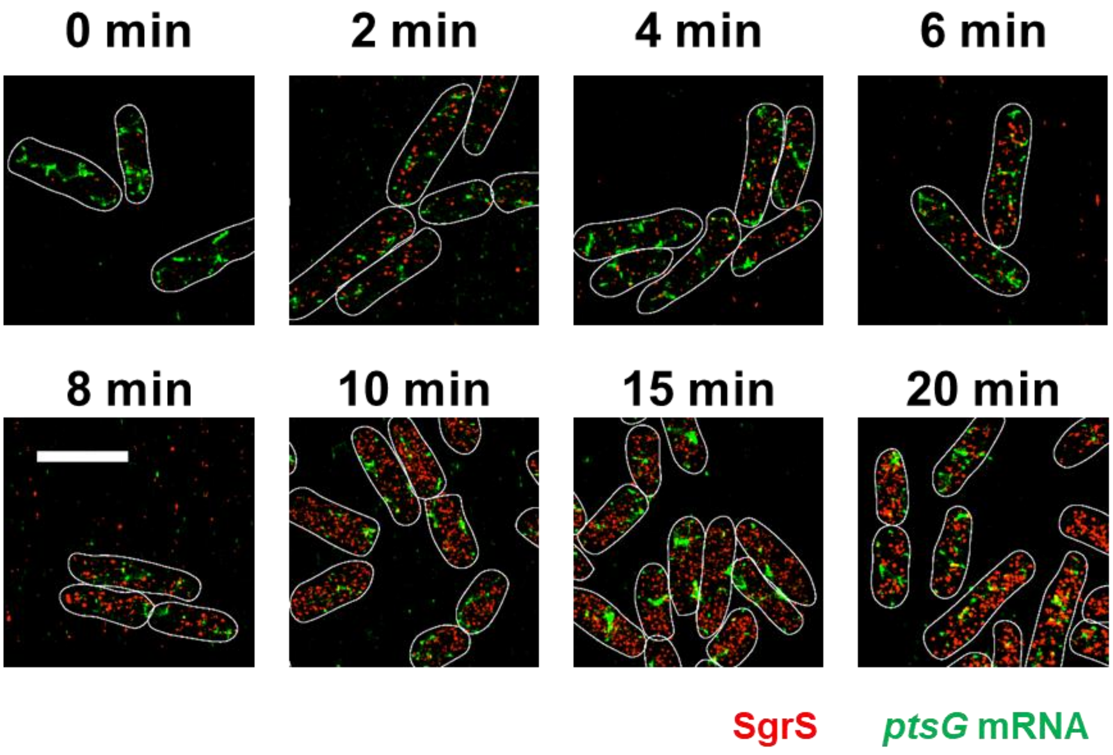


**Supplementary Figure 16. 3D super-resolution images of SgrS (red) and *ptsG* mRNA (green) in the SgrS G178A RNase E mutant strain projected on 2D planes.** The panels show the multi-color images of SgrS G178A RNase E mutant cells before (0 min) and 2, 4, 6, 8, 10, 15, 20 min after αMG (non-metabolizable sugar analog) induction. White lines denote cell boundaries. Scale bar is 2 μm.

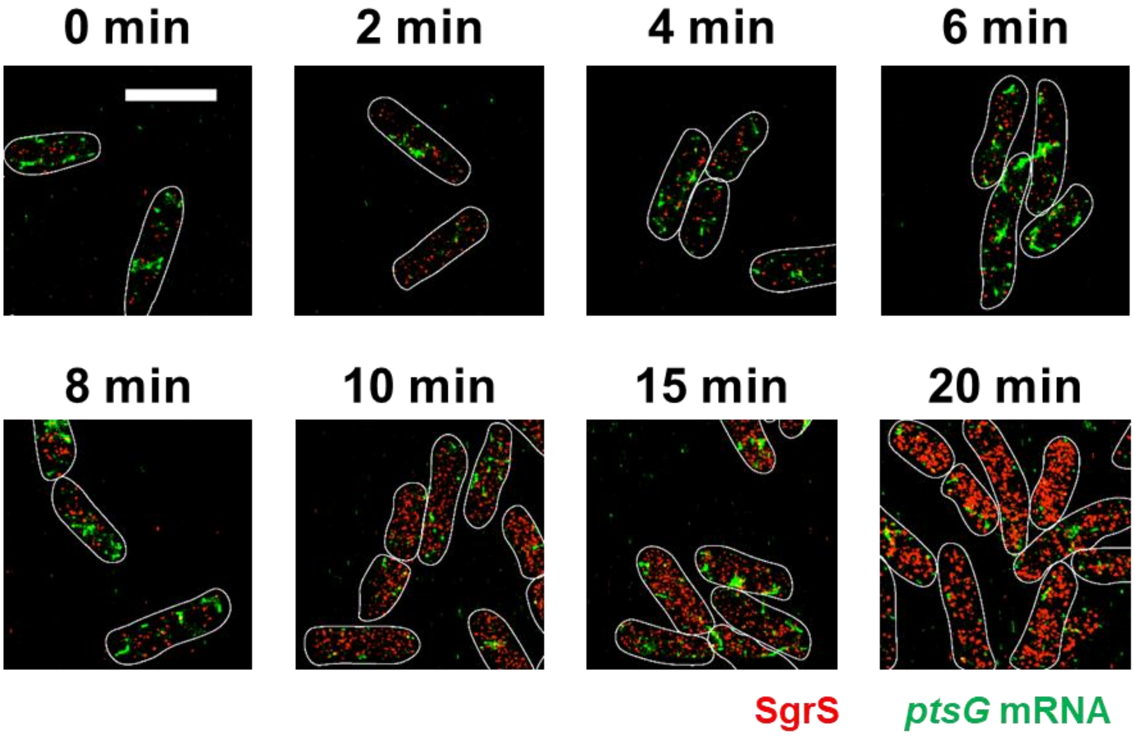


**Supplementary Figure 17. 3D super-resolution images of SgrS (red) and *ptsG* mRNA (green) in the SgrS G178U RNase E mutant strain projected on 2D planes.** The panels show the multi-color images of SgrS G178U RNase E mutant cells before (0 min) and 2, 4, 6, 8, 10, 15, 20 min after αMG (non-metabolizable sugar analog) induction. White lines denote cell boundaries. Scale bar is 2 μm.

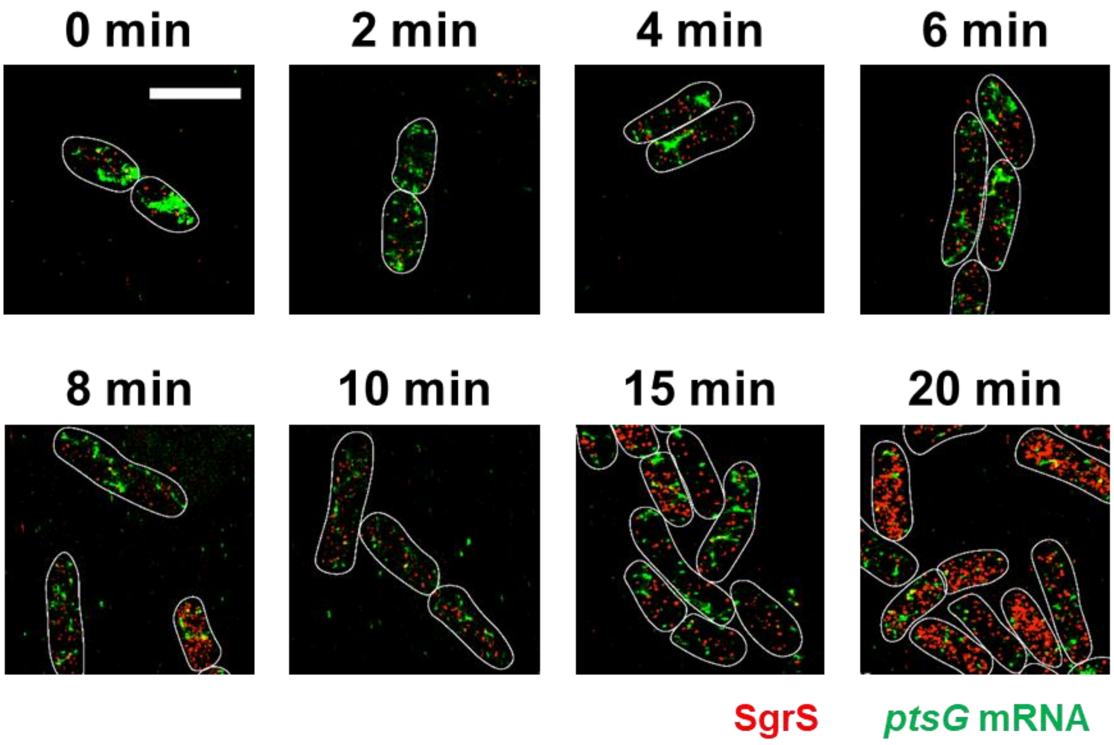


**Supplementary Figure 18. 3D super-resolution images of SgrS (red) and *ptsG* mRNA (green) in the SgrS U181A RNase E mutant strain projected on 2D planes.** The panels show the multi-color images of SgrS U181A RNase E mutant cells before (0 min) and 2, 4, 6, 8, 10, 15, 20 min after αMG (non-metabolizable sugar analog) induction. White lines denote cell boundaries. Scale bar is 2 μm.

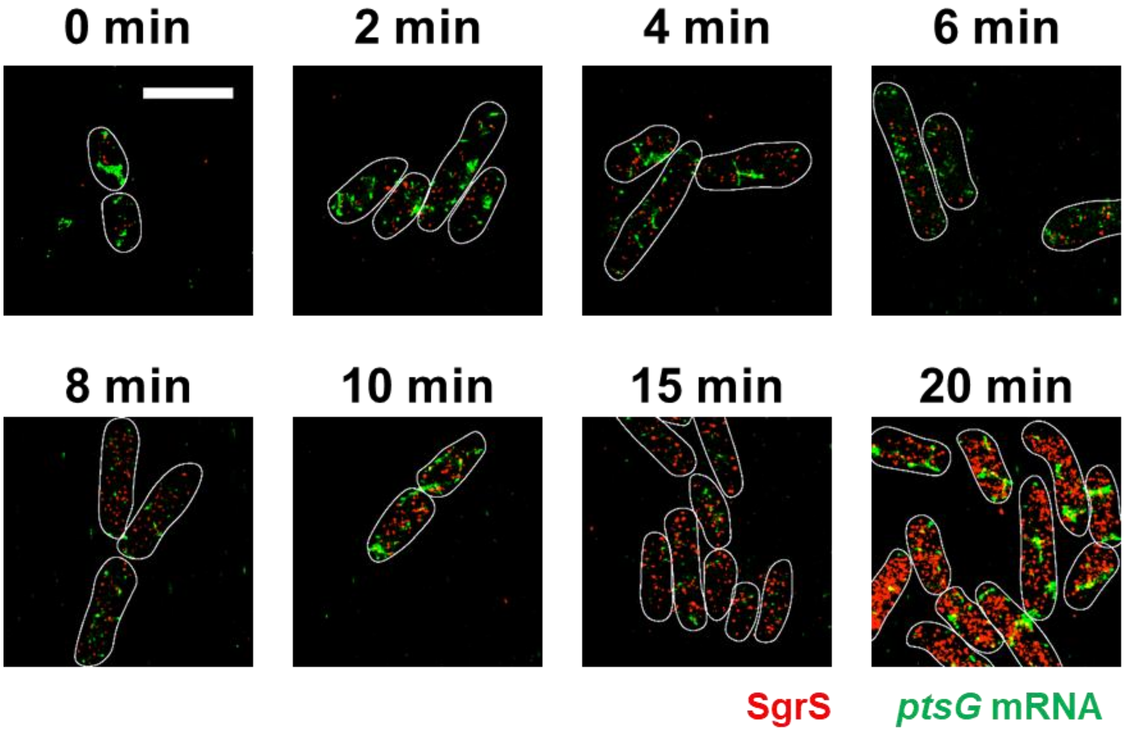


**Supplementary Figure 19. 3D super-resolution images of SgrS (red) and *ptsG* mRNA (green) in the SgrS U182A RNase E mutant strain projected on 2D planes.** The panels show the multi-color images of SgrS U182A RNase E mutant cells before (0 min) and 2, 4, 6, 8, 10, 15, 20 min after αMG (non-metabolizable sugar analog) induction. White lines denote cell boundaries. Scale bar is 2 μm.

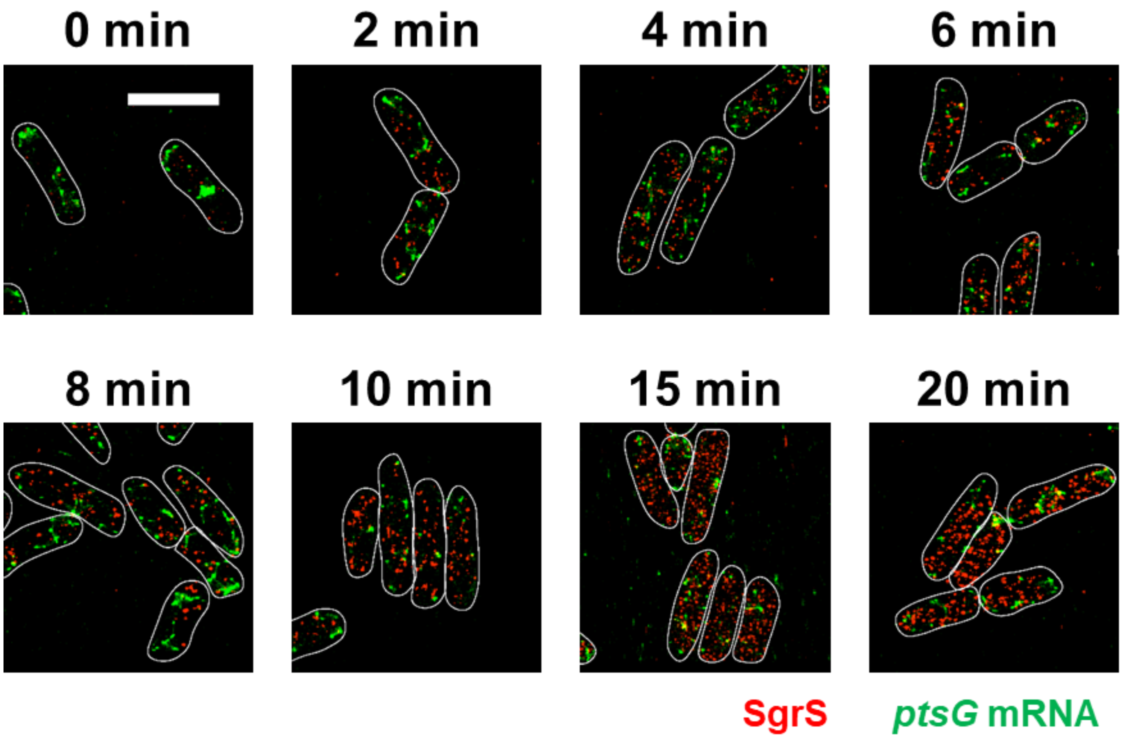


**Supplementary Figure 20. 3D super-resolution images of SgrS (red) and *ptsG* mRNA (green) in the SgrS G184A RNase E mutant strain projected on 2D planes.** The panels show the multi-color images of SgrS G184A RNase E mutant cells before (0 min) and 2, 4, 6, 8, 10, 15, 20 min after αMG (non-metabolizable sugar analog) induction. White lines denote cell boundaries. Scale bar is 2 μm.

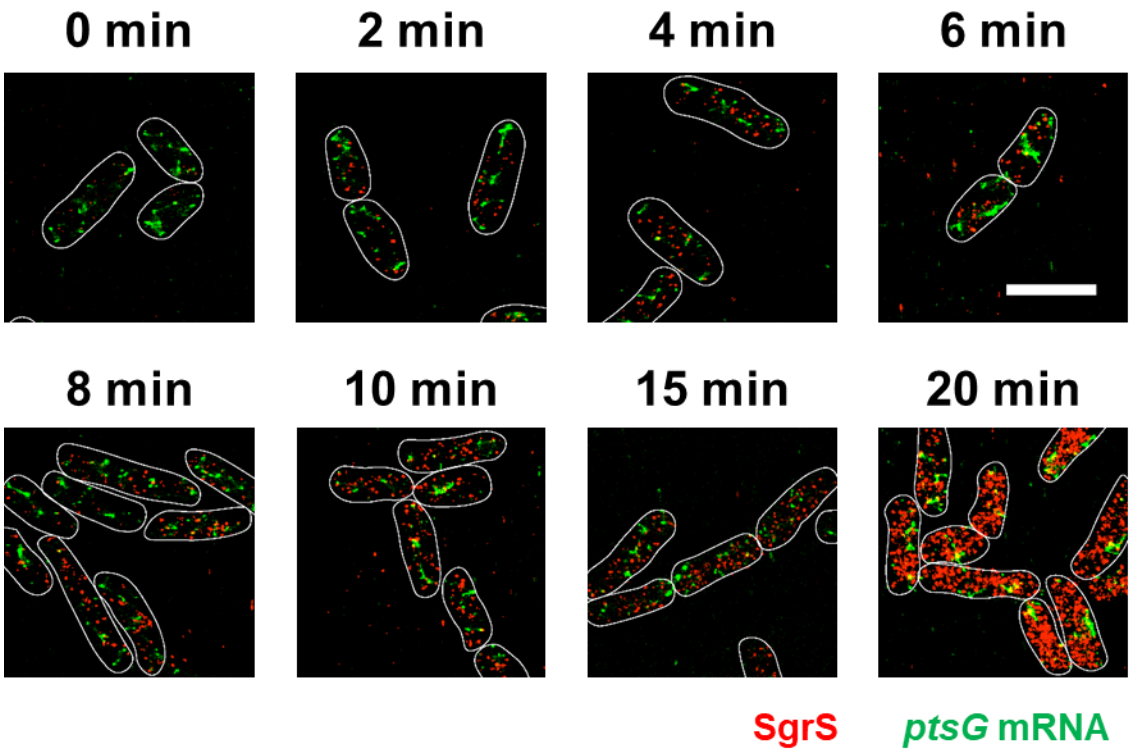


**Supplementary Figure 21. 3D super-resolution images of SgrS (red) and *ptsG* mRNA (green) in the SgrS G184A-C195U RNase E mutant strain projected on 2D planes.** The panels show the multi-color images of SgrS G184A-C195U RNase E mutant cells before (0 min) and 2, 4, 6, 8, 10, 15, 20 min after αMG (non-metabolizable sugar analog) induction. White lines denote cell boundaries. Scale bar is 2 μm.

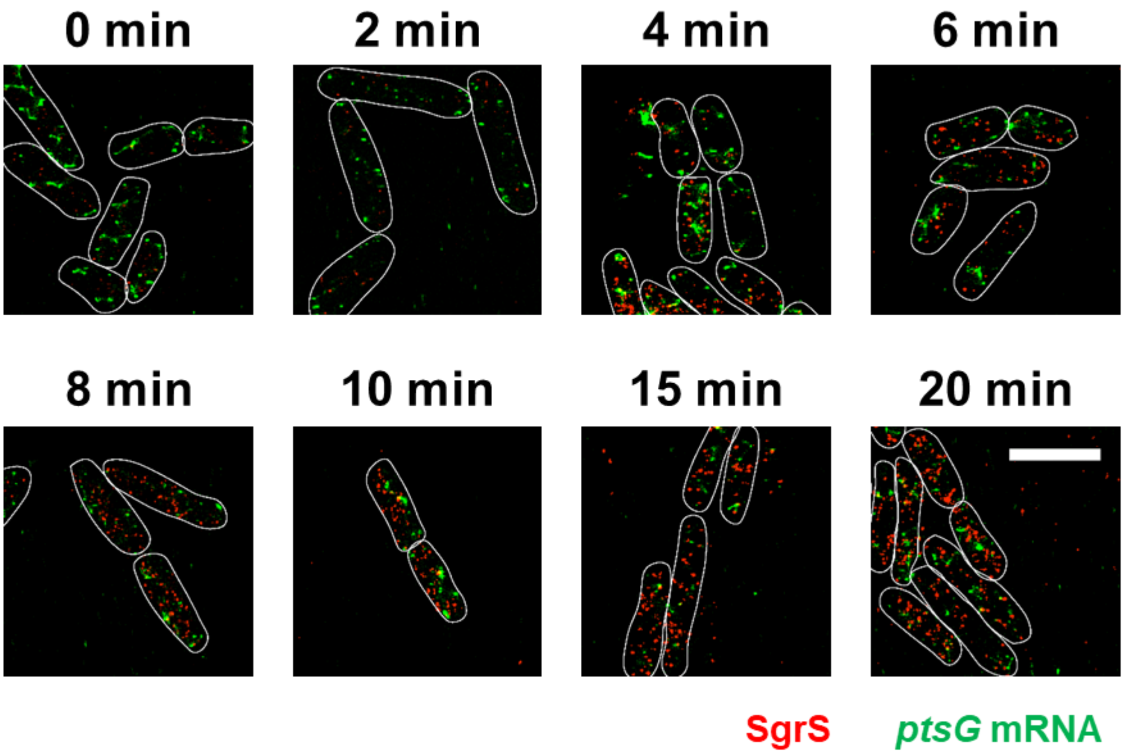


**Supplementary Figure 22. 3D super-resolution images of SgrS (red) and *ptsG* mRNA (green) in the SgrS G215A RNase E mutant strain projected on 2D planes.** The panels show the multi-color images of SgrS G215A RNase E mutant cells before (0 min) and 2, 4, 6, 8, 10, 15, 20 min after αMG (non-metabolizable sugar analog) induction. White lines denote cell boundaries. Scale bar is 2 μm.

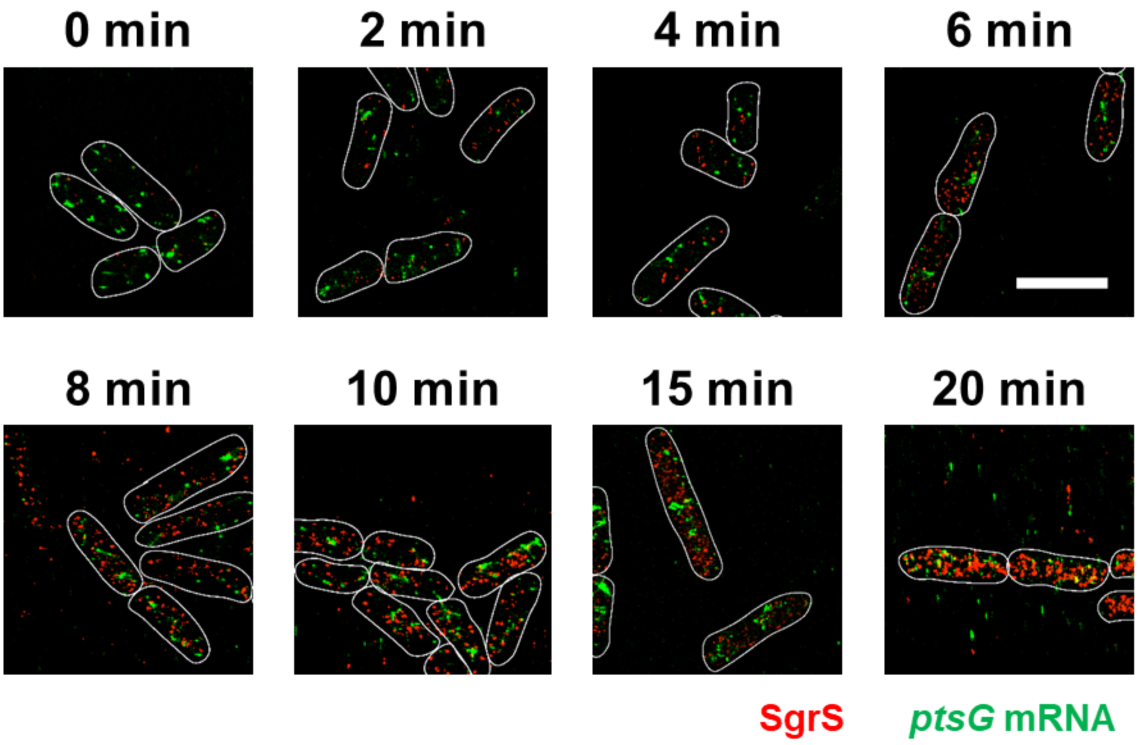


**Supplementary Figure 23. 3D super-resolution images of SgrS (red) and *ptsG* mRNA (green) in the SgrS U224A RNase E mutant strain projected on 2D planes.** The panels show the multi-color images of SgrS U224A RNase E mutant cells before (0 min) and 2, 4, 6, 8, 10, 15, 20 min after αMG (non-metabolizable sugar analog) induction. White lines denote cell boundaries. Scale bar is 2 μm.

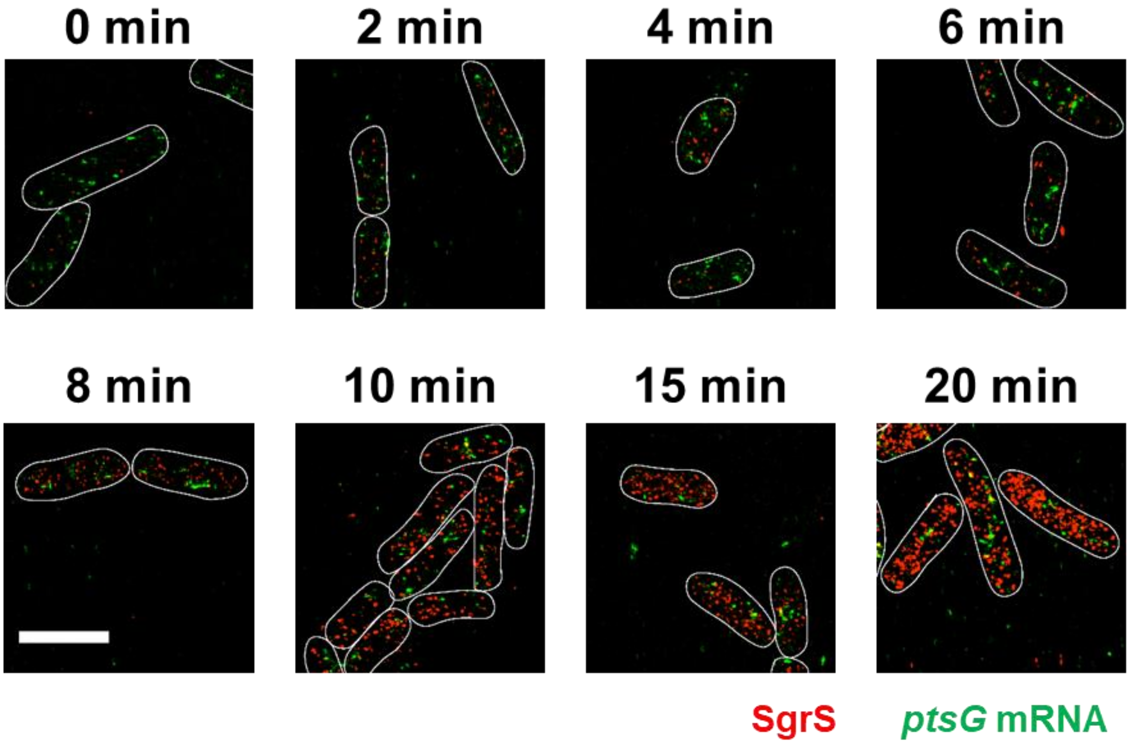


**Supplementary Figure 24. 3D super-resolution images of SgrS (red) and *ptsG* mRNA (green) in the SgrS U224G RNase E mutant strain projected on 2D planes.** The panels show the multi-color images of SgrS U224G RNase E mutant cells before (0 min) and 2, 4, 6, 8, 10, 15, 20 min after αMG (non-metabolizable sugar analog) induction. White lines denote cell boundaries. Scale bar is 2 μm.

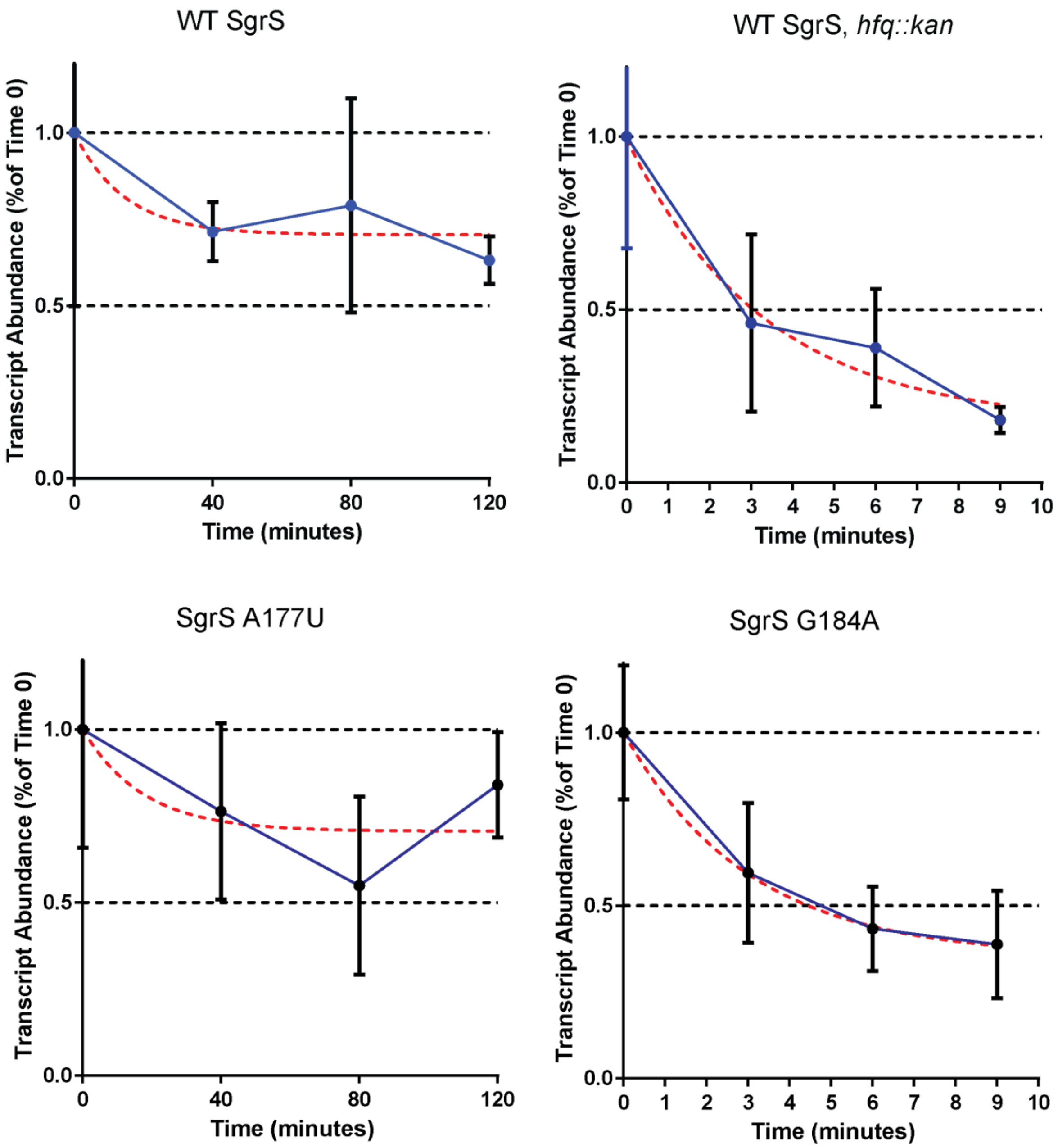


**Supplementary Figure 25. Stability of SgrS transcripts in different strains, (a)** wild-type SgrS, **(b)** Δ*hfq* wild-type strain, **(c)** SgrS A177U strain, **(d)** SgrS G184A strain. Rifampicin was added to the SgrS induced culture to inhibit the synthesis of all cellular transcripts (including SgrS and all its targets). To measure target independent half-life of SgrS, the starting time point was set 5 minutes after the addition of rifampicin. The cells were harvested for total RNA extraction at the indicated time points, and mRNA levels were measured with RT-PCR. The transcript concentrations were normalized to 16S ribosome gene (*rrsA*) transcripts, and were expressed as fractions of abundance in time point 0 samples. The transcript turnover rates were calculated based on the non-linear fit one phase exponential decay curves using GraphPad software (red dotted lines).

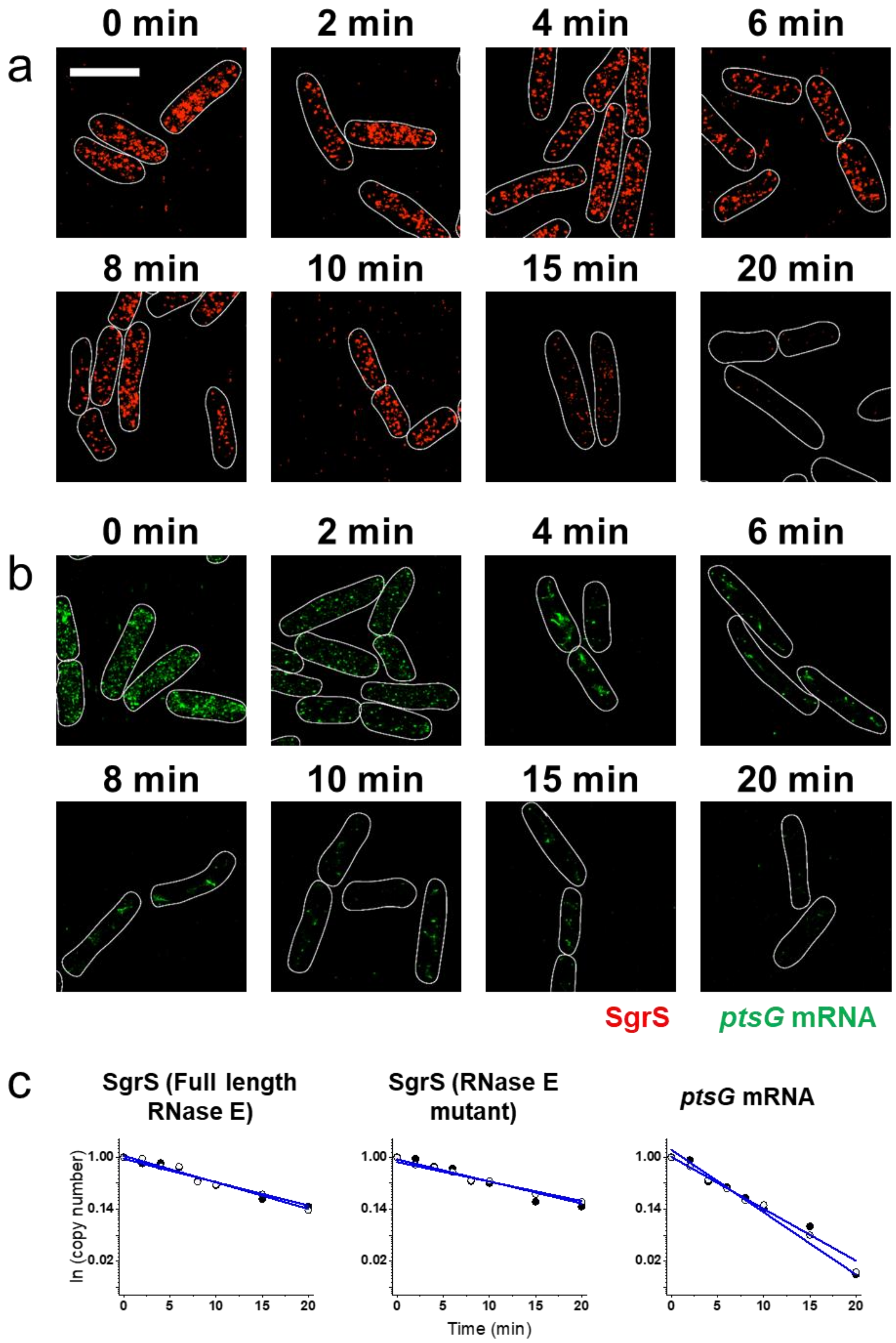
a
0 min
2 min
4 min
6 min
8 min
10 min
15 min
20 min
b
0 min
2 min
4 min
6 min
8 min
10 min
15 min
20 min
SgrS
ptsG mRNA
c
SgrS (Full length RNase E)
SgrS (RNase E mutant)
ptsG mRNA
ln (copy number)
1.00
0.14
0.02
0
5
10
15
20
Time (min)

224

**Supplementary Figure 26. RNA lifetime measurements for the wild-type strain. (a)** SgrS degradation in the wild-type strain. **(b)** *ptsG* mRNA degradation in the wild-type strain. **(c)** Calculation of RNA lifetime. Filled and open circles are two independent measurements with mean value from ~80 cells in each case. The copy numbers have been normalized to time t=0 in each case. The degradation rates calculated from the lifetimes are shown in Fig. 6a, Supplementary Fig. 37 and Supplementary Table 3. Scale bar is 2 μm.

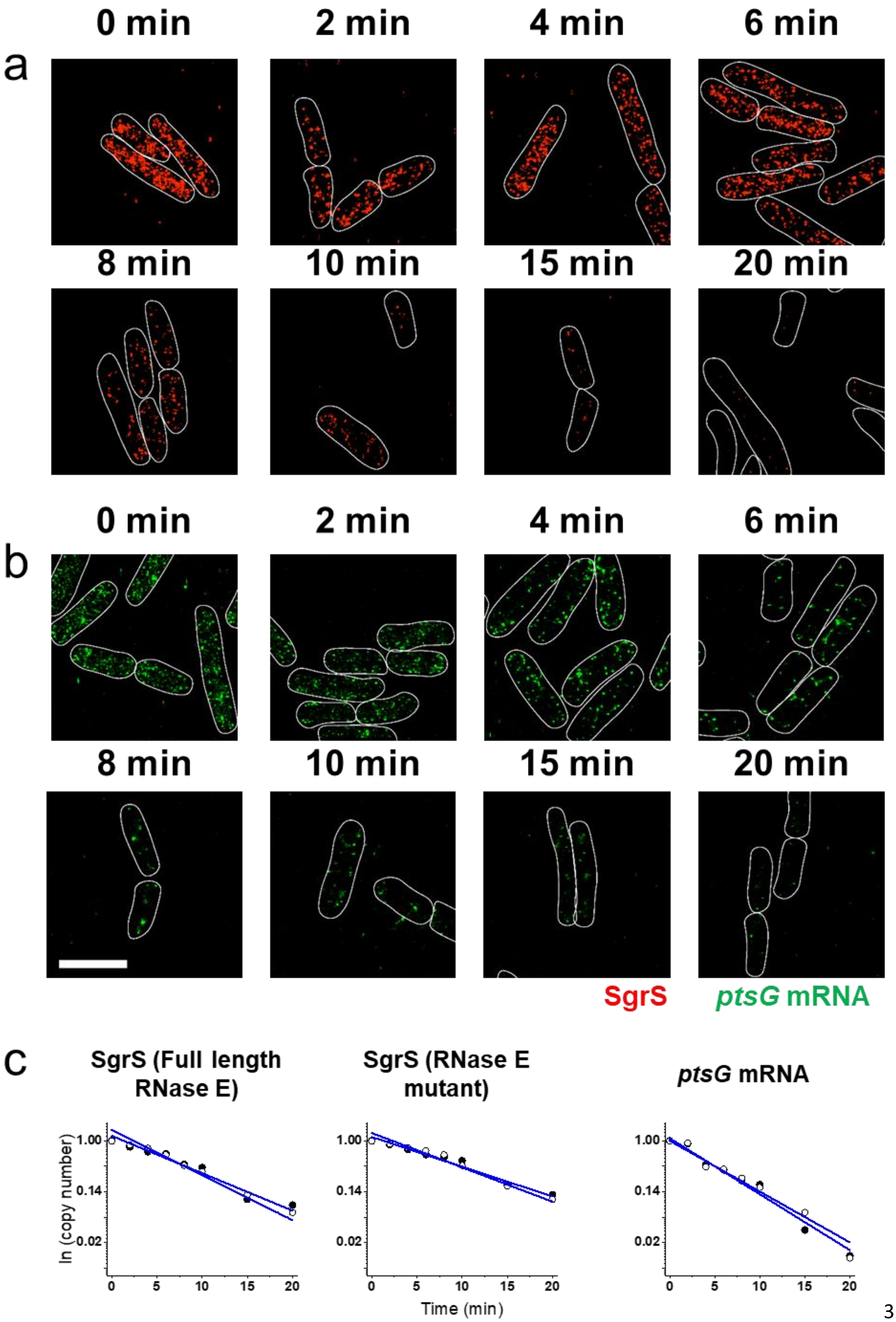
a
0 min
2 min
4 min
6 min
8 min
10 min
15 min
20 min
b
0 min
2 min
4 min
6 min
8 min
10 min
15 min
20 min
SgrS
ptsG mRNA
c
SgrS (Full length RNase E)
SgrS (RNase E mutant)
ptsG mRNA
ln (copy number)
1.00
0.14
0.02
0
5
10
15
20
Time (min)

**Supplementary Figure 27. RNA lifetime measurements for the A177U strain. (a)** SgrS degradation in the A177U strain. **(b)** *ptsG* mRNA degradation in the A177U strain. **(c)** Calculation of RNA lifetime. Filled and open circles are two independent measurements with mean value from ~80 cells in each case. The copy numbers have been normalized to time t=0 in each case. The degradation rates calculated from the lifetimes are shown in Fig. 6a, Supplementary Fig. 37 and Supplementary Table 3. Scale bar is 2 μm.

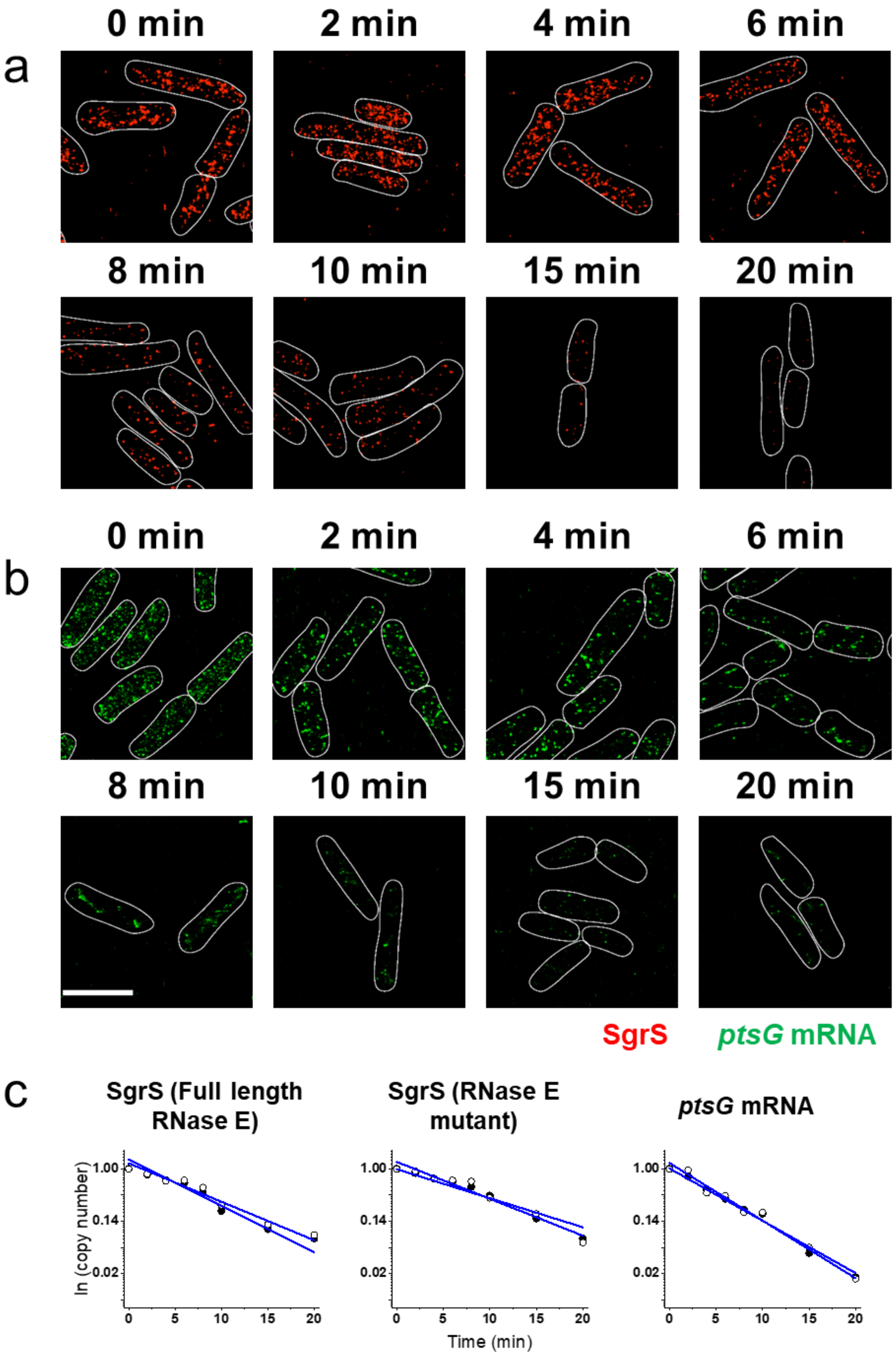

a
0 min
2 min
4 min
6 min
8 min
10 min
15 min
20 min
b
0 min
2 min
4 min
6 min
8 min
10 min
15 min
20 min
SgrS
ptsG mRNA
c
SgrS (Full length RNase E)
SgrS (RNase E mutant)
ptsG mRNA
ln (copy number)
1.00
0.14
0.02
0
5
10
15
20
Time (min)

**Supplementary Figure 28. RNA lifetime measurements for the G178A strain. (a)** SgrS degradation in the G178A strain. **(b)** *ptsG* mRNA degradation in the G178A strain. **(c)** Calculation of RNA lifetime. Filled and open circles are two independent measurements with mean value from ~80 cells in each case. The copy numbers have been normalized to time t=0 in each case. The degradation rates calculated from the lifetimes are shown in Fig. 6a, Supplementary Fig. 37 and Supplementary Table 3. Scale bar is 2 μm.

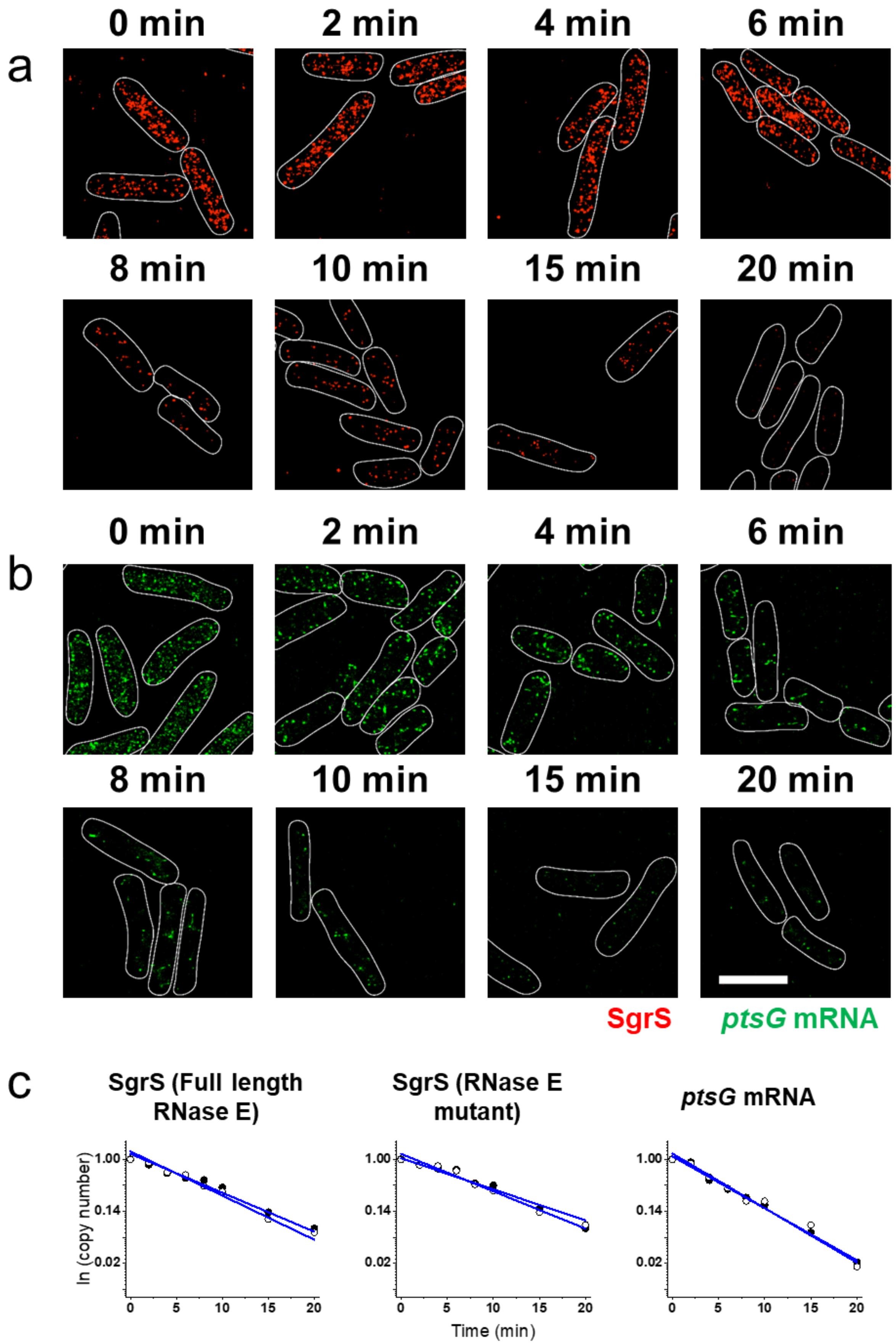
a
0 min
2 min
4 min
6 min
8 min
10 min
15 min
20 min
b
0 min
2 min
4 min
6 min
8 min
10 min
15 min
20 min
SgrS
ptsG mRNA
c
SgrS (Full length RNase E)
SgrS (RNase E mutant)
ptsG mRNA
ln (copy number)
1.00
0.14
0.02
0
5
10
15
20
Time (min)

**Supplementary Figure 29. RNA lifetime measurements for the G178U strain. (a)** SgrS degradation in the G178U strain. **(b)** *ptsG* mRNA degradation in the G178U strain. **(c)** Calculation of RNA lifetime. Filled and open circles are two independent measurements with mean value from ~80 cells in each case. The copy numbers have been normalized to time t=0 in each case. The degradation rates calculated from the lifetimes are shown in Fig. 6a, Supplementary Fig. 37 and Supplementary Table 3. Scale bar is 2 μm.

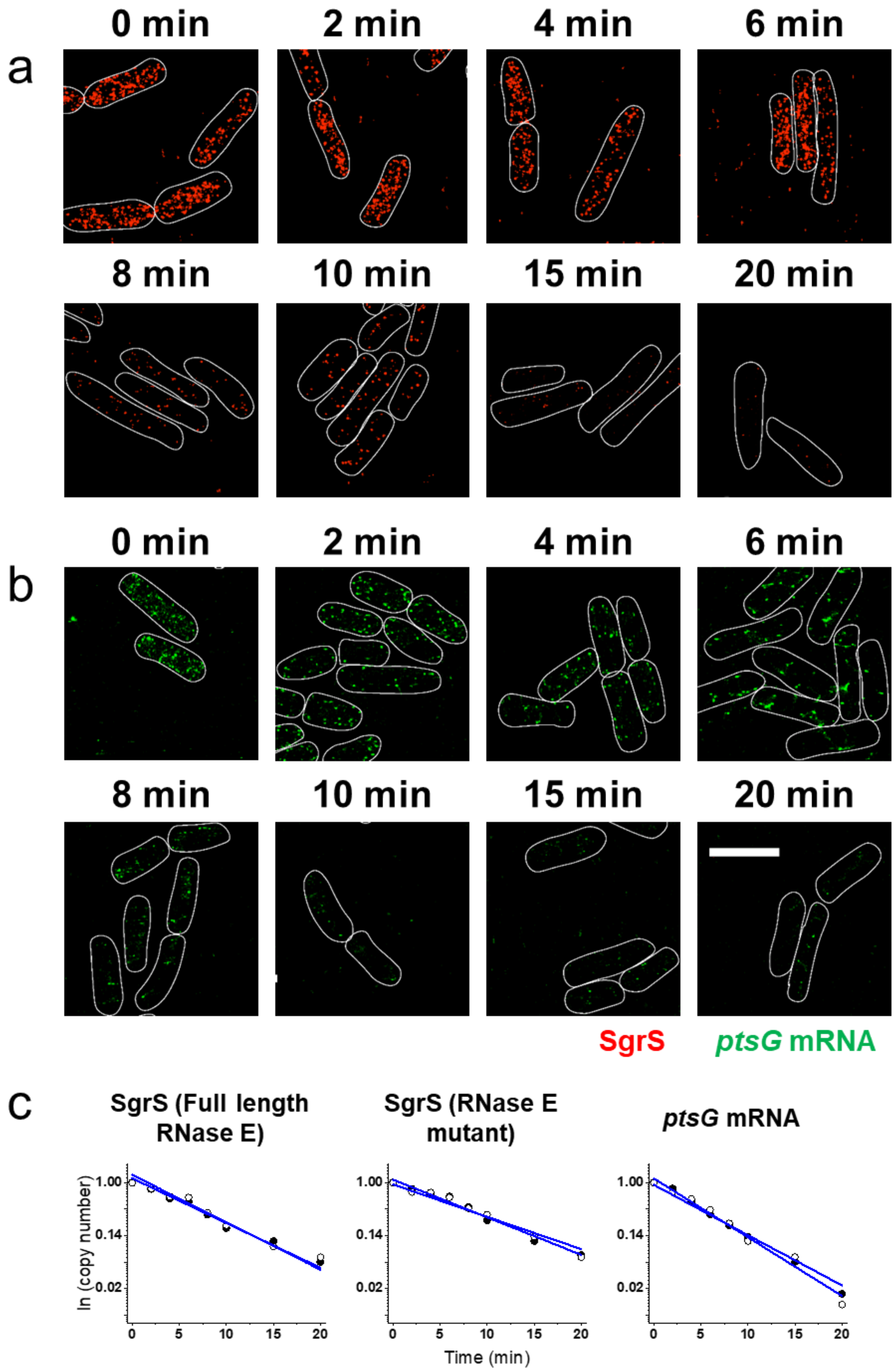
0 min
2 min
4 min
6 min
a
8 min
10 min
15 min
20 min
0 min
2 min
4 min
6 min
b
8 min
10 min
15 min
20 min
SgrS
ptsG mRNA
c
SgrS (Full length RNase E)
SgrS (RNase E mutant)
ptsG mRNA
ln (copy number)
1.00
0.14
0.02
0
5
10
15
20
Time (min)

**Supplementary Figure 30. RNA lifetime measurements for the U181A strain. (a)** SgrS degradation in the U181A strain. **(b)** *ptsG* mRNA degradation in the U181A strain. **(c)** Calculation of RNA lifetime. Filled and open circles are two independent measurements with mean value from ~80 cells in each case. The copy numbers have been normalized to time t=0 in each case. The degradation rates calculated from the lifetimes are shown in Fig. 6a, Supplementary Fig. 37 and Supplementary Table 3. Scale bar is 2 μm.

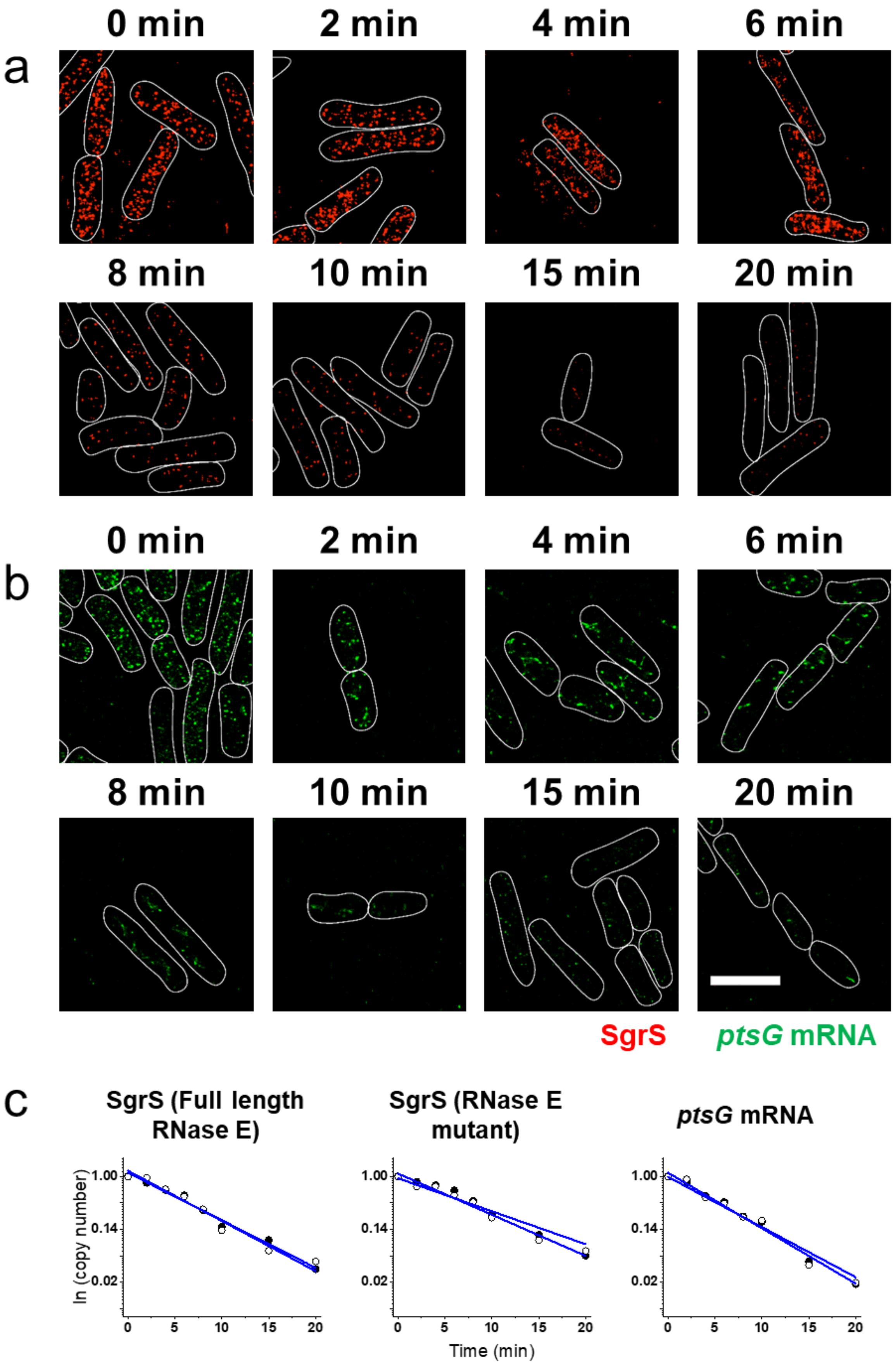
0 min
2 min
4 min
6 min
a
8 min
10 min
15 min
20 min
0 min
2 min
4 min
6 min
b
8 min
10 min
15 min
20 min
SgrS
ptsG mRNA
c
SgrS (Full length RNase E)
SgrS (RNase E mutant)
ptsG mRNA
ln (copy number)
1.00
0.14
0.02
0
5
10
15
20
Time (min)

**Supplementary Figure 31. RNA lifetime measurements for the U182A strain. (a)** SgrS degradation in the U182A strain. **(b)** *ptsG* mRNA degradation in the U182A strain. **(c)** Calculation of RNA lifetime. Filled and open circles are two independent measurements with mean value from ~80 cells in each case. The copy numbers have been normalized to time t=0 in each case. The degradation rates calculated from the lifetimes are shown in Fig. 6a, Supplementary Fig. 37 and Supplementary Table 3. Scale bar is 2 μm.

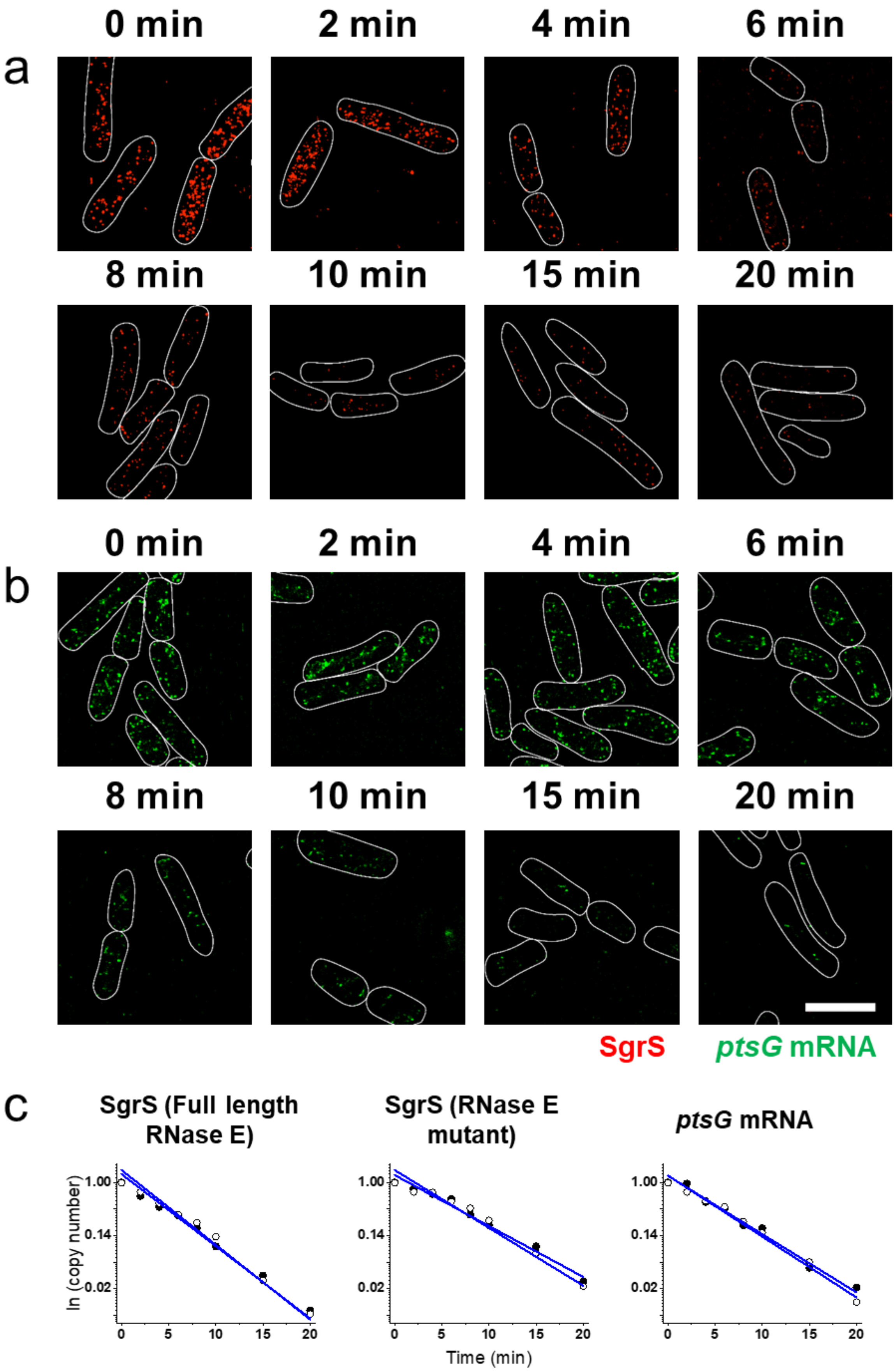

a
0 min
2 min
4 min
6 min
8 min
10 min
15 min
20 min
b
0 min
2 min
4 min
6 min
8 min
10 min
15 min
20 min
SgrS
ptsG mRNA
c
SgrS (Full length RNase E)
SgrS (RNase E mutant)
ptsG mRNA
ln (copy number)
1.00
0.14
0.02
0
5
10
15
20
Time (min)

**Supplementary Figure 32. RNA lifetime measurements for the G184A strain. (a)** SgrS degradation in the G184A strain. **(b)** *ptsG* mRNA degradation in the G184A strain. **(c)** Calculation of RNA lifetime. Filled and open circles are two independent measurements with mean value from ~80 cells in each case. The copy numbers have been normalized to time t=0 in each case. The degradation rates calculated from the lifetimes are shown in Fig. 6a, Supplementary Fig. 37 and Supplementary Table 3. Scale bar is 2 μm.

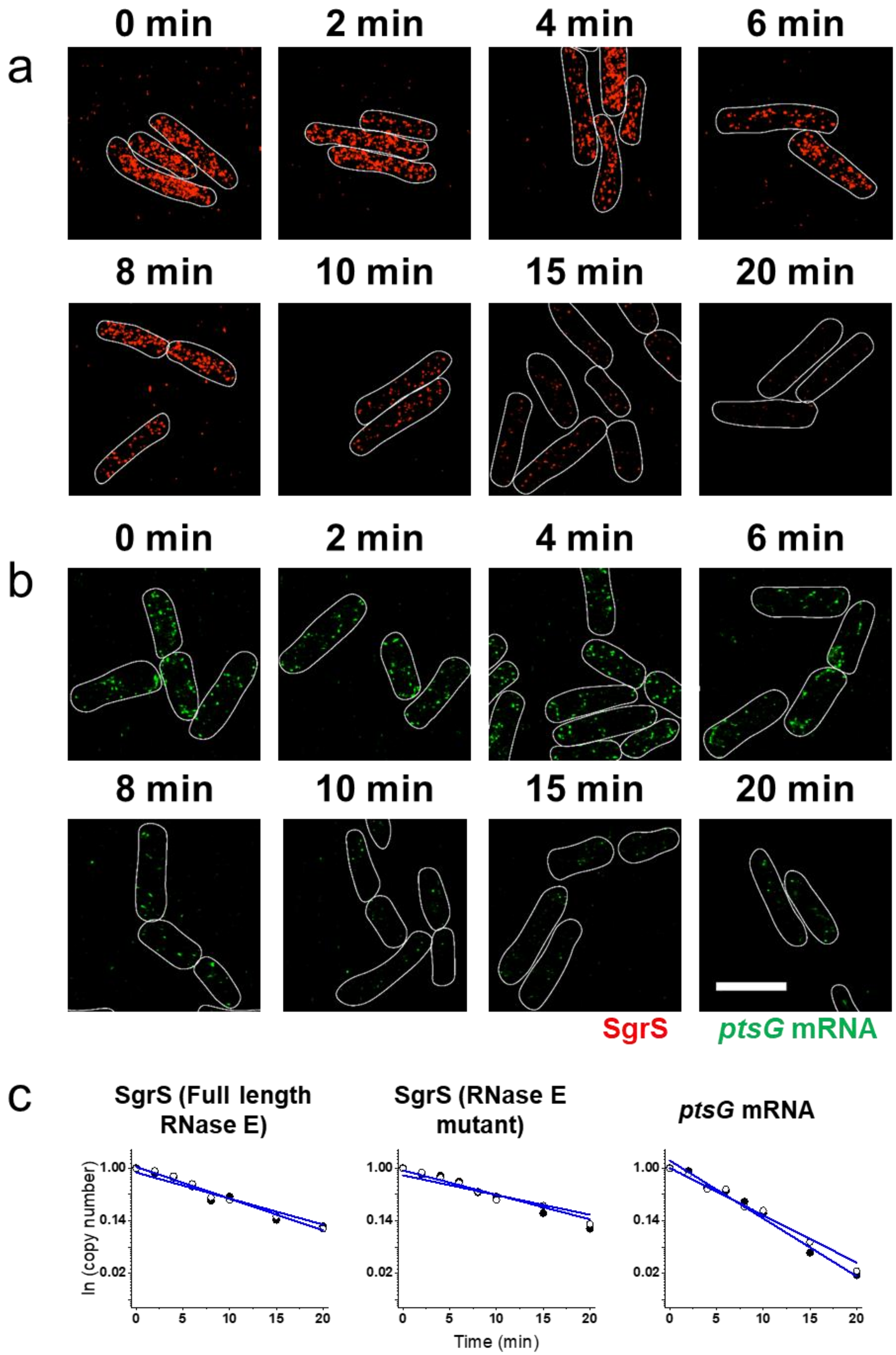
0 min
2 min
4 min
6 min
a
8 min
10 min
15 min
20 min
0 min
2 min
4 min
6 min
b
8 min
10 min
15 min
20 min
SgrS
ptsG mRNA
c
SgrS (Full length RNase E)
SgrS (RNase E mutant)
ptsG mRNA
ln (copy number)
1.00
0.14
0.02
0
5
10
15
20
Time (min)

287

**Supplementary Figure 33. RNA lifetime measurements for the G184A-C195U strain. (a)** SgrS degradation in the G184A-C195U strain. **(b)** *ptsG* mRNA degradation in the G184A-C195U strain. **(c)** Calculation of RNA lifetime. Filled and open circles are two independent measurements with mean value from ~80 cells in each case. The copy numbers have been normalized to time t=0 in each case. The degradation rates calculated from the lifetimes are shown in Fig. 6a, Supplementary Fig. 37 and Supplementary Table 3. Scale bar is 2 μm.

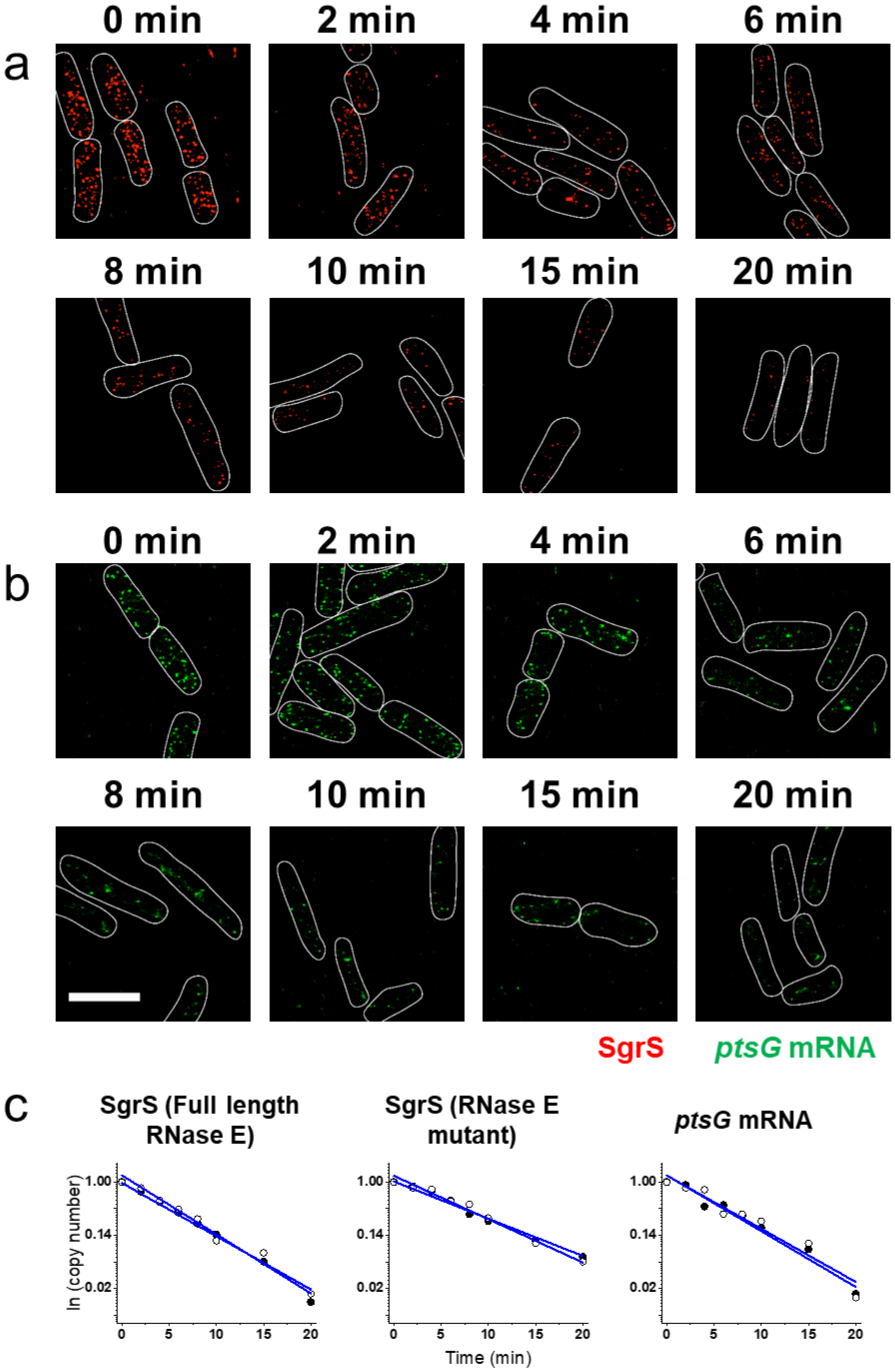
a
0 min
2 min
4 min
6 min
8 min
10 min
15 min
20 min
b
0 min
2 min
4 min
6 min
8 min
10 min
15 min
20 min
SgrS
ptsG mRNA
c
SgrS (Full length RNase E)
SgrS (RNase E mutant)
ptsG mRNA
ln (copy number)
1.00
0.14
0.02
0
5
10
15
20
Time (min)

**Supplementary Figure 34. RNA lifetime measurements for the G215A strain. (a)** SgrS degradation in the G215A strain. **(b)** *ptsG* mRNA degradation in the G215A strain. **(c)** Calculation of RNA lifetime. Filled and open circles are two independent measurements with mean value from ~80 cells in each case. The copy numbers have been normalized to time t=0 in each case. The degradation rates calculated from the lifetimes are shown in Fig. 6a, Supplementary Fig. 37 and Supplementary Table 3. Scale bar is 2 μm.

305

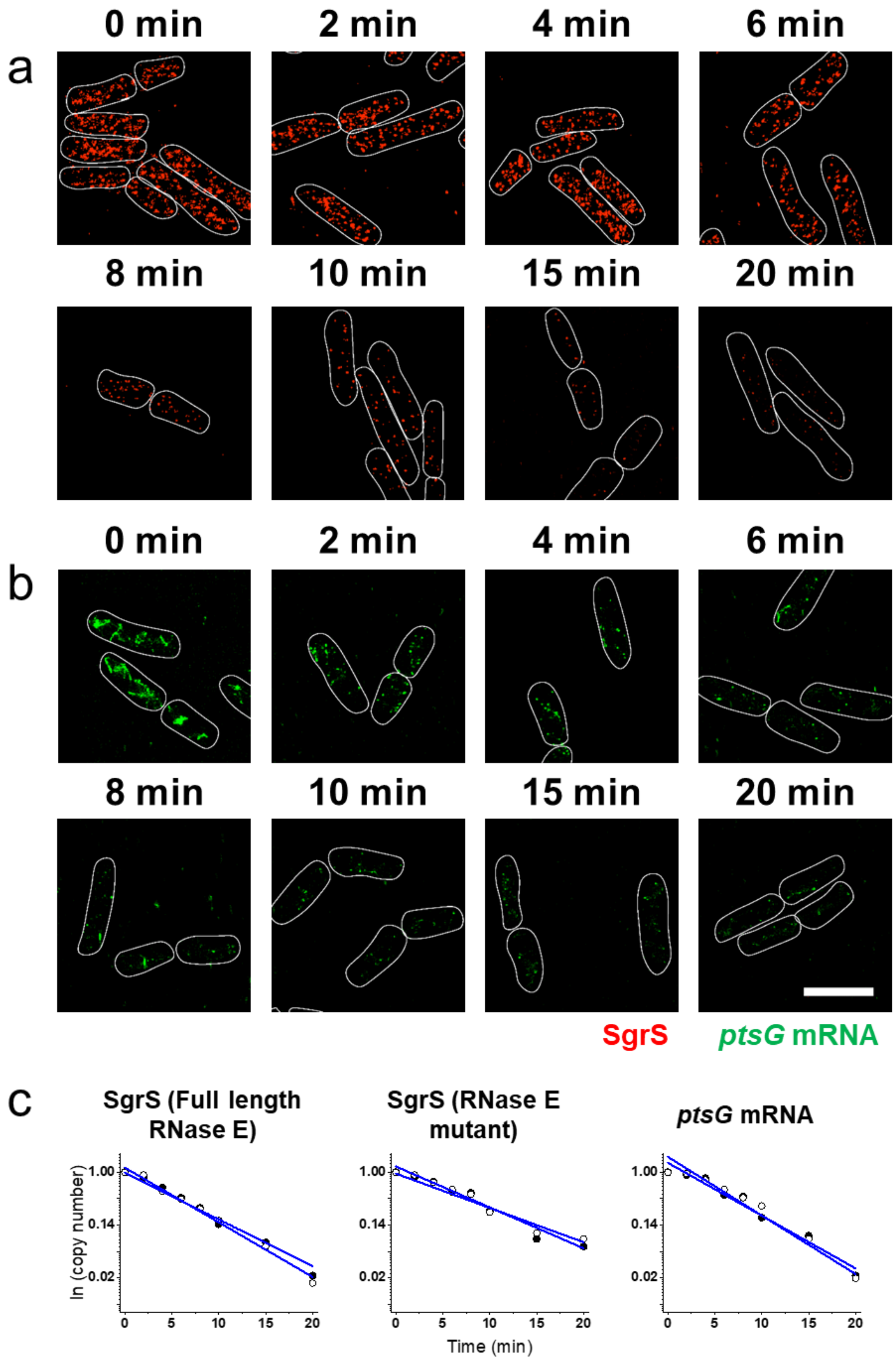
a
0 min
2 min
4 min
6 min
8 min
10 min
15 min
20 min
b
0 min
2 min
4 min
6 min
8 min
10 min
15 min
20 min
SgrS
ptsG mRNA
c
SgrS (Full length RNase E)
SgrS (RNase E mutant)
ptsG mRNA
ln (copy number)
1.00
0.14
0.02
0
5
10
15
20
Time (min)

**Supplementary Figure 35. RNA lifetime measurements for the U224A strain. (a)** SgrS degradation in the U224A strain. **(b)** *ptsG* mRNA degradation in the U224A strain. **(c)** Calculation of RNA lifetime. Filled and open circles are two independent measurements with mean value from ~80 cells in each case. The copy numbers have been normalized to time t=0 in each case. The degradation rates calculated from the lifetimes are shown in Fig. 6a, Supplementary Fig. 37 and Supplementary Table 3. Scale bar is 2 μm.

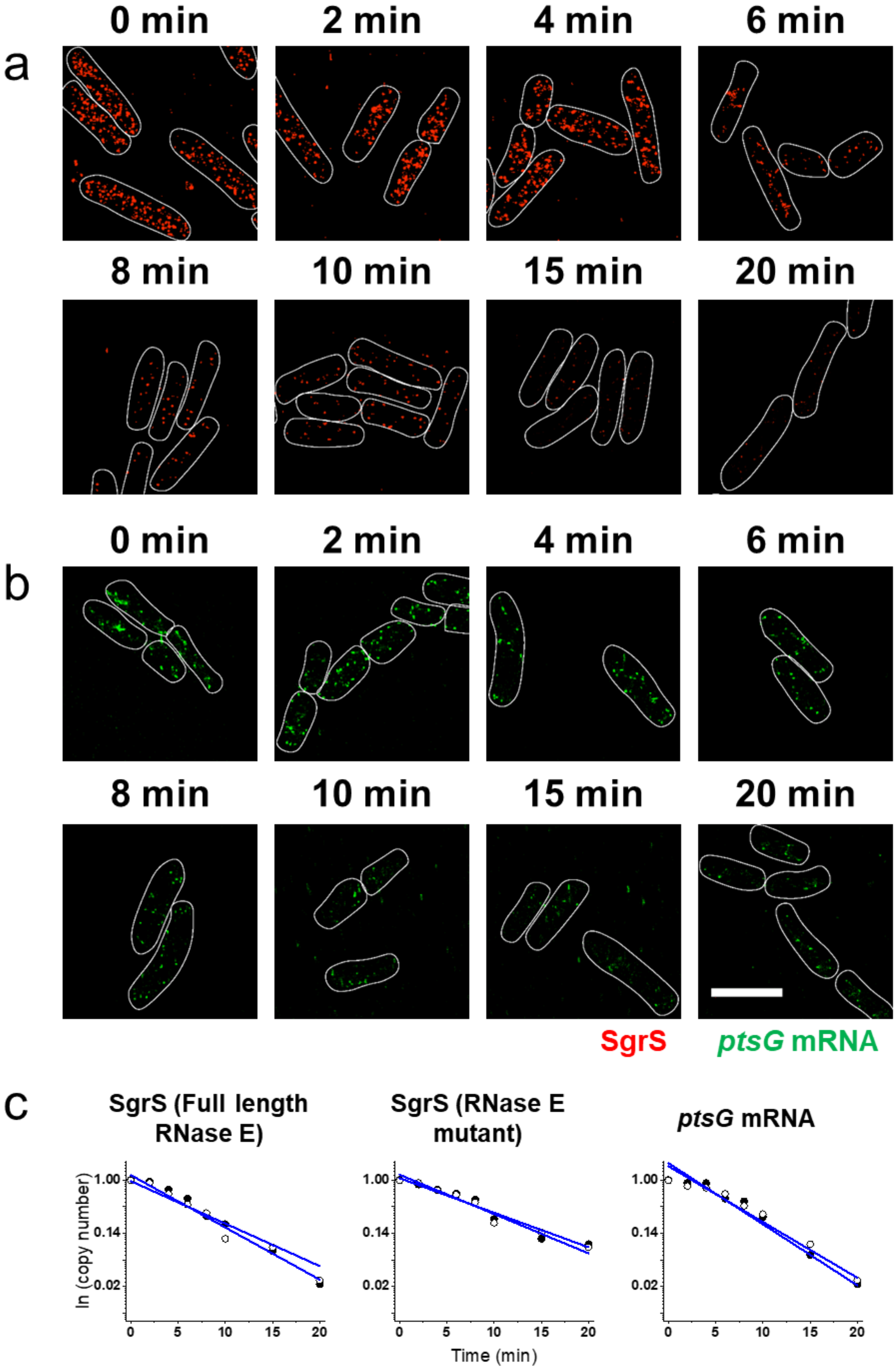

0 min
2 min
4 min
6 min
a
8 min
10 min
15 min
20 min
0 min
2 min
4 min
6 min
b
8 min
10 min
15 min
20 min
SgrS
ptsG mRNA
c
SgrS (Full length RNase E)
SgrS (RNase E mutant)
ptsG mRNA
ln (copy number)
1.00
0.14
0.02
0
5
10
15
20
Time (min)

**Supplementary Figure 36. RNA lifetime measurements for the U224G strain. (a)** SgrS degradation in the U224G strain. **(b)** *ptsG* mRNA degradation in the U224G strain. **(c)** Calculation of RNA lifetime. Filled and open circles are two independent measurements with mean value from ~80 cells in each case. The copy numbers have been normalized to time t=0 in each case. The degradation rates calculated from the lifetimes are shown in Fig. 6a, Supplementary Fig. 37 and Supplementary Table 3. Scale bar is 2 μm.

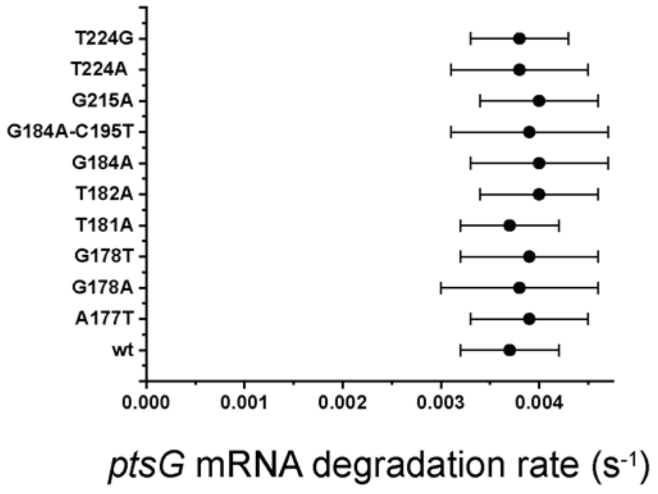


**Supplementary Figure 37. Degradation rates of *ptsG* mRNA in wild-type and the mutant strains.** Degradation rates of *ptsG* mRNA in wild-type and the strains A177U, G178A, G178U, U181A, U182A, G184A, G184A-C195U, G215A, U224A and U224G. Error bars represent standard deviation obtained from two experimental replicates.

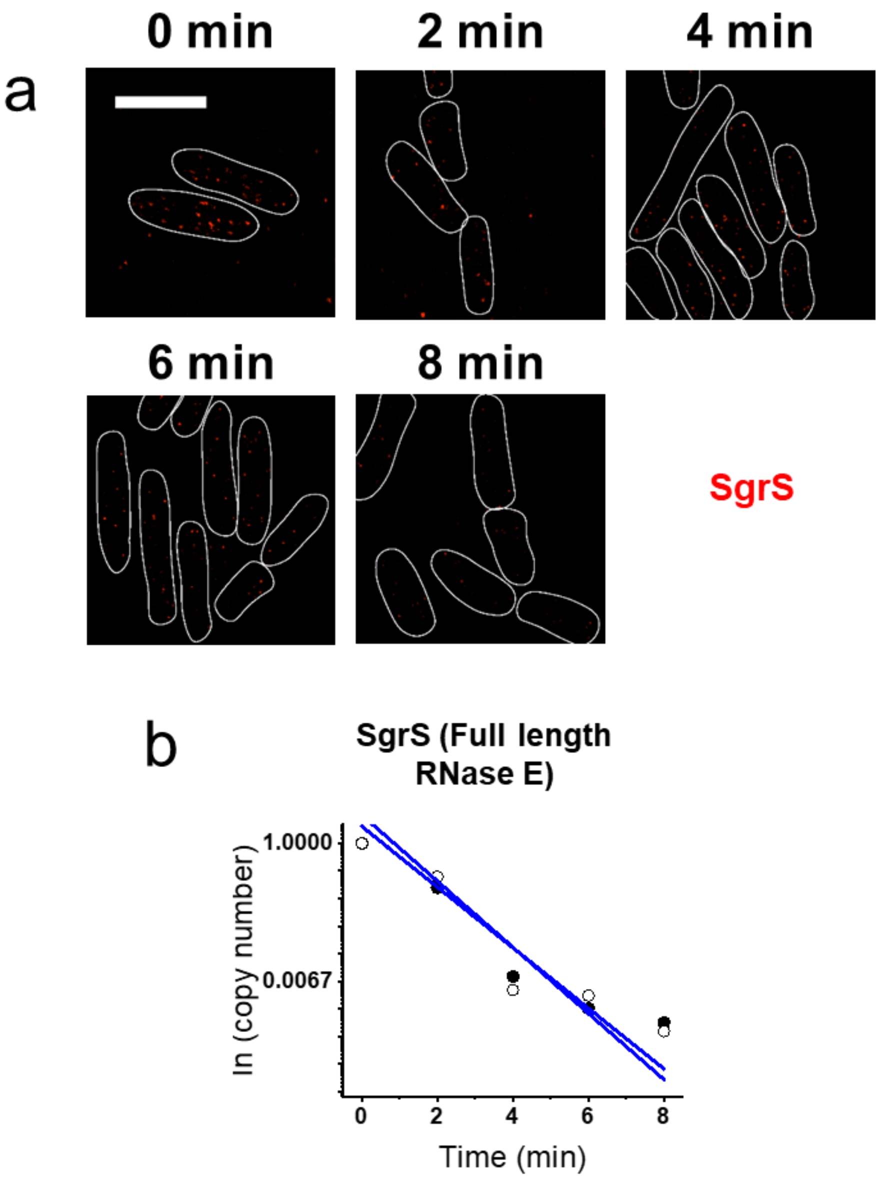


**Supplementary Figure 38. RNA lifetime measurements for the Δ*hfq* wild-type strain. (a)** SgrS degradation in the Δ*hfq* wild-type strain. Scale bar is 2 µm. **(b)** Calculation of RNA lifetime. Filled and open circles are two independent measurements with mean value from ~70 cells in each case. The copy numbers have been normalized to time t=0 in each case. The degradation rates calculated from the lifetimes are shown in Fig 6a.

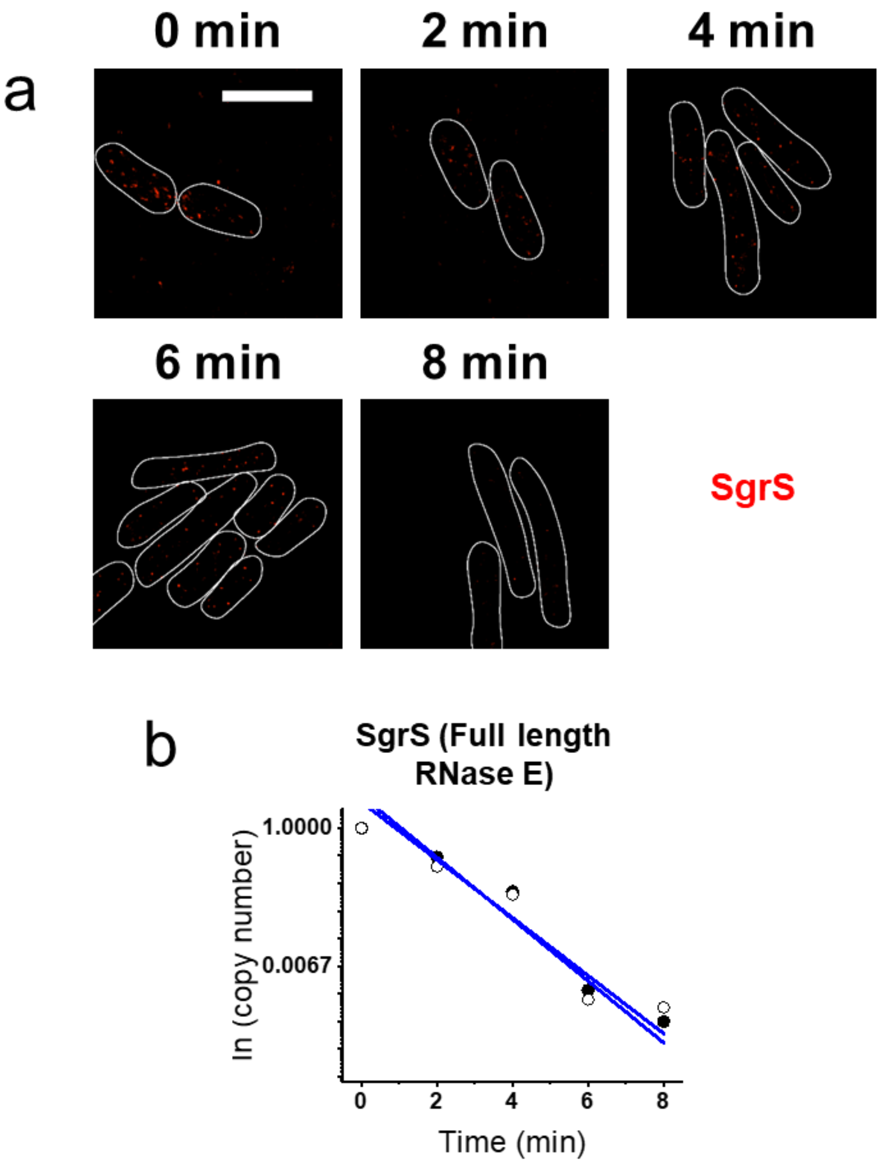


**Supplementary Figure 39. RNA lifetime measurements for the Δ*hfq* A177U strain. (a)** SgrS degradation in the Δ*hfq* A177U strain. Scale bar is 2 μm. **(b)** Calculation of RNA lifetime. Filled and open circles are two independent measurements with mean value from ~70 cells in each case. The copy numbers have been normalized to time t=0 in each case. The degradation rates calculated from the lifetimes are shown in Fig 6a.

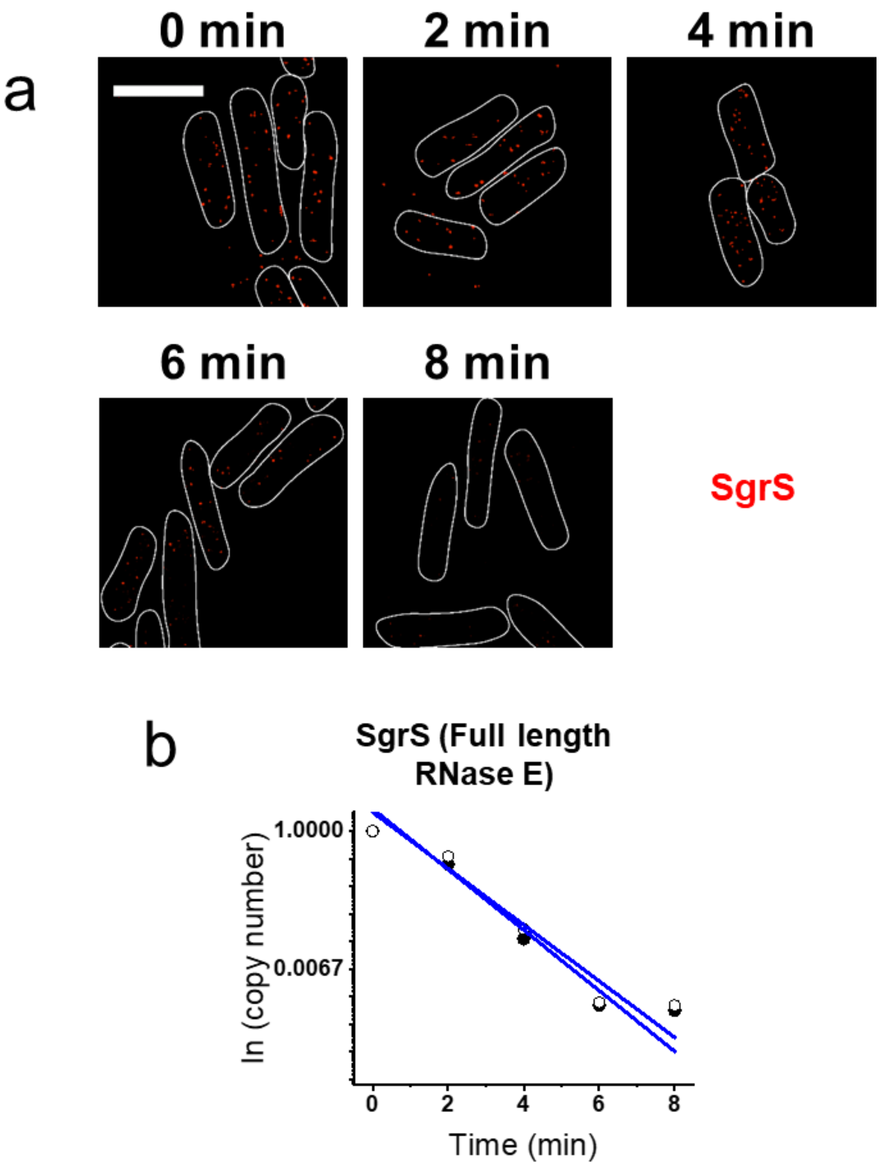


**Supplementary Figure 40. RNA lifetime measurements for the Δ*hfq* G184A strain. (a)** SgrS degradation in the Δ*hfq* G184A strain. Scale bar is 2 μm. **(b)** Calculation of RNA lifetime. Filled and open circles are two independent measurements with mean value from ~70 cells in each case. The copy numbers have been normalized to time t=0 in each case. The degradation rates calculated from the lifetimes are shown in Fig 6a.

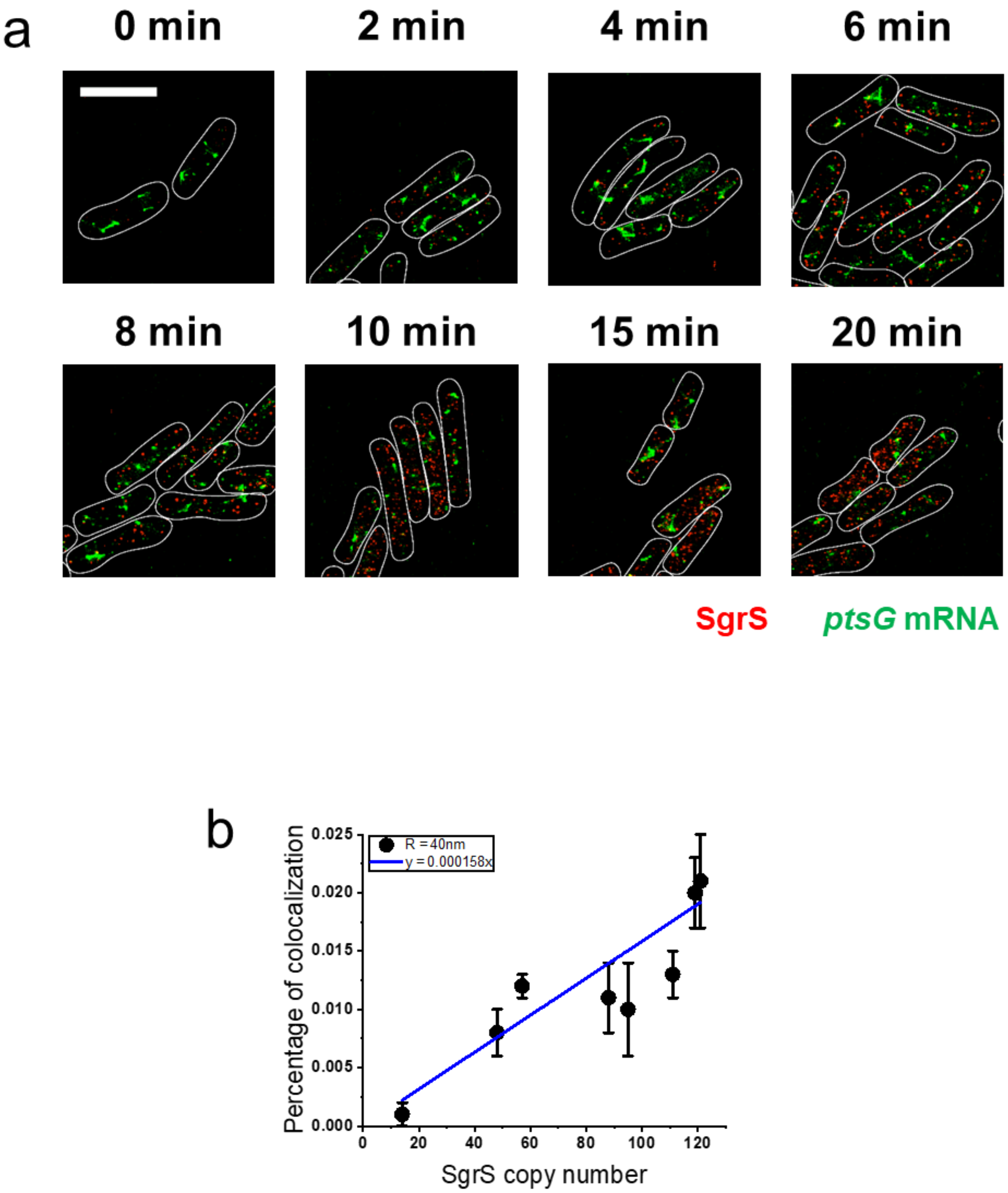


**Supplementary Figure 41. Colocalization analysis for base-pairing mutant strain. (a)** 3D super-resolution images of SgrS (red) and *ptsG* mRNA (green) in the base-pairing mutant strain projected on 2D planes. The panels show the multi-color images of cells before (0 min) and 2, 4,

6, 8, 10, 15, 20 min after αMG (non-metabolizable sugar analog) induction. White lines denote cell boundaries. Scale bar is 2 μm. (b) The percentage of colocalization in the base-pairing mutant strain with R=40 nm as a function of mean SgrS copy number. The plot is fit with a linear function for correction of colocalization by chance. Error bars are standard deviations from 3-6 images.

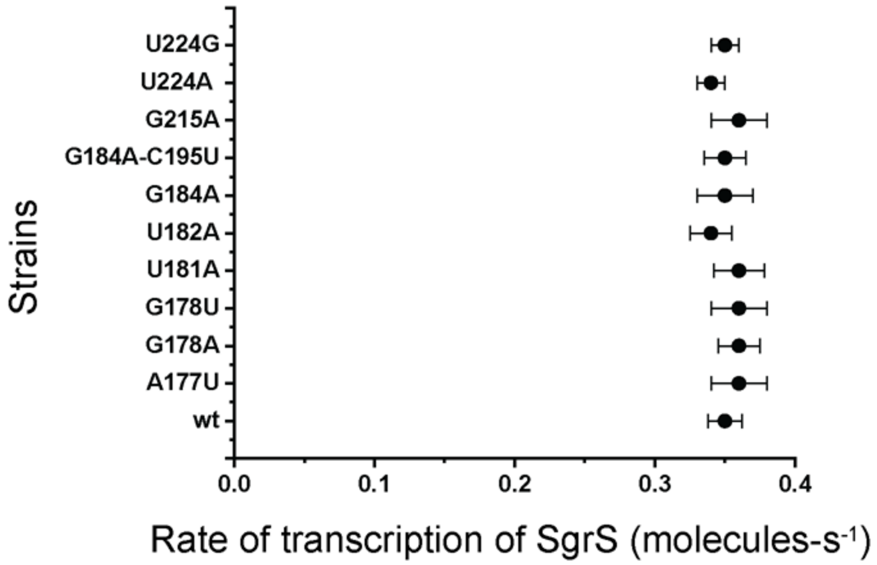


**Supplementary Figure 42. Rates of transcription of SgrS for wild-type and the mutant strains.** Rates of transcription of SgrS for wild-type and the strains A177U, G178A, G178U, U181A, U182A, G184A, G184A-C195U, G215A, U224A and U224G calculated from the time dependent modeling curves of the SgrS, *ptsG* mRNA and SgrS-*ptsG* mRNA complexes. These were determined simultaneously in the wild-type and the RNase E mutants. Error bars represent standard deviation from the independent fitting of two replicates.

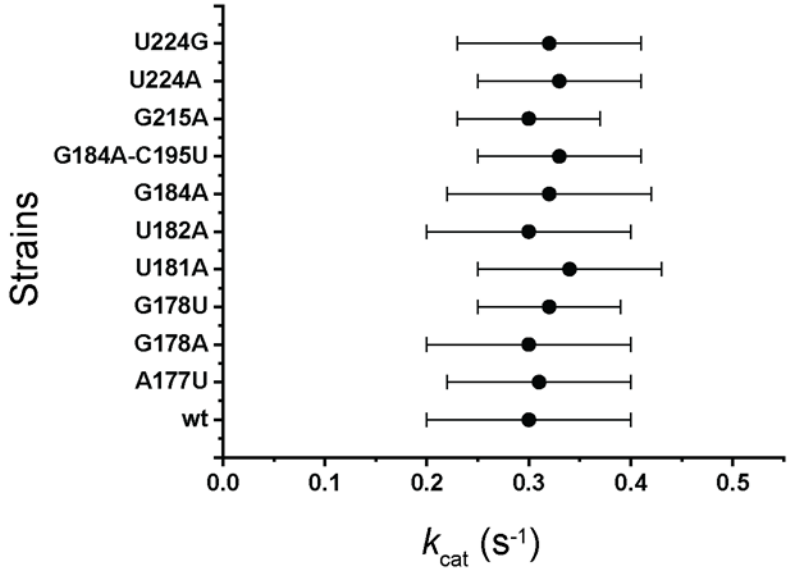


**Supplementary Figure 43. Co-degradation rate of SgrS-*ptsG* mRNA complex for wild-type and the mutants.** $k_{cat}$ measured from the time dependent modeling curves of the SgrS, *ptsG* mRNA and SgrS-*ptsG* mRNA complexes for the wild-type and strains A177U, G178A, G178U, U181A, U182A, G184A, G184A-C195U, G215A, U224A, U224G. These were determined simultaneously in the wild-type and RNase E mutants. Error bars represent standard deviation from the independent fitting on two replicates.

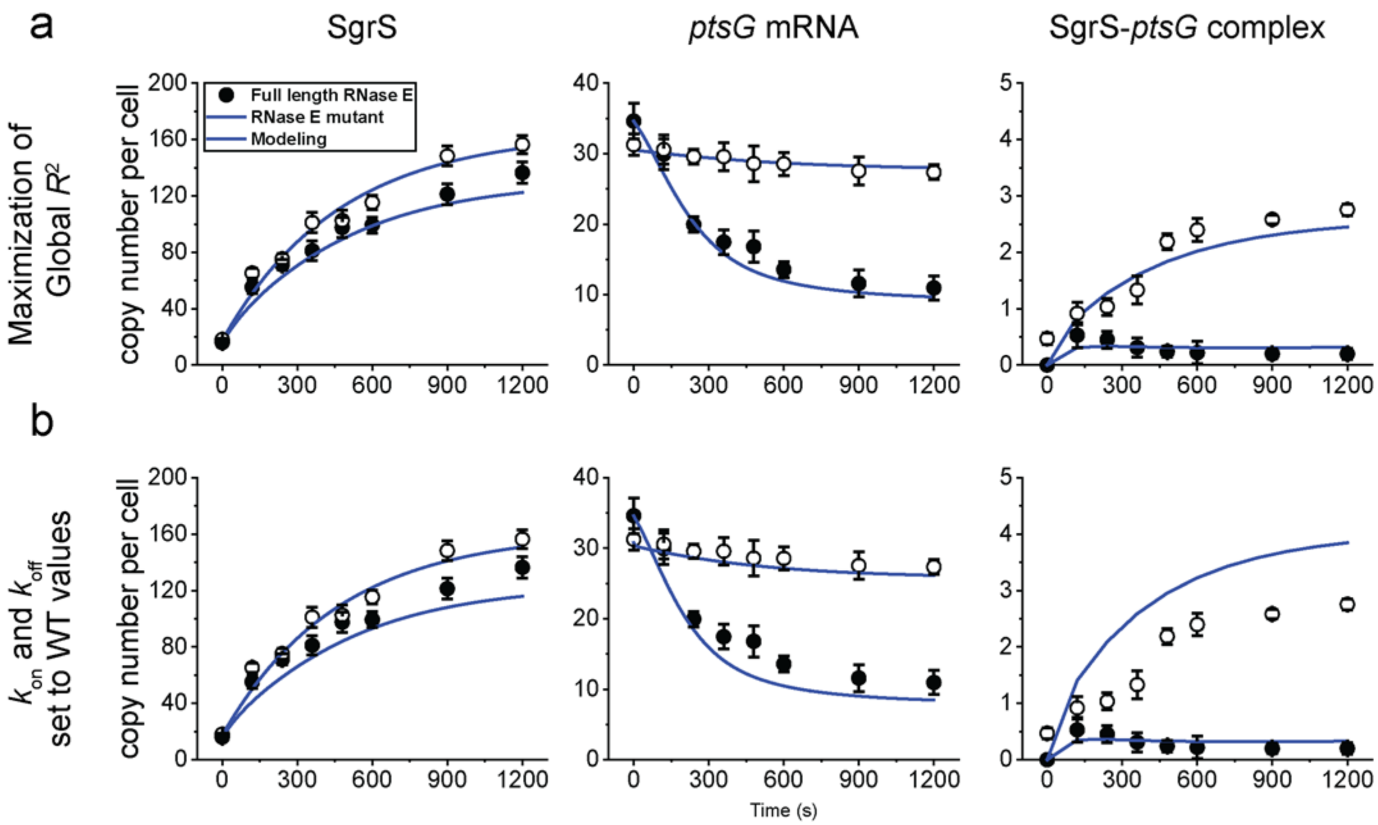


**Supplementary Figure 44. Change in the copy numbers of SgrS and *ptsG* mRNA over time for SgrS A177U mutant strain.** Time course changes from 0 min (0 s) to 20 min (1200 s) after the induction of glucose-phosphate stress using αMG (non-metabolizable sugar analog) and the corresponding modeling curves for SgrS, *ptsG* mRNA and SgrS-*ptsG* complexes for SgrS A177U strain by **(a)** maximization of global $R^2$ and **(b)** setting $k_{on}$ and $k_{off}$ to WT values. The average copy numbers per cell are plotted over time in each case. Rate constants obtained from (a) are shown in Fig. 6b, c, d and in Supplementary Table 3. Weighted $R^2$'s for the modeling curves are reported in Supplementary Table 4. Error bars in (a), (b) are standard errors from 80 to 150 cells in each case.

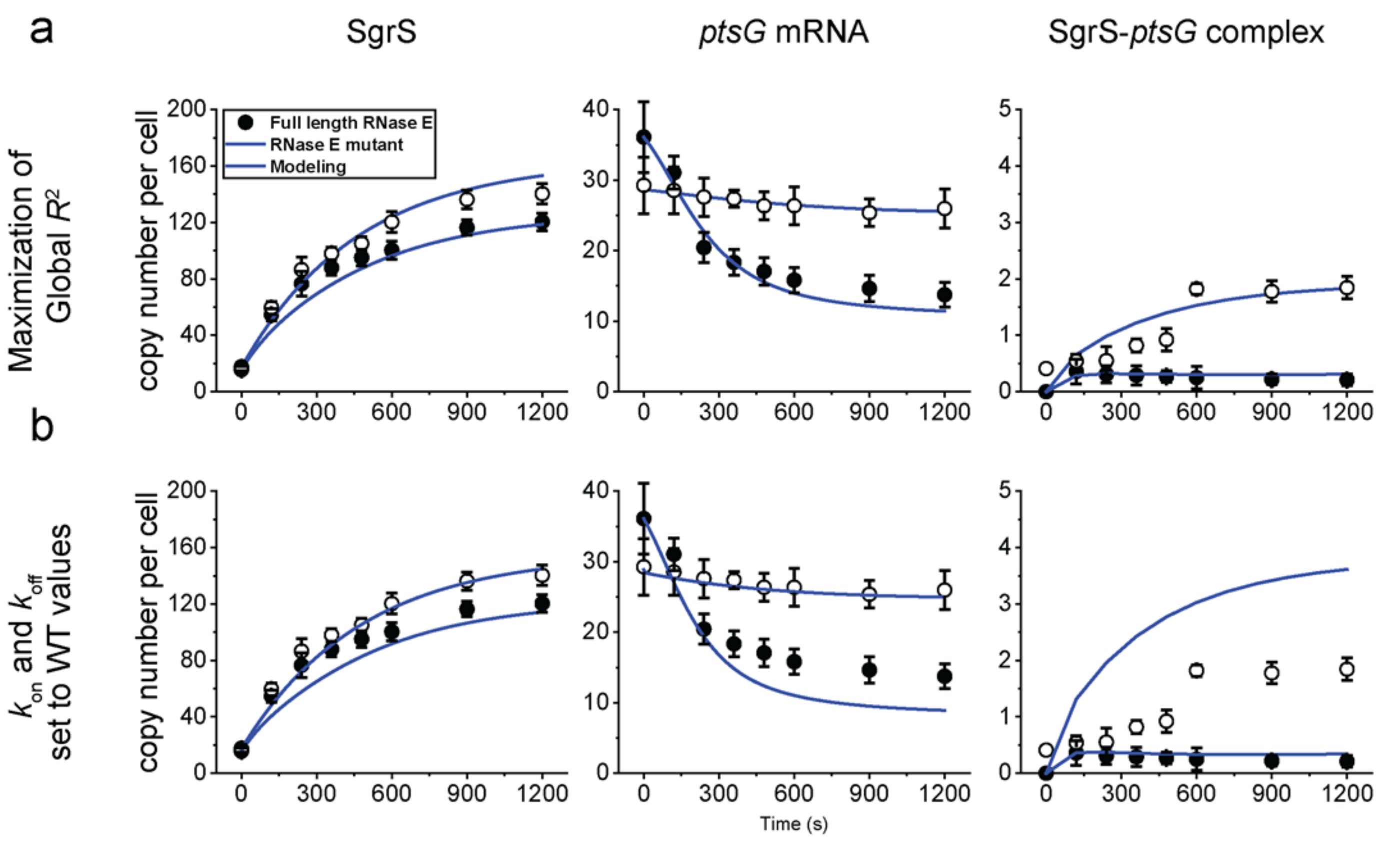


**Supplementary Figure 45. Change in the copy numbers of SgrS and *ptsG* mRNA over time for SgrS G178A mutant strain.** Time course changes from 0 min (0 s) to 20 min (1200 s) after the induction of glucose-phosphate stress using αMG (non-metabolizable sugar analog) and the corresponding modeling curves for SgrS, *ptsG* mRNA and SgrS-*ptsG* complexes for SgrS G178A strain by **(a)** maximization of global $R^2$ and **(b)** setting $k_{on}$ and $k_{off}$ to WT values. The average copy numbers per cell are plotted over time in each case. Rate constants obtained from (a) are shown in Fig. 6b, c, d and in Supplementary Table 3. Weighted $R^2$'s for the modeling curves are reported in Supplementary Table 4. Error bars in (a), (b) are standard errors from 80 to 150 cells in each case.

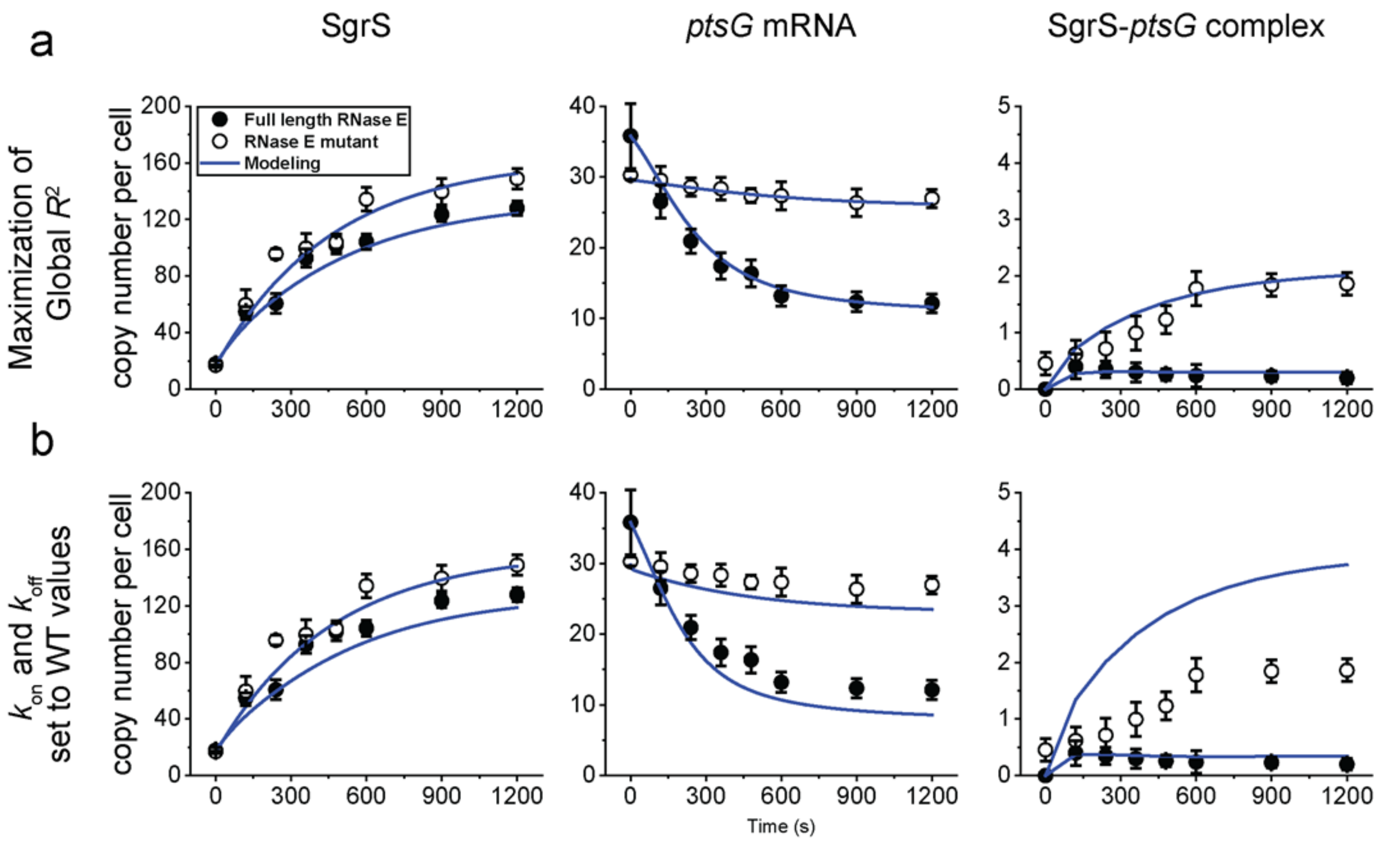


**Supplementary Figure 46. Change in the copy numbers of SgrS and *ptsG* mRNA over time for SgrS G178U mutant strain.** Time course changes from 0 min (0 s) to 20 min (1200 s) after the induction of glucose-phosphate stress using αMG (non-metabolizable sugar analog) and the corresponding modeling curves for SgrS, *ptsG* mRNA and SgrS-*ptsG* complexes for SgrS G178U strain by **(a)** maximization of global $R^2$ and **(b)** setting $k_{on}$ and $k_{off}$ to WT values. The average copy numbers per cell are plotted over time in each case. Rate constants obtained from (a) are shown in Fig. 6b, c, d and in Supplementary Table 3. Weighted $R^2$'s for the modeling curves are reported in Supplementary Table 4. Error bars in (a), (b) are standard errors from 80 to 150 cells in each case.

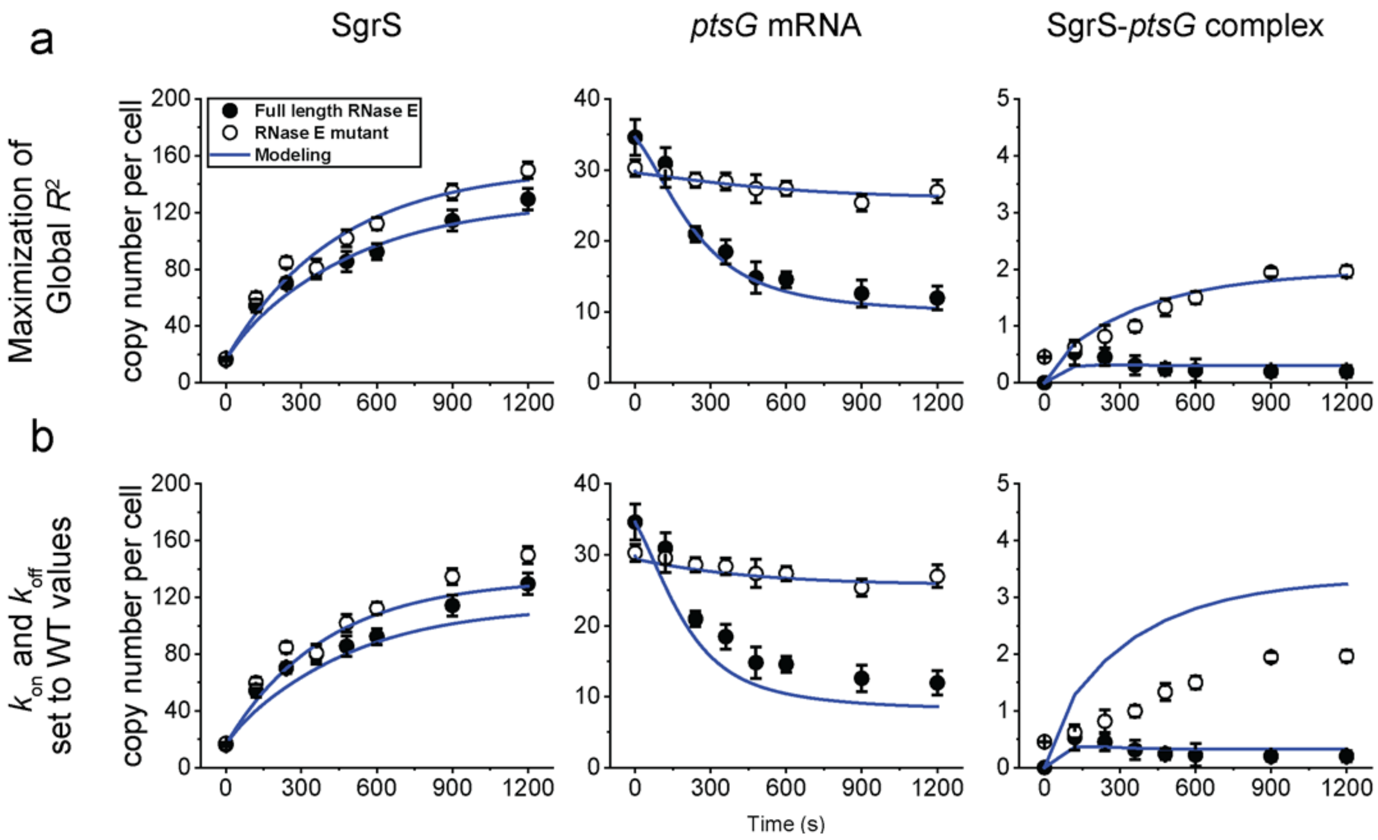


**Supplementary Figure 47. Change in the copy numbers of SgrS and *ptsG* mRNA over time for SgrS U181A mutant strain.** Time course changes from 0 min (0 s) to 20 min (1200 s) after the induction of glucose-phosphate stress using αMG (non-metabolizable sugar analog) and the corresponding modeling curves for SgrS, *ptsG* mRNA and SgrS-*ptsG* complexes for SgrS U181A strain by **(a)** maximization of global $R^2$ and **(b)** setting $k_{on}$ and $k_{off}$ to WT values. The average copy numbers per cell are plotted over time in each case. Rate constants obtained from (a) are shown in Fig. 6b, c, d and in Supplementary Table 3. Weighted $R^2$'s for the modeling curves are reported in Supplementary Table 4. Error bars in (a), (b) are standard errors from 80 to 150 cells in each case.

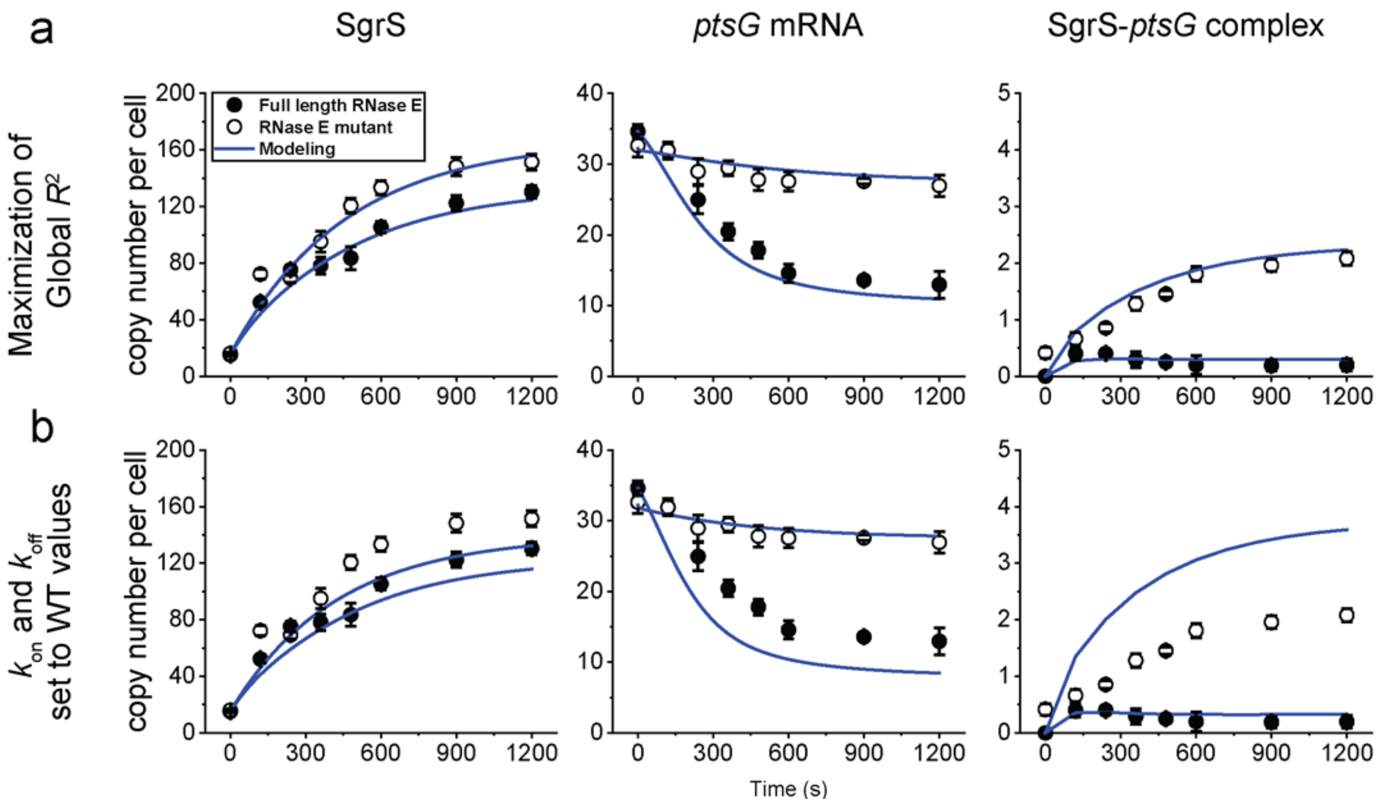


**Supplementary Figure 48. Change in the copy numbers of SgrS and *ptsG* mRNA over time for SgrS U182A mutant strain.** Time course changes from 0 min (0 s) to 20 min (1200 s) after the induction of glucose-phosphate stress using αMG (non-metabolizable sugar analog) and the corresponding modeling curves for SgrS, *ptsG* mRNA and SgrS-*ptsG* complexes for SgrS U182A strain by **(a)** maximization of global $R^2$ and **(b)** setting $k_{on}$ and $k_{off}$ to WT values. The average copy numbers per cell are plotted over time in each case. Rate constants obtained from (a) are shown in Fig. 6b, c, d and in Supplementary Table 3. Weighted $R^2$'s for the modeling curves are reported in Supplementary Table 4. Error bars in (a), (b) are standard errors from 80 to 150 cells in each case.

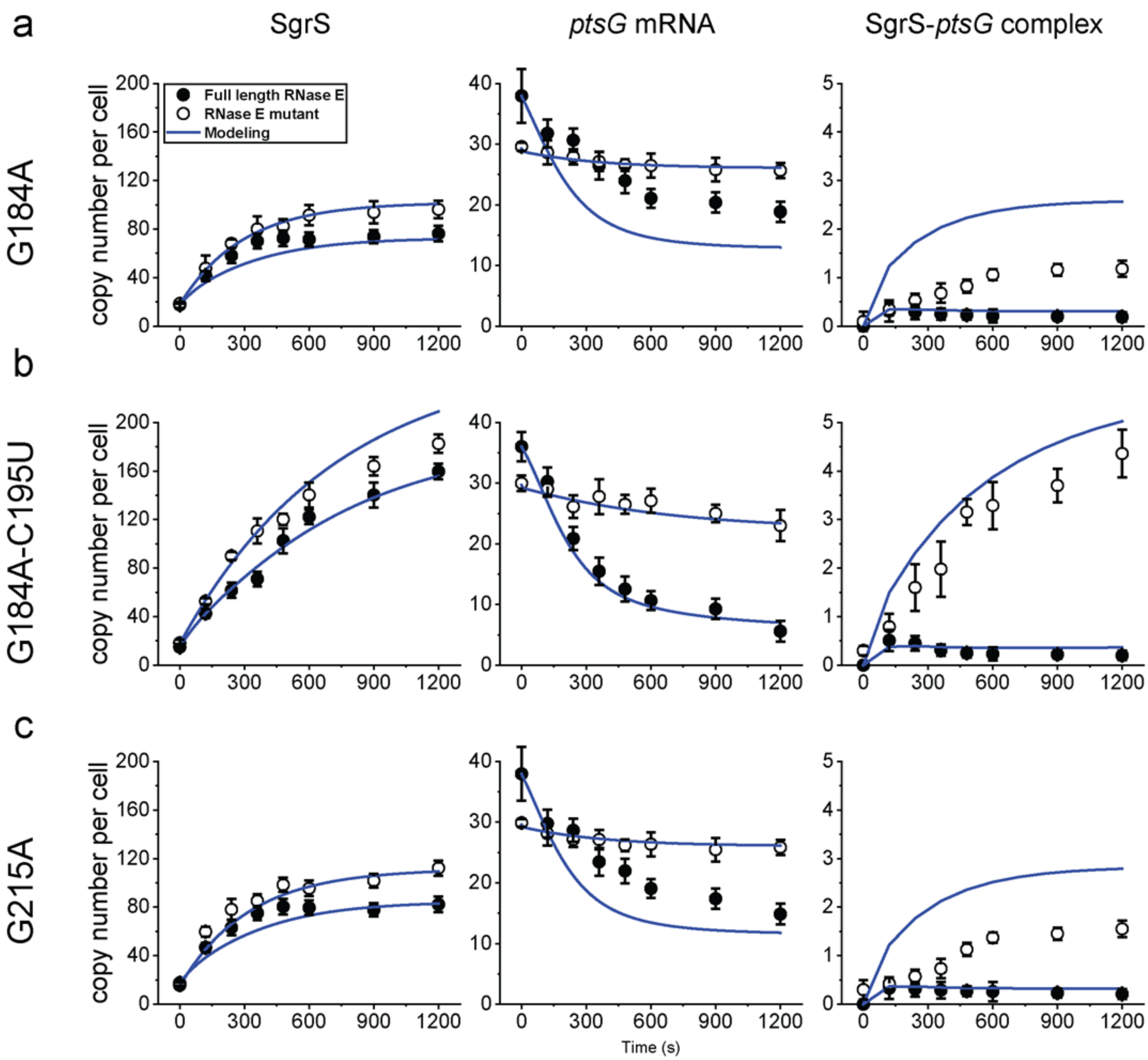


**Supplementary Figure 49. Change in the copy numbers of SgrS and *ptsG* mRNA over time for SgrS mutant strains.** Time course changes from 0 min (0 s) to 20 min (1200 s) after the induction of glucose-phosphate stress using αMG (non-metabolizable sugar analog) and the corresponding modeling curves for SgrS, *ptsG* mRNA and SgrS-*ptsG* complexes for (a) SgrS G184A strain, (b) SgrS G184A-C195U strain, (c) SgrS G215A strain by setting $k_{on}$ and $k_{off}$ to WT values. The average copy numbers per cell are plotted over time in each case. Weighted $R^2$'s for the modeling curves are reported in Supplementary Table 4. Error bars in (a), (b) are standard errors from 80 to 150 cells in each case.

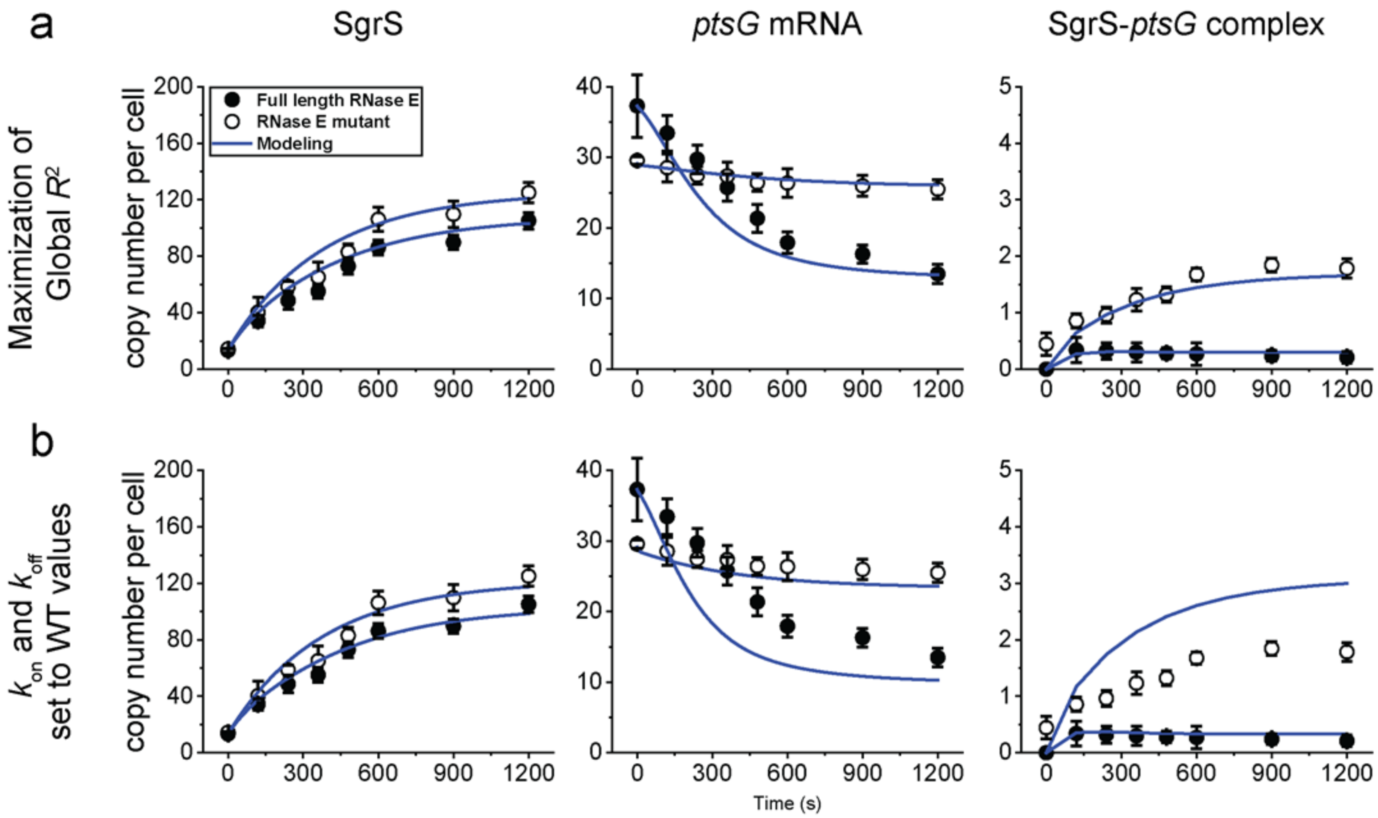


**Supplementary Figure 50. Change in the copy numbers of SgrS and *ptsG* mRNA over time for SgrS U224A mutant strain.** Time course changes from 0 min (0 s) to 20 min (1200 s) after the induction of glucose-phosphate stress using αMG (non-metabolizable sugar analog) and the corresponding modeling curves for SgrS, *ptsG* mRNA and SgrS-*ptsG* complexes for SgrS U224A strain by **(a)** maximization of global $R^2$ and **(b)** setting $k_{on}$ and $k_{off}$ to WT values. The average copy numbers per cell are plotted over time in each case. Rate constants obtained from (a) are shown in Fig. 6b, c, d and in Supplementary Table 3. Weighted $R^2$'s for the modeling curves are reported in Supplementary Table 4. Error bars in (a), (b) are standard errors from 80 to 150 cells in each case.

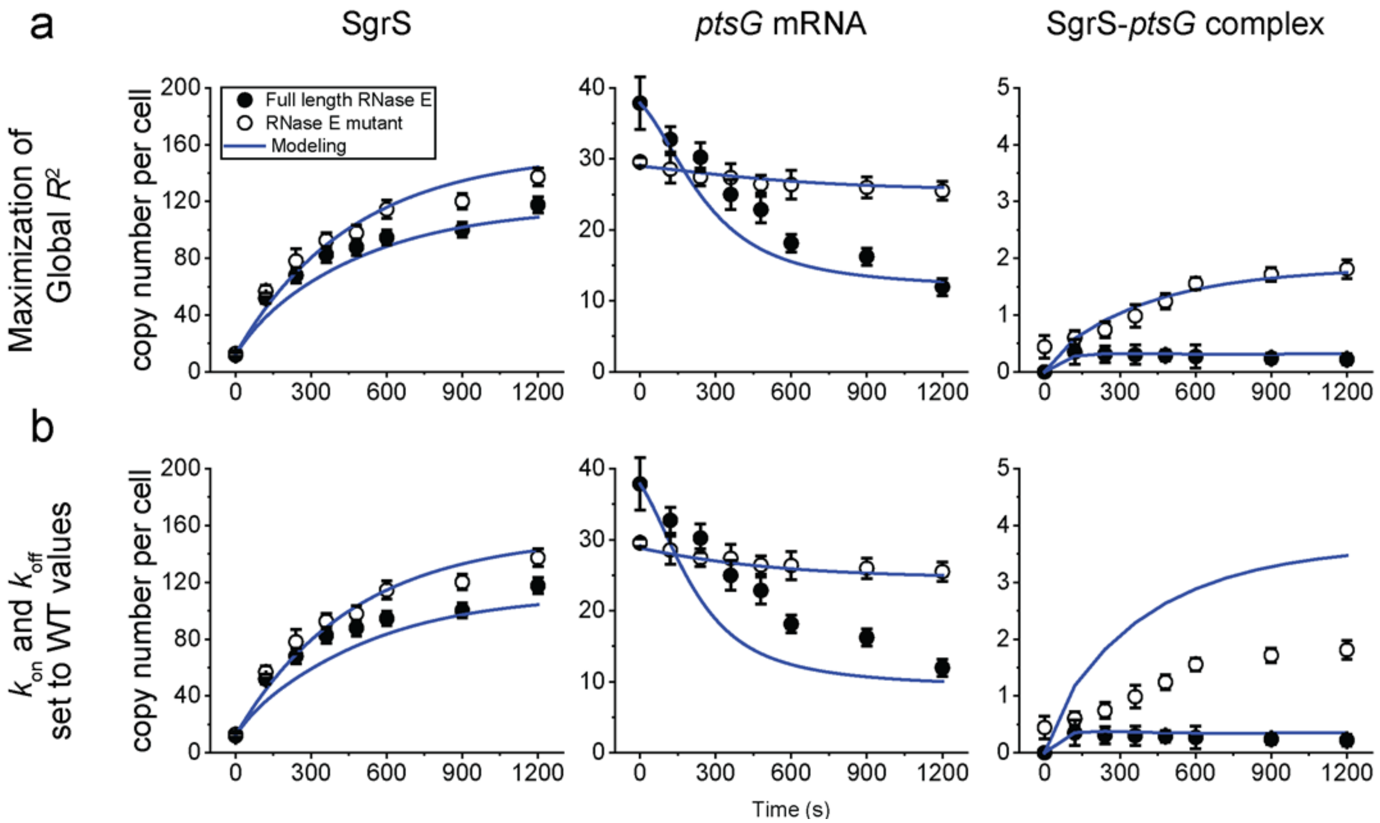


**Supplementary Figure 51. Change in the copy numbers of SgrS and *ptsG* mRNA over time for SgrS U224G mutant strain.** Time course changes from 0 min (0 s) to 20 min (1200 s) after the induction of glucose-phosphate stress using αMG (non-metabolizable sugar analog) and the corresponding modeling curves for SgrS, *ptsG* mRNA and SgrS-*ptsG* complexes for SgrS U224G strain by **(a)** maximization of global $R^2$ and **(b)** setting $k_{on}$ and $k_{off}$ to WT values. The average copy numbers per cell are plotted over time in each case. Rate constants obtained from (a) are shown in Fig. 6b, c, d and in Supplementary Table 3. Weighted $R^2$'s for the modeling curves are reported in Supplementary Table 4. Error bars in (a), (b) are standard errors from 80 to 150 cells in each case.

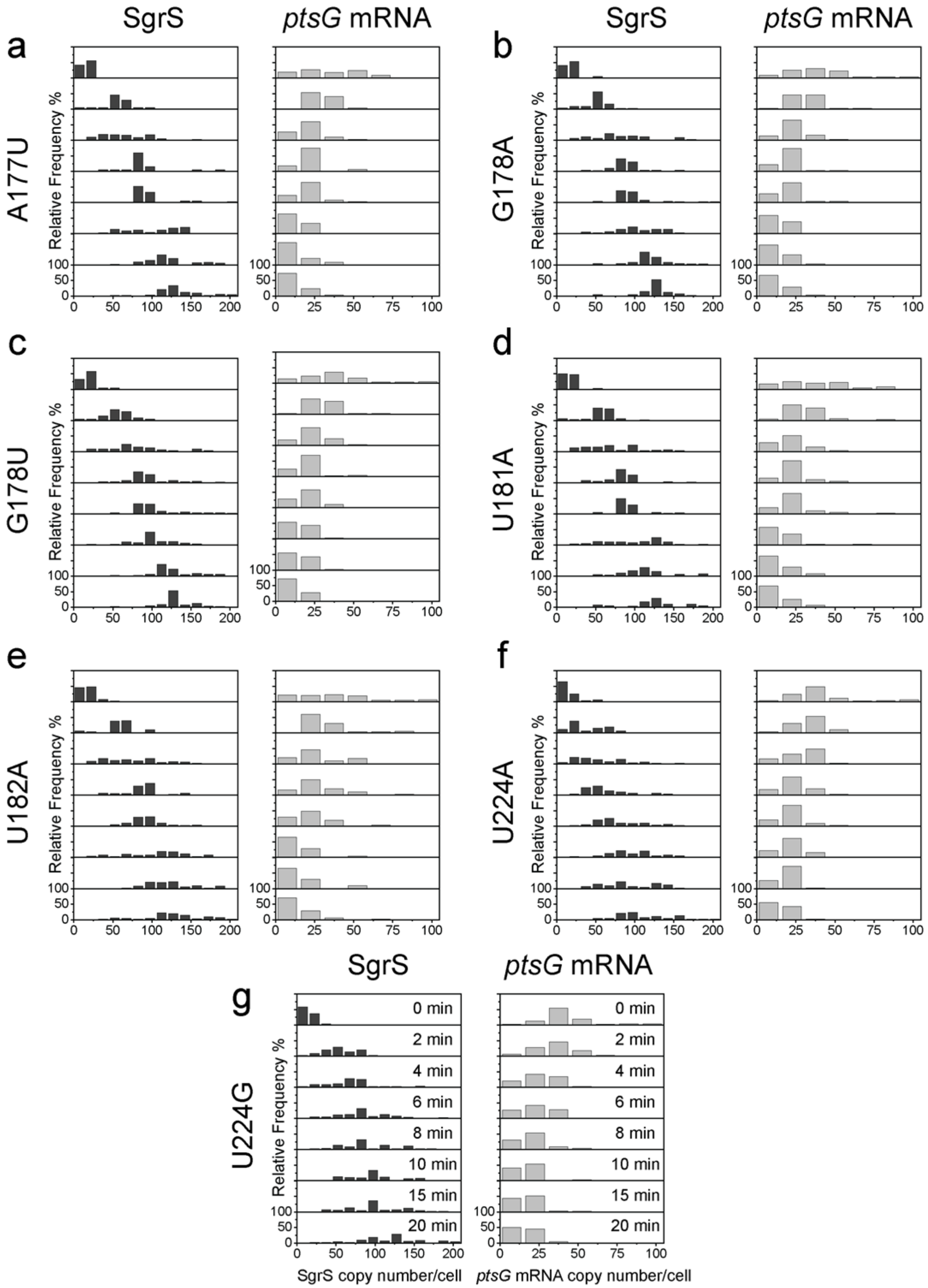


**Supplementary Figure 52. Time dependent changes in the copy numbers of SgrS and ptsG mRNA.** Histograms showing the change in distribution of SgrS and *ptsG* mRNA copy numbers for (a) SgrS A177U mutant strain, (b) G178A mutant strain, (c) G178U mutant strain, (d) U181A mutant strain, (e) U182A mutant strain, (f) U224A mutant strain, (g) U224G mutant strain for 80-150 cells in each case.

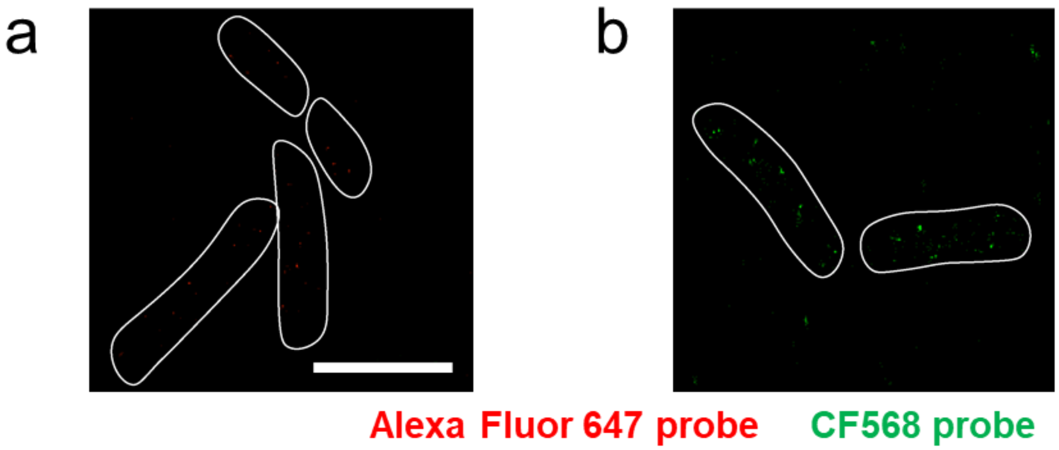


**Supplementary Figure 53. Negative control for copy number calculation. (a)** Background due to the non-specific binding of Alexa Fluor 647-labeled probes against SgrS in Δ*sgrS* strain. **(b)** Background due to the non-specific binding of CF568-labeled probes against *ptsG* mRNA in Δ*ptsG* strain. Scale bar is 2 μm.

### Supplementary Tables

### Supplementary Table 1. Plasmids and strains used in this study.

| Plasmid | Background | Source or Reference |
|---|---|---|
| pSIM6 | $P_L$ *-gam-bet-exo* genes under the control of CI857 repressor, $Amp^R$ (Ts) | Don Court |
| pZAMB1 | $P_{Ltet0\text{-}1}$-*sgrS* | [1] |
| pZEMB8 | $P_{Llac0\text{-}1}$-*ptsG-gfpsf* | [1] |
| pZAMB1A177T | $P_{Ltet0\text{-}1}$-*sgrS* A177T | This work |
| pZAMB1G178T | $P_{Ltet0\text{-}1}$-*sgrS* G178T | This work |
| pZAMB1G178A | $P_{Ltet0\text{-}1}$-*sgrS* G178A | This work |
| pZAMB1G184A | $P_{Ltet0\text{-}1}$-*sgrS* G184A | This work |
| pZAMB1C215A | $P_{Ltet0\text{-}1}$-*sgrS* C215A | This work |
| pZAMB1T224G | $P_{Ltet0\text{-}1}$-*sgrS* T224G | This work |
| pZAMB1T224A | $P_{Ltet0\text{-}1}$-*sgrS* T224A | This work |
| **Strain** | **Background** | **Source or Reference** |
| DJ480 | MG1655 Δ*lac* X74 | D. Jin, NCI |
| CV104 | DJ480 Δ*lac* X74, *mal::lacI*$^q$, Δ*sgrS::kan*$^R$ | C. K. Vanderpool, S. Gottesman, 2004 |
| DB166 | MG1655 ΔX74*lac, lacI*$^q$*-tetR-spec*$^R$ | Divya Balasubramanian |
| MB1 | Δ*ptsG,* Δ*sgrS, lacI*$^q$*-tetR-spec*$^R$ | This work |
| MB130 | lacI’::$P_{BAD}$-*ptsG'-'lacZ, λattB::lacI*$^q$*-PN25tetR-spec*$^R$*,* miniλ*tet*$^R$, Δ*araBAD araC*$^+$, *mal::lacI*$^q$ | This work |
| MB205 | *ΔsgrS::cat-sacB, lacI*$^q$*-tetR-spec*$^R$ | This work |
| MB206 | *sgrS* A177T, *lacI*$^q$*-tetR-spec*$^R$ | This work |

| | | |
|---|---|---|
| MB207 | *sgrS* G178T, *lacI*$^q$-*tetR-spec*$^R$ | This work |
| MB208 | *sgrS* G178A, *lacI*$^q$-*tetR-spec*$^R$ | This work |
| MB209 | *sgrS* G184A, *lacI*$^q$-*tetR-spec*$^R$ | This work |
| SA1701 | *sgrS* G215A, *lacI*$^q$-*tetR-spec*$^R$ | This work |
| XM180 | *sgrS* G184A-C195T, *lacI*$^q$-*tetR-spec*$^R$ | This work |
| XM181 | *sgrS* T181A, *lacI*$^q$-*tetR-spec*$^R$ | This work |
| XM182 | *sgrS* T182A, *lacI*$^q$-*tetR-spec*$^R$ | This work |
| SA1708 | *sgrS* T224A, *lacI*$^q$-*tetR-spec*$^R$ | This work |
| SA1709 | *sgrS* T224G, *lacI*$^q$-*tetR-spec*$^R$ | This work |
| TM528 | *rne701-FLAG-cat* | [2] |
| SA1740 | *sgrS* A177T, *lacI*$^q$-*tetR-spec*$^R$, *rne131::kan*$^R$ | This work |
| SA1741 | *sgrS* A178T, *lacI*$^q$-*tetR-spec*$^R$, *rne131::kan*$^R$ | This work |
| SA1742 | *sgrS* A178A, *lacI*$^q$-*tetR-spec*$^R$, *rne131::kan*$^R$ | This work |
| SA1743 | *sgrS* A184A, *lacI*$^q$-*tetR-spec*$^R$, *rne131::kan*$^R$ | This work |
| SA1744 | *sgrS* G215A, *lacI*$^q$-*tetR-spec*$^R$, *rne131::kan*$^R$ | This work |
| SA1745 | *sgrS* T224G, *lacI*$^q$-*tetR-spec*$^R$, *rne131::kan*$^R$ | This work |
| SA1746 | *sgrS* T224A, *lacI*$^q$-*tetR-spec*$^R$, *rne131::kan*$^R$ | This work |
| SA1908 | *sgrS* G184A-C195T, *lacI*$^q$-*tetR-spec*$^R$, *rne131::kan*$^R$ | This work |
| SA1909 | *sgrS* T181A, *lacI*$^q$-*tetR-spec*$^R$, *rne131::kan*$^R$ | This work |
| SA1910 | *sgrS* T182A, *lacI*$^q$-*tetR-spec*$^R$, *rne131::kan*$^R$ | This work |
| XM199 | *lacI*$^q$-*tetR-spec*$^R$, *Δhfq::kan*$^R$ | This work |
| XM200 | *sgrS* A177T, *lacI*$^q$-*tetR-spec*$^R$, *Δhfq::kan*$^R$ | This work |

| | | |
|---|---|---|
| XM201 | *sgrS* G184A, *lacI$^{q}$-tetR-spec$^{R}$, Δhfq::kan$^{R}$* | This work |
| CS196 | *ΔptsG*::*tet$^{R}$* | [3] |
| CS123 | *DJ480 strain carrying G178C/G176C mutations on sgrS* | [4,5] |

### Supplementary Table 2. Oligonucleotides used in this study.

| Oligo | Description | Sequence 5'-3' |
|---|---|---|
| 1 | Forward Primer for mutagenesis PCR of SgrS | CTATCAGTGATAGAGATACTGAGCACATATG |
| 2 | Reverse Primer for mutagenesis PCR of SgrS | GAGCCTTTCGTTTTATTTGATGGATCC |
| 3 | Forward primer for sequencing desired region of SgrS | TGGGATGACCGCAATTCTGAAA |
| 4 | Reverse primer for sequencing desired region of SgrS | GAGCCTTTCGTTTTATTTGATGGATCC |
| MBP251F | Forward primer for PCR amplification of the *cat-sacB* cassette | GCAATTTTATTTTCCCTATATTAAGTCAATAATTCCTAACAAA ATGAGACGTTGATCGGCACGTAAG |
| MBP251R | Reverse primer for PCR amplification of the *cat-sacB* cassette | TTATCCAGATCATACGTTCCCTTTTTAGCGCGGCGAGAATGTA TCAAAGGGAAACTGTCCATATGC |
| OSA766 | Forward primer for PCR amplification of | GCAATTTTATTTTCCCTATATTAAGTCAATAATTCCTAACgat GAAGCAAGGGGGTGCCC |

| | | |
|---|---|---|
| | *sgrS*, for allelic exchange between *sgrS* and the *cat-sacB* cassette | |
| MBP252R | Reverse primer for PCR amplification of *sgrS*, for allelic exchange between *sgrS* and the *cat-sacB* cassette | TTATCCAGATCATACGTTCCCTTTTTAGCGCGGCGAGAATAAA AAAAACCAGCAGGTATAATCTGCTGGCGGG |
| OSA768 | Reverse primer for PCR amplification of *sgrS* T181A, for allelic exchange between *sgrS* and the *cat-sacB* cassette | TTATCCAGATCATACGTTCCCTTTTTAGCGCGGCGAGAATaaa AAAAACCAGCAGGTATAATCTGCTGGCGGGTGATTTTACACCA TTACTCAGTCACACATGATGCAGGCA |
| OSA769 | Reverse primer for PCR amplification of *sgrS* T182A, for allelic exchange between *sgrS* and the *cat-sacB* cassette | TTATCCAGATCATACGTTCCCTTTTTAGCGCGGCGAGAATaaa AAAAACCAGCAGGTATAATCTGCTGGCGGGTGATTTTACACCT ATACTCAGTCACACATGATGCAGGC |
| MBP252R2 | Reverse primer for PCR amplification of *sgrS* G215A, for allelic exchange between *sgrS* and the *cat-sacB* cassette | TTATCCAGATCATACGTTCCCTTTTTAGCGCGGCGAGAATAAA AAAAACCAGTAGGTATAATCTGCTGGCGGG |

| | | |
|---|---|---|
| MBP252R3 | Reverse primer for PCR amplification of *sgrS* T224G, for allelic exchange between *sgrS* and the *cat-sacB* cassette | TTATCCAGATCATACGTTCCCTTTTTAGCGCGGCGAGAATAAA CAAAACCAGTAGGTATAATCTGCTGGCGGG |
| MBP252R4 | Reverse primer for PCR amplification of *sgrS* T224A, for allelic exchange between *sgrS* and the *cat-sacB* cassette | TTATCCAGATCATACGTTCCCTTTTTAGCGCGGCGAGAATAAA TAAAACCAGTAGGTATAATCTGCTGGCGGG |
| OSA499 | Primer to amplify SgrS | GATGAAGCAAGGGGGTGCCC |
| OSA500 | Primer to amplify SgrS | CAATACTCAGTCACACATGATGCAGGC |
| OXM187 | Primer to amplify *rrsA* | ATTCCGATTAACGCTTGCAC |
| OXM188 | Primer to amplify *rrsA* | AGGCCTTCGGGTTGTAAAGT |
| MBP201F | | ACCTGACGCTTTTTATCGCAACTCTCTACTGTTTCTCCATATAA ATAAAGGGCGCTTAGATGCCCTGTAC |
| MBP201R | | TAACGCCAGGTTTTCCCAGTCACGACGTTGTAAAACGACTTGC AGGTTAGCAAATGCATTCTTAAACAT |
| A177T-F | Forward primer to generate SgrS A177T mutant | CTTGCCTGCATCATGTGTGACTGTGTATTGGTGTAAAATC |
| A177T-F | Reverse primer to generate SgrS A177T mutant | GATTTTACACCAATACACAGTCACACATGATGCAGGCAA |

| | | |
|---|---|---|
| G178T-F | Forward primer to generate SgrS G178T mutant | CTTGCCTGCATCATGTGTGACTGATTATTGGTGTAAAATC |
| G178T-R | Reverse primer to generate SgrS G178T mutant | GATTTTACACCAATAATCAGTCACACATGATGCAGGCAAG |
| G178A-F | Forward primer to generate SgrS G178A mutant | CTTGCCTGCATCATGTGTGACTGAATATTGGTGTAAAATC |
| G178A-R | Reverse primer to generate SgrS G178A mutant | GATTTTACACCAATTATCAGTCACACATGATGCAGGCAAG |
| G184A-F | Forward primer to generate SgrS G184A mutant | CCTGCATCATGTGTGACTGAGTATTGATGTAAAATCACCC |
| G184A-R | Reverse primer to generate SgrS G184A mutant | GGGTGATTTTACATCAATACTCAGTCACACATGATGCAGG |
| C215A-F | Forward primer to generate SgrS C215A mutant | CCCGCCAGCAGATTATACCTACTGGTTTTTTTATTCTC |
| C215A-R | Reverse primer to generate SgrS C215A mutant | GAGAATAAAAAAACCAGTAGGTATAATCTGCTGGCGGG |
| T224G-F | Forward primer to generate SgrS T224G mutant | GATTATACCTGCTGGTTTTATTTATTCTCGCCGCG |
| T224G-R | Reverse primer to generate SgrS T224G mutant | CGCGGCGAGAATAAATAAAACCAGCAGGTATAATC |
| T224A-F | Forward primer to generate SgrS T224A mutant | GATTATACCTGCTGGTTTTGTTTATTCTCGCCGCG |
| T224A-R | Reverse primer to generate SgrS T224A mutant | CGCGGCGAGAATAAACAAAACCAGCAGGTATAATC |

| | | |
|---|---|---|
| sgrS-bio | SgrS probe used in the Northern blot analysis | GCAACCAGCACAACTTCGCTGTCGCGGTAAAATAGTG |
| ssrA-bio | 5S rRNA probe used in Northern blot analysis | CGCCACTAACAAACTAGCCTGACGCCACTAACAAA |
| SgrS | Probes* for smFISH | GTGCTGATAAAACTGACGCA<br>ACTTCGCTGTCGCGGTAAAA<br>CTTAACCAACGCAACCAGCA<br>CATGGTTAATCGTTGTGGGA<br>ATCCCACTGCATCAGTCCTT<br>GTCAACTTTCAGAATTGCGG<br>TCAGTCACACATGATGCAGG<br>GCGGGTGATTTTACACCAAT<br>AACCAGCAGGTATAATCTGC |
| *ptsG* | Probes* for smFISH | GCATCTAAGCGCCCTTTATT<br>GCAGGTTAGCAAATGCATTC<br>ATCAGCGATTTACCGACCTT<br>CTGCCATAACATGCGATACA<br>CATGTTTGCAAAGACGGAAC<br>GACACCGATCGCAAAAATCA<br>GATACGCCATCGTTATTGGT<br>ATGATGCCATAGGCAACAAC<br>AACCACGGCCATGGTTTTAA<br>CAGGTGTTTAGAGGCGATTT<br>TAAACATGTACGCTGCGATC<br>GGCAGCTTAATACGGTAGAA<br>GGCAAAGAAGCCAAGATACT<br>CAGAAATGATCGGCACAAAG<br>CCAAATGAAGGACAGCACAA<br>ACTGAGAGAAGGTCTGGATT<br>CAACGTTCGATGAAACCGTA<br>GGTGTATTCACCAATCTGCA<br>GGCAGACCGTACATTTTGAA<br>CGGTTTTCTGGTTTAGCAGA<br>AACGATGAAGTCGATCAGAC<br>GCGGAAGATGGTGTAGTAAA |

| | | |
|---|---|---|
| | | CGTTTTCAGATCCAGTGCTT<br>TCGCTTTTGCATCTTCAGTC<br>TCTTTACCACCAAATGCAGC<br>TACATGCGTCGAGGTTAGTA<br>TTAGTACCGAAAATCGCCTG<br>AGTGGTTACGGATGTACTCA |

*The probes were complementary sequences across the length of the RNAs.

**Supplementary Table 3. Rate constants for SgrS, *ptsG* mRNA association, dissociation and SgrS-*ptsG* complex codegradation.**

| | $\alpha_s$ (molecules-s$^{-1}$) | $\beta_s$ (s$^{-1}$) | $\alpha_p$ (molecules-s$^{-1}$) | $\beta_p$ (s$^{-1}$) | $k_{on}$ (M$^{-1}$-s$^{-1}$) x $10^5$ | $k_{off}$ (s$^{-1}$) | $k_{cat}$ (s$^{-1}$) |
|---|---|---|---|---|---|---|---|
| **Wild Type** | 0.35 ± 0.01 | 0.0013 ± 0.0001 | 0.13 ± 0.02 | 0.0037 ± 0.0005 | 1.9 ± 0.2 | 0.22 ± 0.02 | 0.3 ± 0.1 |
| **Wild Type RNase E Mutant** | 0.35 ± 0.01 | 0.0013 ± 0.0001 | 0.11 ± 0.02 | 0.0037 ± 0.0005 | 1.9 ± 0.2 | 0.22 ± 0.02 | 0.0062 ± 0.0010 |
| **A177U** | 0.36 ± 0.02 | 0.0021 ± 0.0002 | 0.14 ± 0.02 | 0.0039 ± 0.0006 | 1.5 ± 0.1 | 0.25 ± 0.01 | 0.31 ± 0.09 |
| **A177U RNase E Mutant** | 0.36 ± 0.02 | 0.0021 ± 0.0002 | 0.12 ± 0.02 | 0.0039 ± 0.0006 | 1.5 ± 0.1 | 0.25 ± 0.01 | 0.0060 ± 0.0021 |
| **G178A** | 0.36 ± 0.02 | 0.0021 ± 0.0003 | 0.14 ± 0.03 | 0.0038 ± 0.0008 | 1.3 ± 0.1 | 0.27 ± 0.01 | 0.3 ± 0.1 |
| **G178A RNase E Mutant** | 0.36 ± 0.02 | 0.0021 ± 0.0003 | 0.11 ± 0.03 | 0.0038 ± 0.0008 | 1.3 ± 0.1 | 0.27 ± 0.01 | 0.0078 ± 0.0023 |
| **G178U** | 0.36 ± 0.02 | 0.0021 ± 0.0003 | 0.14 ± 0.03 | 0.0039 ± 0.0007 | 1.3 ± 0.1 | 0.27 ± 0.01 | 0.32 ± 0.07 |
| **G178U RNase E Mutant** | 0.36 ± 0.02 | 0.0021 ± 0.0003 | 0.12 ± 0.02 | 0.0039 ± 0.0007 | 1.3 ± 0.1 | 0.27 ± 0.01 | 0.0075 ± 0.0020 |
| **U181A** | 0.36 ± 0.02 | 0.0022 ± 0.0003 | 0.13 ± 0.02 | 0.0037 ± 0.0005 | 1.4 ± 0.1 | 0.26 ± 0.01 | 0.34 ± 0.09 |
| **U181A RNase E Mutant** | 0.36 ± 0.02 | 0.0022 ± 0.0003 | 0.11 ± 0.02 | 0.0037 ± 0.0005 | 1.4 ± 0.1 | 0.26 ± 0.01 | 0.0065 ± 0.0010 |
| **U182A** | 0.34 ± 0.02 | 0.0023 ± 0.0003 | 0.14 ± 0.02 | 0.0040 ± 0.0006 | 1.4 ± 0.2 | 0.265 ± 0.01 | 0.3 ± 0.1 |
| **U182A RNase E Mutant** | 0.34 ± 0.02 | 0.0023 ± 0.0003 | 0.13 ± 0.02 | 0.0040 ± 0.0006 | 1.4 ± 0.2 | 0.265 ± 0.01 | 0.0060 ± 0.0020 |

| | | | | | | | |
|---|---|---|---|---|---|---|---|
| **G184A** | 0.35 ± 0.02 | 0.0034 ± 0.0003 | 0.15 ± 0.03 | 0.0040 ± 0.0007 | 0.95 ± 0.1 | 0.293 ± 0.002 | 0.32 ± 0.1 |
| **G184A RNase E Mutant** | 0.35 ± 0.02 | 0.0034 ± 0.0003 | 0.12 ± 0.02 | 0.0040 ± 0.0007 | 0.95 ± 0.1 | 0.293 ± 0.002 | 0.0080 ± 0.0028 |
| **G184A-C195U** | 0.35 ± 0.02 | 0.0014 ± 0.0002 | 0.14 ± 0.03 | 0.0039 ± 0.0008 | 1.85 ± 0.18 | 0.225 ± 0.015 | 0.33 ± 0.08 |
| **G184A-C195U RNase E Mutant** | 0.35 ± 0.02 | 0.0014 ± 0.0002 | 0.12 ± 0.02 | 0.0039 ± 0.0008 | 1.85 ± 0.18 | 0.225 ± 0.015 | 0.0068 ± 0.0020 |
| **G215A** | 0.36 ± 0.02 | 0.0025 ± 0.0003 | 0.15 ± 0.02 | 0.0040 ± 0.0006 | 1.2 ± 0.1 | 0.28 ± 0.01 | 0.30 ± 0.07 |
| **G215A RNase E Mutant** | 0.36 ± 0.02 | 0.0025 ± 0.0003 | 0.12 ± 0.02 | 0.0040 ± 0.0006 | 1.2 ± 0.1 | 0.28 ± 0.01 | 0.0082 ± 0.0030 |
| **U224A** | 0.34 ± 0.01 | 0.0024 ± 0.0003 | 0.14 ± 0.03 | 0.0038 ± 0.0007 | 1.28 ± 0.03 | 0.258 ± 0.02 | 0.33 ± 0.08 |
| **U224A RNase E Mutant** | 0.34 ± 0.01 | 0.0024 ± 0.0003 | 0.11 ± 0.02 | 0.0038 ± 0.0007 | 1.28 ± 0.03 | 0.258 ± 0.02 | 0.0080 ± 0.0018 |
| **U224G** | 0.35 ± 0.01 | 0.0022 ± 0.0003 | 0.14 ± 0.02 | 0.0038 ± 0.0005 | 1.3 ± 0.1 | 0.27 ± 0.01 | 0.32 ± 0.09 |
| **U224G RNase E Mutant** | 0.35 ± 0.01 | 0.0022 ± 0.0003 | 0.11 ± 0.02 | 0.0038 ± 0.0005 | 1.3 ± 0.1 | 0.27 ± 0.01 | 0.0075 ± 0.0020 |

**Supplementary Table 4. Goodness of fit for the time dependent curves.**

| | Global $R^2$ | $R^2$ for SgrS | $R^2$ for ptsG | $R^2$ for complex | $\chi^2$ for complex | $\alpha$ for complex |
|---|---|---|---|---|---|---|
| **Wild-type** | 0.999 | 0.999 | 0.995 | - | 0.37 | 0.01 |
| **Wild-type RNase E Mutant** | 0.999 | 0.999 | 0.963 | 0.902 | - | - |
| **A177U** | 0.998 | 0.998 | 0.993 | - | 0.33 | 0.001 |
| **A177U with WT $k_{on}$ and $k_{off}$** | 0.990 | 0.996 | 0.970 | - | 0.27 | 0.001 |
| **A177U RNase E Mutant** | 0.998 | 0.999 | 0.927 | 0.805 | - | - |
| **A177U RNase E Mutant with WT $k_{on}$ and $k_{off}$** | 0.990 | 0.998 | 0.796 | 0.581 | - | - |
| **G178A** | 0.999 | 0.999 | 0.990 | - | 0.09 | 0.001 |
| **G178A with WT $k_{on}$ and $k_{off}$** | 0.990 | 0.993 | 0.931 | - | 0.15 | 0.001 |
| **G178A RNase E Mutant** | 0.999 | 0.999 | 0.962 | 0.700 | - | - |
| **G178A RNase E Mutant with WT $k_{on}$ and $k_{off}$** | 0.990 | 0.998 | 0.749 | - | 17.73 | 0.99 |
| **G178U** | 0.999 | 0.999 | 0.994 | - | 0.13 | 0.001 |
| **G178U with WT $k_{on}$ and $k_{off}$** | 0.991 | 0.994 | 0.959 | - | 0.15 | 0.001 |
| **G178U RNase E Mutant** | 0.999 | 0.999 | 0.970 | 0.795 | - | - |
| **G178U RNase E Mutant with WT $k_{on}$ and $k_{off}$** | 0.991 | 0.999 | 0.772 | - | 21.36 | 0.999 |
| **U181A** | 0.998 | 0.998 | 0.997 | - | 0.27 | 0.001 |
| **U181A with WT $k_{on}$ and $k_{off}$** | 0.982 | 0.982 | 0.979 | - | 2.63 | 0.15 |
| **U181A RNase E Mutant** | 0.998 | 0.998 | 0.989 | 0.789 | - | - |
| **U181A RNase E Mutant with WT $k_{on}$ and $k_{off}$** | 0.982 | 0.988 | 0.978 | - | 24.59 | 0.9995 |

| | | | | | | |
|---|---|---|---|---|---|---|
| **U182A** | 0.997 | 0.998 | 0.986 | - | 0.69 | 0.01 |
| **U182A with WT $k_{on}$ and $k_{off}$** | 0.987 | 0.995 | 0.968 | - | 2.57 | 0.15 |
| **U182A RNase E Mutant** | 0.997 | 0.997 | 0.990 | 0.854 | - | - |
| **U182A RNase E Mutant with WT $k_{on}$ and $k_{off}$** | 0.987 | 0.994 | 0.975 | - | 17.26 | 0.99 |
| **G184A** | 0.998 | 0.998 | 0.996 | - | 0.12 | 0.001 |
| **G184A with WT $k_{on}$ and $k_{off}$** | 0.982 | 0.987 | 0.837 | - | 0.18 | 0.001 |
| **G184A RNase E Mutant** | 0.998 | 0.999 | 0.997 | 0.704 | - | - |
| **G184A RNase E Mutant with WT $k_{on}$ and $k_{off}$** | 0.982 | 0.999 | 0.969 | - | 5.47 | 0.5 |
| **G184A-C195U** | 0.999 | 0.999 | 0.976 | - | 0.28 | 0.001 |
| **G184A-C195U with WT $k_{on}$ and $k_{off}$** | 0.996 | 0.997 | 0.984 | - | 0.29 | 0.001 |
| **G184A-C195U RNase E Mutant** | 0.999 | 0.999 | 0.957 | 0.953 | - | - |
| **G184A-C195U RNase E Mutant with WT $k_{on}$ and $k_{off}$** | 0.996 | 0.997 | 0.975 | 0.905 | - | - |
| **G215A** | 0.998 | 0.998 | 0.992 | - | 0.04 | 0.001 |
| **G215A with WT $k_{on}$ and $k_{off}$** | 0.988 | 0.990 | 0.913 | - | 0.11 | 0.001 |
| **G215A RNase E Mutant** | 0.998 | 0.998 | 0.836 | 0.712 | - | - |
| **G215A RNase E Mutant with WT $k_{on}$ and $k_{off}$** | 0.988 | 0.998 | 0.928 | - | 4.56 | 0.4 |
| **U224A** | 0.998 | 0.998 | 0.982 | - | 0.06 | 0.001 |
| **U224A with WT $k_{on}$ and $k_{off}$** | 0.989 | 0.999 | 0.913 | - | 0.12 | 0.001 |
| **U224A RNase E Mutant** | 0.998 | 0.998 | 0.912 | 0.702 | - | - |
| **U224A RNase E Mutant with WT $k_{on}$ and $k_{off}$** | 0.989 | 0.997 | 0.645 | - | 37.39 | 0.999995 |
| **U224G** | 0.998 | 0.998 | 0.964 | - | 0.08 | 0.001 |

| | | | | | | |
|---|---|---|---|---|---|---|
| **U224G with WT $k_{on}$ and $k_{off}$** | 0.988 | 0.991 | 0.888 | - | 0.13 | 0.001 |
| **U224G RNase E Mutant** | 0.998 | 0.998 | 0.933 | 0.941 | - | - |
| **U224G RNase E Mutant with WT $k_{on}$ and $k_{off}$** | 0.988 | 0.997 | 0.981 | - | 5.731 | 0.55 |

### Supplementary Text

#### Impact of SgrS mutations on *ptsG* mRNA intrinsic degradation rates

In order to solve the deterministic model (equations shown in Fig. 1), we also needed to obtain the SgrS-independent degradation rates of *ptsG* mRNA. To determine this in various SgrS mutant strains, we grew cells without SgrS induction and added rifampicin to stop transcription. Samples were collected over time and fixed using paraformaldehyde. The fixed cells were then subjected to the same imaging and analysis protocol and the average copy number of *ptsG* mRNA per cell was calculated as a function of time after transcription inhibition. The degradation rate thus determined was 0.0037 ± 0.0005 $s^{-1}$ (4.5 ± 0.6 min lifetime) for the wild type strain and remained similar for all the mutants (Fig. S24-S34, S39). This verified that the mutations in the SgrS by themselves do not affect the *ptsG* mRNA stability in the absence of sugar stress and co-degradation.
